# Bayesian data driven modelling of kinetochore dynamics: space-time organisation of the human metaphase plate

**DOI:** 10.1101/2025.01.22.634311

**Authors:** Constandina Koki, Alessio V. Inchingolo, Abdullahi Daniyan, Enyu Li, Andrew D. McAinsh, Nigel J. Burroughs

## Abstract

Mitosis is a complex self-organising process that achieves high fidelity separation of duplicated chromosomes into two daughter cells through capture and alignment of chromosomes to the spindle mid-plane. Chromosome movements are driven by kinetochores, multi-protein machines that attach chromosomes to microtubules (MTs), both controlling and generating directional forces. Using lattice light sheet microscopy imaging and automated near-complete tracking of kinetochores at fine spatio-temporal resolution, we produce a detailed atlas of kinetochore metaphase-anaphase dynamics in untransformed human cells (RPE1). We determined the support from this dataset for 17 models of metaphase dynamics using Bayesian inference, demonstrating (1) substantial sister asymmetry that transversely organises the metaphase plate (MPP), (2) substantial spatial organisation of KT dynamic properties within the MPP, and (3) mechanical parameter time dependence, K-fiber forces tuning over the last 5 mins of metaphase towards a set point referred to as the anaphase ready state. These spatio-temporal trends are robust to spindle assembly pathways that are error-prone, suggesting the underpinning processes of kinetochore heterogeneity are intrinsic to mitosis and possibly by design.

## 1 Introduction

Chromosome segregation relies on the self-assembly of a microtubule-based dynamic bipolar spindle. The microtubules (MTs) are dynamic, undergoing cycles of polymerisation and depolymerisation (dynamic instability), [Mitchison and Kirschner, 1984]. They are nucleated at the centrosomes (spindle pole), and extend radially with a subset forming discrete bundles of ∼ 10 microtubules, the Kinetochore (K)-Fibres, Kiewisz et al. [2022], that connect to each of the 92 sister chromatids (replicated chromosomes). These connections are mediated by kinetochores (KTs), multi-protein machines that are able to maintain attachment to the plus-ends of microtubule bundles as they grow and shrink thereby generating pushing and pulling forces, respectively. Each microtubule-KT attachment independently generates, and also brokers, these forces to orchestrate chromosome movements and, ultimately segregation of sister chromatids into daughter cells during anaphase, [Rago and Cheeseman, 2013, McAinsh and Marston, 2022].

Each of the 46 sister KT pairs must become bi-orientated, *i*.*e*., sister chromatids are attached to K-fibers emanating from opposite poles of the spindle – this is the only geometry compatible with accurate chromosome segregation. Concurrently with biorientation, chromosomes align at the spindle equatorial plane (termed congression) forming the metaphase plate (MPP). During metaphase, which lasts ∼10 mins, chromosomes undergo quasi-periodic oscillations along the spindle axis, [Skibbens et al., 1993, Wan et al., 2012]. These oscillations are largely driven by sister KTs switching between poleward (P; attached microtubules depolymerising) and away-from-the-pole (AP; attached microtubules polymerising) moving states. When one sister is P and the other AP the sister chromatids undergo sustained directional movement. Sister chromatids are connected by centromeric chromatin which operates as a spring, [Jaqaman et al., 2010]. Directional switches arise when both sister KTs switch directional state (P, AP), that is believed to be regulated by the centromeric spring tension, [Wan et al., 2012, Burroughs et al., 2015]. Metaphase oscillations thus provide a unique opportunity to examine the mechanisms by which KTs generate and sense forces.

The purpose of metaphase remains a mystery. One possibility is that it simply reflects a “ waiting” state before anaphase onset, the duration being linked to the rates of biochemical events necessary to initiate chromosome segregation in anaphase *i*.*e*., Cyclin and Securin destruction that starts after checkpoint satisfaction, although mitotic slippage can occur, [Dick and Gerlich, 2013]. However, in asymmetric centriole distributed spindle poles there is a SAC dependent anaphase delay that enables the MPP to centralise [Tan et al., 2015], suggesting a role in quality control. There is also evidence of mechanical changes during this time suggesting active processes are present that modify the dynamics. This includes a thinning of the MPP (reduction in width along spindle axis) that is also observed in PTK1 cells, [Cimini et al., 2001], thought to be related to a decrease in kinetochore speed, [Jaqaman et al., 2010], a decrease in KT swivel (increased alignment of the intra-kinetochore axis with the sister-sister axis) reflecting a decrease in the torque acting on KTs, [Smith et al., 2016] and a synchronisation of tension across sister pairs, [Matos et al., 2009]. We refer to this body of changes as *metaphase maturation*. Of note is that metaphase oscillations in cancer cells are attenuated, [Iemura et al., 2021a,b], and that centromeric spring maturation from prometaphase to metaphase is disrupted in aneuploid cell lines, [Harasymiw et al., 2019].

However, a cell level or mechanistic understanding of the time evolution of forces in the mitotic spindle during metaphase and anaphase is missing. One challenge is that mitotic events occur over multiple time scales, specifically there is i) fast directional switching with both sisters switching direction on the timescale of seconds, [Burroughs et al., 2015], ii) quasi-periodic metaphase oscillations with a period of a minute, iii) slow maturation of the MPP on the timescale of minutes (reduction in MPP width, increased sister-sister axis alignment), iv) the spindle turns over on a ∼5 minute timescale (MT retrograde flux is 0.8-1.5 *µ*m/min, Risteski et al. [2022] half-spindle size is approximately 6 *µ*m in RPE1 cells), and v) mitosis occurs over a duration of 1 hour (nuclear envelope breakdown (NEBD) to anaphase onset is around 25 mins). A second challenge is that although chromosomes are typically treated as identical objects, there is a high degree of heterogeneity. Specifically, metaphase kinetochore oscillation quality varies substantially within a cell, including a fraction of non-oscillating pairs, whilst the position of the chromosome within the 3D spindle has been reported to influence mechanical forces with both Polar Ejection Forces (PEF), [Armond et al., 2015a, Civelekoglu-Scholey et al., 2013b] and KT swivel, [Smith et al., 2016], increasing towards the periphery of the metaphase plate. Non-sister kinetochores may also influence each others’ behaviour, their motion being correlated to that of neighbouring KTs, [Vladimirou et al., 2013], hypothesised to be due to cross-linking between K-fibres, [Vladimirou et al., 2013, Elting et al., 2017]. The biophysical properties and dynamics of KTs are also expected to vary simply because of stochasticity inherent in the spindle self-assembly process; this would include outer KT assembly in prophase and the formation of K-fibers following nuclear breakdown.

Understanding this complex multi-scale mechanical system requires development of quantitative mathematical models that can capture crucial elements of the system’s biophysics and regulatory properties, which can then provide quantitative support for conceptual ideas and generate testable predictions. Efforts in this direction have been ongoing since the 1980’s Vladimirou et al. [2011], Civelekoglu-Scholey and Cimini [2014] with previous work focusing on microscopic models of kinetochore-microtubule attachment, [Hill, 1985, Joglekar and Hunt, 2002], and a range of dynamic processes, including (but not limited to) the origin of metaphase oscillations, Civelekoglu-Scholey et al. [2006, Civelekoglu-Scholey et al. 2013b], Banigan et al. [2015], Schwietert and Kierfeld [2020], Medina et al. [2021], the role of bridging fibres and spindle geometry [Kajtez et al., 2016, Miles et al., 2022], chromosome congression dynamics to the spindle equator, [Holy and Leibler, 1994, Liu et al., 2008, Mogilner et al., 2006, Loughlin et al., 2010, Zaytsev and Grishchuk, 2015, Blackwell et al., 2017, Kliuchnikov et al., 2023, Risteski et al., 2022], formation and stability of bipolar spindles Chatterjee et al. [2020], Li et al. [2023], and the anaphase transition Gay et al. [2012]. Careful calibration of models to experimental data is crucial to ensure model validity. However, model parameters cannot typically be directly estimated from data because of its high dynamic complexity and mechanical coupling; direct measures such as KT speed and oscillation period have a nonlinear dependence on multiple mechanical parameters and thus present a complex inverse problem. Therefore, few studies have inferred model parameters directly from experimental data. In a previous work, Armond et al. [2015a] fitted a biophysical model of metaphase oscillations to 3D kinetochore tracking data from HeLa cells, a transformed human cancer cell line which features extensive chromosome instability. The fitted model provided fundamental insight into the forces acting on kinetochores and how sister kinetochores coordinated directional switching, [Armond et al., 2015a, Burroughs et al., 2015]. A similar analysis on non-cancer cells has not been carried out, thus given the known dynamic perturbations in cancer cells, [Iemura et al., 2021a, Harasymiw et al., 2019], there is a fundamental gap in our knowledge.

In this work, we generalise the paired sister kinetochore mechanical model of Armond et al. [2015a] to incorporate sister asymmetry (so that sister KTs/K-fibers are not dynamically identical), time dependence in model parameters (thereby capturing dynamic maturation), and extend the model from metaphase to anaphase. Using Bayesian inference, specifically through Markov Chain Monte Carlo (MCMC) algorithms, we parametrised our biophysical models from experimental sister kinetochore trajectory data acquired from living human cells with lattice light sheet microscopy; all model parameters were inferrable. We then used model selection methods (Bayes factor assessment of model preference, Kass and Raftery [1995b]), to determine which models are supported (preferred) by the data. This data-drive approach has provided key insights into how sister kinetochore dynamics are defined by both spatial and temporal cues.

## 2 Results

### 2.1 Near-complete kinetochore tracking through the metaphase-anaphase transition

To obtain insight into chromosome dynamics in metaphase and at the anaphase transition, we developed a tracking algorithm (KiT v3.0, Daniyan et al. [2024], based on earlier versions Harrison et al. [2022]), that achieves near-complete tracking of fluorescently-labelled KTs in human RPE1 cells [Roscioli et al., 2020]. The tracking pipeline consists of: deconvolving the 4D movies; detecting candidate spots with a Constant False Alarm Rate (CFAR) based spot detection algorithm, [Daniyan et al., 2024]; refining spot locations using a Gaussian mixture model to provide subpixel resolution; fitting a plane to the KT population thus defining the metaphase plate and an associated reference coordinate system; linking detected particles betwethe sister mid-points en frames over time to form tracks; and pairing sister KTs based on their metaphase dynamics. This provides sub-pixel resolution for the positions of each KT, and allows us to study dynamics of sister KT pairs, rather than simply individual KTs.

We performed live-cell imaging of RPE1 cells expressing endogenous NDC80-GFP (a kinetochore marker; [Roscioli et al., 2020]) using lattice light sheet microscopy (Figure 1). Data were collected at a high temporal resolution of 2.05s per *z*-stack over long timescales, typically tens of minutes, starting during metaphase through to anaphase. A typical cell is shown in Figure 1, where we detected an average of 90 spots over 350 frames (724.5 secs), with 82 KTs tracked throughout the movie and 43 KT pairs tracked for at least 75% of the movie (38 KT tracks remaining paired for the entire duration of metaphase, see Appendix C1, Appendix Figure B1). This is close to the expected 92 KTs (46 paired chromatids) for a human cell line with a diploid 46,XY karyotype. The tracks along the metaphase plate normal are shown in Figure 1E. We imaged 36 cells. We required good sister pair coverage per cell, specifically at least 30 sister pairs both tracked for 75% of movie; 31 cells satisfied this criteria giving 1281 tracked sister pairs. On average, we obtained 40 sister pairs per cell (quartiles Q1=38.5, Q3=43), where both sisters were tracked for at least 200 seconds (100 frames).

**Figure 1.**
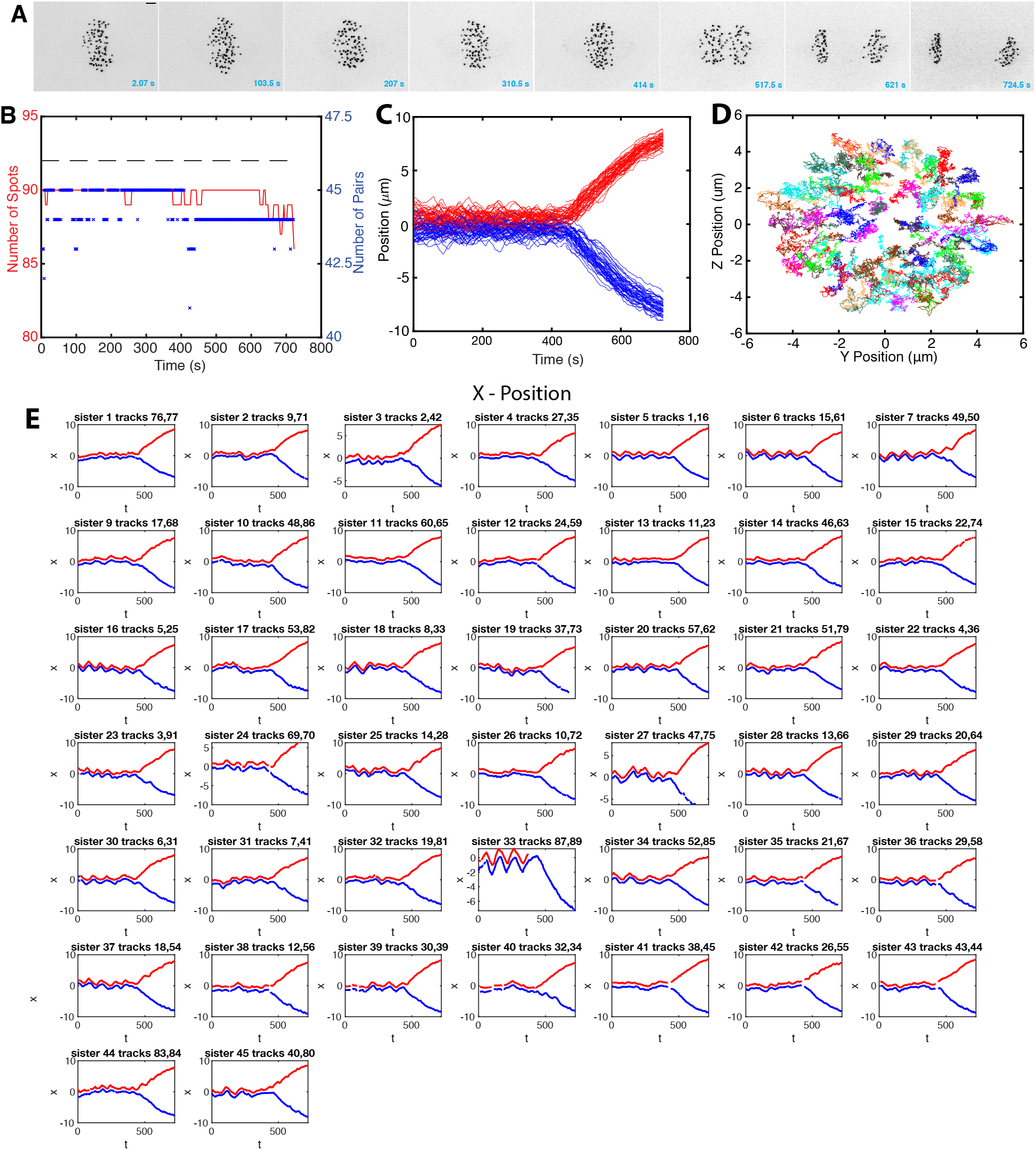
Near-complete tracking of kinetochores through metaphase and anaphase in human RPE1 cells. **A** Sequence of *z*-projected LLSM images through metaphase and the metaphase-anaphase transition (movie duration 724.5 s (2 s/frame)). Scale bar 2 microns. **B** The number of kinetochores tracked through time (red). The number of kinetochore pair trajectories at each time point (blue; only trajectories with both KTs tracked for at least 80% of the movie are shown). The dashed grey line indicates 92 kinetochores, the number in RPE1 cells. **C** Track overlay time course showing displacement of kinetochore positions from the metaphase plate for single cell; red and blue tracks demarcate KTs that descend to respective daughter cells. **D** *yz* overlay of tracks in metaphase, viewed from above the metaphase plate. **E** Individual kinetochore pair tracks over time, red and blue show tracks of each KT for one randomly chosen cell.

As is typical for mammalian cells, (see HeLa cells, [Jaqaman et al., 2010, Armond et al., 2015a]), KTs form a metaphase plate (Figure 1A) and undergo saw-toothed oscillations perpendicular to the metaphase plate (Figure 1E) before separating in anaphase when KTs segregate towards their respective spindle poles (Figure 1C). RPE1 cells oscillate in metaphase with a period of 84s and have a median inter-sister distance (*i*.*e*., Kinetochore-Kinetochore (KK) distance) of 1.04 microns during metaphase, averaged over cells and time, Figure 2A,C. Oscillations of the KK distance (“ breathing”) have double the frequency as expected, with a period of 44s, Appendix Figure B7. By aligning a cell’s KT trajectories to the median anaphase onset time of a cell (see Methods), we can quantify changes over time as anaphase is approached, *i*.*e*., map metaphase maturation. We confirmed that the MPP becomes thinner over time, as measured by both the compaction of the full KT complement, Figure 2D,E, and the sister mid-points (as used in Jaqaman et al. [2010]), Figure 2H,I. The sister KK distance reduces slightly over metaphase; it begins to increase sharply as some kinetochore pairs initiate anaphase (Figure 2F,G). We confirmed these changes by splitting metaphase into mid-metaphase (330-230 s before anaphase) and late metaphase (130-30s before anaphase, avoiding dynamics immediately prior to anaphase), showing that both the KK distance and the MPP width significantly decrease, (*p*_*MW*_ *<* 10^−3^), Appendix Figure B2. However, the average oscillation period and strength is invariant mid to late metaphase, Figure 2B. Thus, the MPP width primarily decreases because sister pair oscillations centralise to the plate, with a smaller contribution from a reduction in oscillations amplitude (reduced average KK distance).

**Figure 2.**
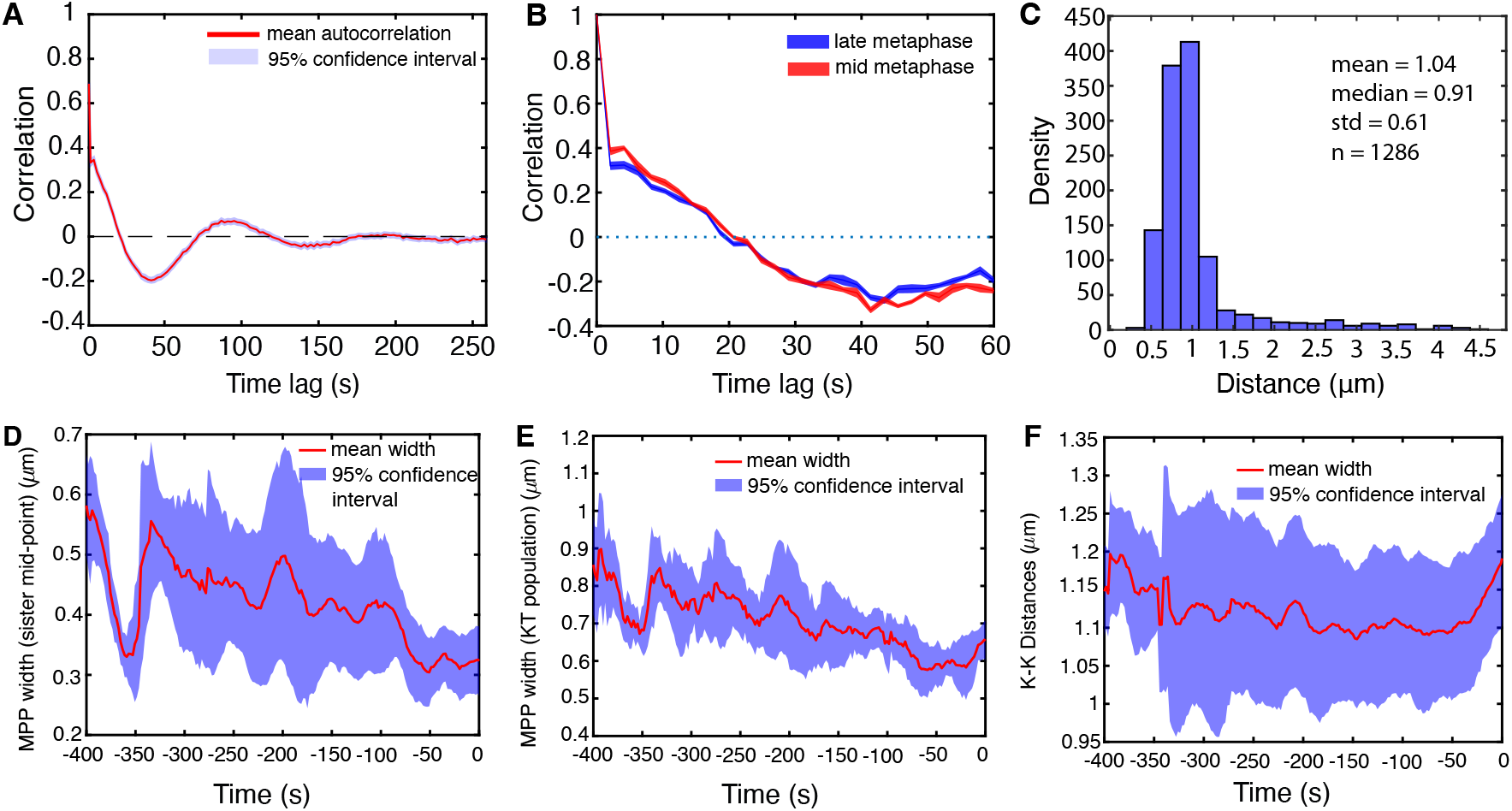
Intrametaphase maturation. **A** Autocorrelation function (ACF) of sister pair midpoints. **B** ACF of metaphase oscillations for late (red) and mid (blue) metaphase. **C** Inter-sister kinetochore (KK) distance pooled over all KT pairs and time. **D-F** Mean (red) and standard deviation (blue) of **D** MPP width as measured by paired sister mid-point width (smallest eigenvalue of the covariance matrix of kinetochore mid-points). MPP statistics, **E** Metaphase plate (MPP) width as measured by the covariance matrix of the KT population (smallest eigenvalue), **F** KK distance against time to anaphase. Data are based on 31 cells having at least 30 sisters both tracked for 75% of movie.

### 2.2 Modelling metaphase kinetochore dynamics

There are 4 forces acting on chromosomes, Figure 3A: the K-fibers can either push or pull the chromosomes, pulling being the substantially stronger force, [Armond et al., 2015a], the centromeric spring attaching the chromatids can be stretched/compressed thus generating an intersister force, centralising forces that push the chromatids/KTs towards the cell mid-plane, and drag forces that damp movements. Following Armond et al. [2015a], we model the dynamics of the two KT sister positions perpendicular to the MPP. Using force balance the dynamics are formulated as a pair of stochastic differential equations, but since time measurements are equispaced, with time interval Δ*t*, we can integrate over the Δ*t* (assumed small) giving a discrete time dynamics for the sister distances, 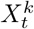, sister *k*, perpendicular to the MPP, (the *vanilla* model),

**Figure 3.**
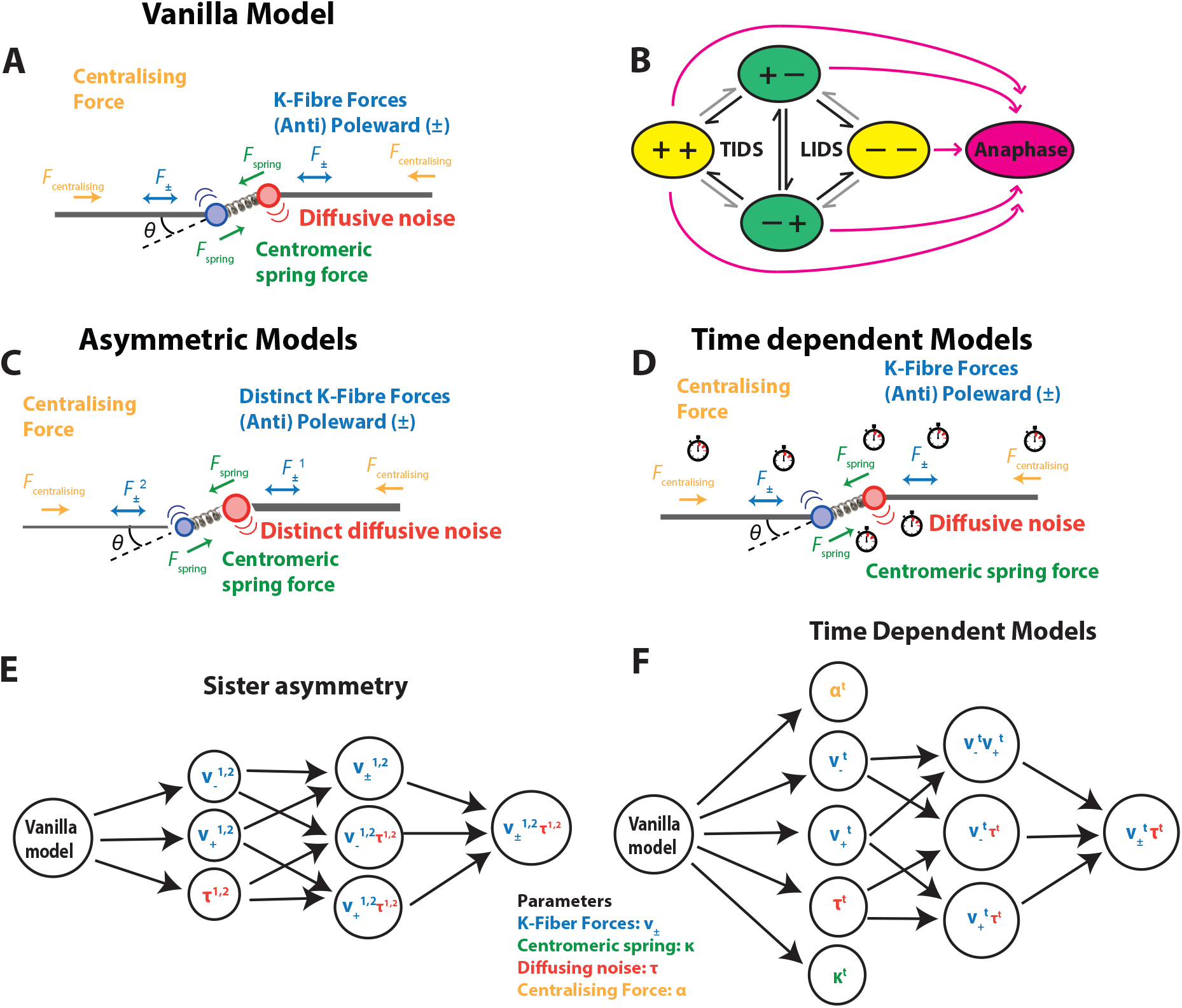
Biophysical model schematics for KT dynamics in metaphase and transition to anaphase. **A** Vanilla metaphase model showing the forces on a KT pair: K-fiber forces (either pulling (poleward) or pushing (anti-poleward)), spring force (green), centralising force (orange) and diffusive noise (arcs). Drag force not shown. **B** Schematic of hidden states comprising sister pair K-fiber states (+ pushing, − pulling). Transitions between states are either initiated by the leading sister (Leading sister Induced Directional Switch, LIDS) or the trailing sister (Trailing sister Induced Directional Switch, TIDS); a joint switch (both sisters switching direction in the same frame) is rarer. Saw tooth oscillations arise because the coherent states (+−,−+) are of longer duration. Transitions in black depict the first sister switching during a directional switching event, gray the second. Anaphase onset is an irreversible transition (pink), *i*.*e*., Anaphase is a absorbing state. **C** Sister asymmetry model variants where sister K-fiber forces (*v*_+_ or *v*_−_) and/or diffusive noise may be unequal. **D** Model for time dependence in the biophysical parameters. **E** Nested model graph of the asymmetric models. **F** Nested graph of time dependent models. Colours refer to biophysical parameters as key in **A**.

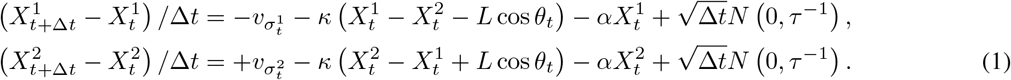

Here 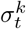 is the state of sister *k* attached K-fiber, either pushing (+) or pulling (−), with a dynamic that is discussed below. The 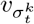 term (taking values *v*_+_ or *v*_−_) is the K-fiber force acting on sister *k* at time *t*. It is a composite force with components from the plus end (de)polymerisation dynamics and the K-fibre flux; *i*.*e*., 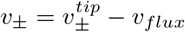 where *v*_*flux*_ is the K-fibre retrograde flux, and 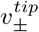 is the tip (plus end) polymerisation, respectively depolymerisation force, (note *v*_−_ ≤ 0, *v*_+_ ≥ 0). The term 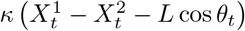 is a Hookean spring force modelling the centromeric chromatin spring (assumed linear) connecting a kinetochore pair, spring constant *κ*, the natural length *L*, and *θ*_*t*_ is the angle between the normal to the metaphase plate and the vector connecting a sister pair at time *t* (thereby projecting the spring force perpendicular to the metaphase plate), Figure 3A. The *αX*^1^ term models the centralising forces, a composite of the polar ejection force (PEF), (K-fiber) length dependent flux Risteski et al. [2022] and HURP dependent control of plus end dynamics (also length dependent, Dudka et al. [2019]), which all produce forces that push KTs/chromatids towards the cell mid-plane. We use a linear model for the centralising forces, linear in the displacement from the metaphase plate. Since the displacement is small relative to the length of the spindle and centralising forces are weaker than the K-fiber forces, Armond et al. [2015a], this will be a good approximation (this is the leading order of a Taylor expansion, the next order being cubic by symmetry). We assume diffusive noise, giving the Gaussian noise distribution *N* (0, Δ*tτ* ^−1^), mean 0, variance parametrised by precision *τ* (inverse of variance). All forces are divided by the unknown drag coefficient as in Armond et al., 2015; the effect of this is that all terms in equation (1) have dimensions of speed, and the force parameters *α* and *κ* have units [*s*^−1^]. The drag force is also a composite, with contributions from viscose drag from the cytosol and drag from the MT matrix.

The hidden states 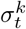 evolve as a discrete time Markov chain parametrised by *p*_*coh*_ and *p*_*incoh*_, the probabilities of a KT remaining in the coherent (sisters move in the same direction, states +−, −+) and incoherent (sisters move in opposite direction, states ++, −−) state over a time interval Δ*t*, respectively, Figure 3B. Specifically, the hidden states are described via the transition matrix *P*, with (states *σ* with ordering {++, +−, −+, −−}),

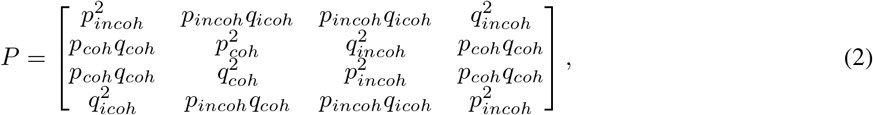

with *q*_*coh*_ = 1 − *p*_*coh*_ and *q*_*incoh*_ = 1 − *p*_*incoh*_. Simulating from this biophysical model produces trajectories with quasi-periodic oscillations qualitatively similar to observed data, [Armond et al., 2015a]; saw-tooth like oscillations occur when the coherent mean lifetime is larger than the incoherent lifetime.

#### Bayesian inference of biophysical parameters

We take a Bayesian approach fitting the models directly to each experimental trajectory, a method that infers all the biophysical model parameters jointly. In essence, Bayesian methods sample parameter values consistent with observed data. Crucially, this allows us to quantify uncertainty in the fitted parameters. We use the STAN programming language through “ rSTAN” package, [Stan Development Team, 2024b, Carpenter et al., 2017] that implements an Hamiltonian Markov Chain Monte Carlo algorithm, [Betancourt and Girolami, 2015, Neal, 2011], an algorithm that generates samples of the posterior parameter distribution *P* (*θ*|*x*_1:*T*_), the distribution of the model p arameters *θ* = (*τ, α, κ, v*_−_, *v*_+_, *L, p*_*coh*_, *p*_*incoh*_) (model (1), (2)), given the observed data, a sister pair trajectory 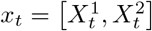in our case. Details on the likelihoods for each model, how we deal with missing data and the parameter and hidden state sampling (from the posterior distribution) are given in the Appendices C2 and C3. There is an identifiability issue in all models such that the natural length *L* is poorly inferred from the data. This is discussed in Armond et al. [2015a] and resolved by measuring the natural length in nocodazole (which depolymerises all MTs); the average natural length (KK distance) for RPE1 cells is 0.78 microns, similar to HeLa cells at 0.76 microns, [Armond et al., 2015a]. All the models analysed in this paper then satisfy practical identifiablity [Hines et al., 2014, Browning et al., 2020].

We imposed additional tracking requirements for the inference, in particular removing trajectories with over 20% missing data, Appendix C1. This gave 26 cells for the model analysis. Inference was successful on the vast majority of trajectories; a small number of KT pairs (∼1.5%) had severe divergences, likely due to the model being inappropriate, *i*.*e*., not capturing the dynamics. Such trajectories are removed from the analysis. Hence, in the model analysis there are slight differences in the number of trajectories considered due to performance differences.

#### Determining which models are supported by the data

Model selection is based on pairwise comparison of models using the Bayes factor, all model comparisons being nested; thus we assess the evidence for increasing the complexity of the model. The Bayes factor for model *M* ^′^ relative to a simpler model *M* is the fraction 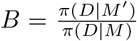 given data *D*; the model marginals *π*(*D* | *M*) are computed using a bridging sampler, Gronau et al. [2020]. We use four strength of evidence criteria against the null hypothesis as defined by Kass and Raftery [1995a]; “ Not worth than a bear mention” *i*.*e*., *BF <* 3.2, “ Substantial” *i*.*e*., 3.2 ≤ *BF <* 10.0, “ Strong” *i*.*e*., 10 *< BF* ≤ 100, and “ Decisive”, *i*.*e*., *BF* ≥ 100. We only chose the more complex model if it had at least substantial preference over the simpler model (Bayes factor *BF >* 3.2). We initially determine which models are supported relative to the vanilla model (eq. (1)), giving a set of preferred models. Within this set, any nested models were then assessed for support for increasing the complexity using the Bayes factor between pairs of models in the nesting, Figure 3E,F, removing the more complex models for which increased complexity is not supported. If multiple models remain, we select the model with the highest Bayes factor (relative to the basic model) as the preferred model. In this way we navigate the model network to determine the model with the greatest support, and with model complexity justified by the data. This iterative process allows us to navigate the model space systematically, identifying the model with the strongest support while ensuring it’s complexity is justified by the data.

### 2.3 Biophysical characteristics of quasi-periodic oscillations in diploid non-transformed human RPE1 cells

We use the *vanilla* model (eq. (1)), (2) to infer all the biophysical model parameters concurrently, see Appendix C. We analysed (after data quality filtering) 26 cells, 798 KT pairs (average of 31.9 sister pairs/cell). The inferred (marginal posterior) distributions for each of the biophysical model parameters are shown in Figure 4 for a single sister pair trajectory. A comparison between the prior and posterior marginals demonstrates that all parameters are identifiable except for the natural length *L* which has a posterior nearly identical to the prior, expected given its identifiability issues, [Armond et al., 2015a]. The model fit to the trajectory provides an automated state annotation, *i*.*e*., the hidden K-fiber pulling/pushing state is inferred at each time point, Figure 4B, thereby allowing switching events of the KT sister pair state to be inferred, Figure 4C. Metaphase quasi-periodic oscillations require that both KTs of a sister pair change direction at a directional switch, [Armond et al., 2015a]. There are two switching choreographies; a switching event initiated by the leading sister (lead induced directional switch, or LIDS) and switching initiated by the trailing sister (trail induced directional switch, or TIDS), Figure 3B; coincident switching (within the time resolution) is rare. In this trajectory directional switching (between coherent states +− to −+, or −+ to +−), occurs through LIDS events, *i*.*e*., the intermediate state is ++, compressing the centromeric spring. Both choreographies are in fact observed with a LIDS/TIDS ratio of 3.2, specifically 68.5% LIDS, and 21.1% TIDS and 10.3% joint switches (26 cells, 798 KT pairs). This is similar to that previously reported for HeLA cells, with a LIDS/TIDS of 3.8, [Armond et al., 2015a].

**Figure 4.**
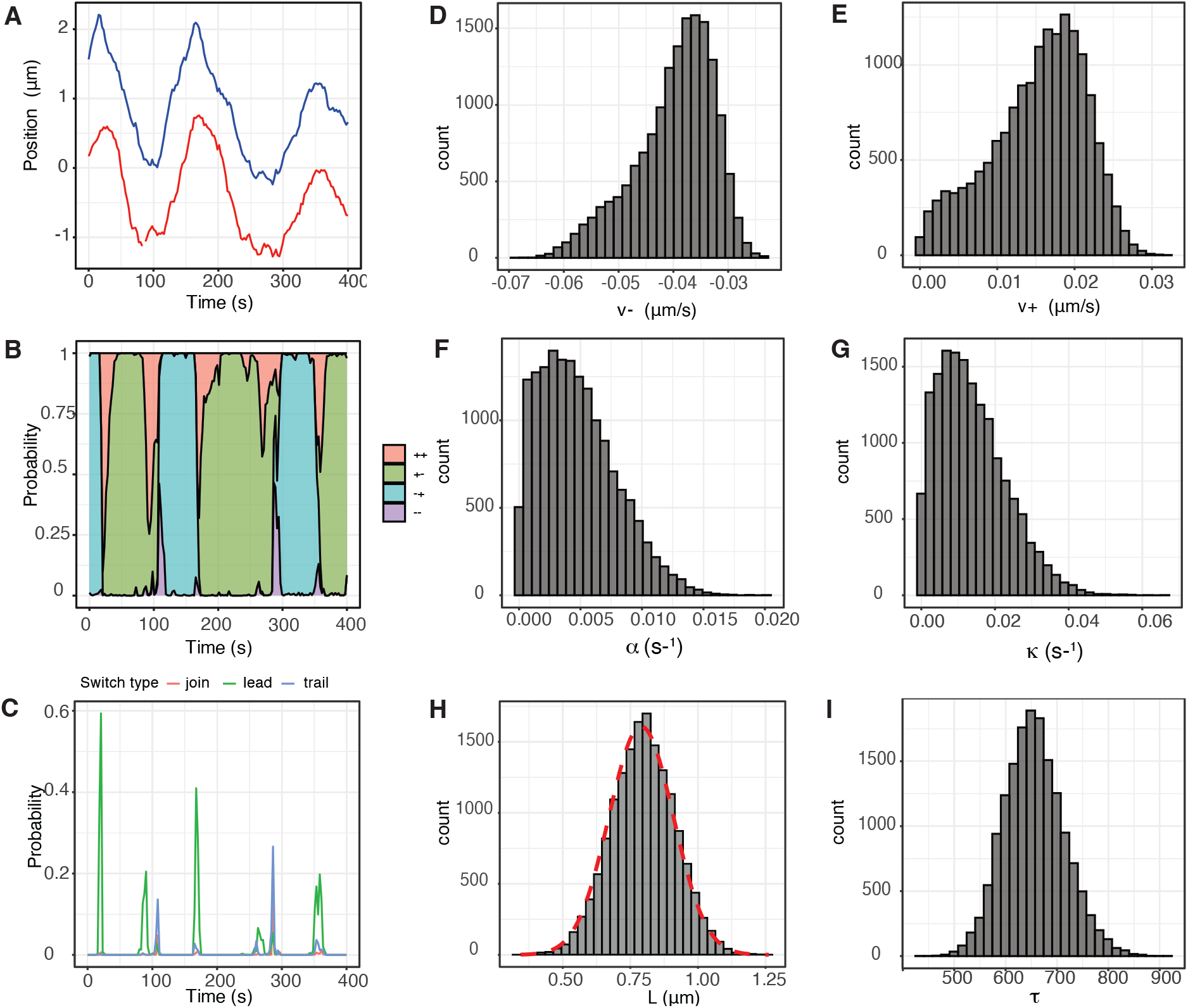
Single KT pair trajectory inference showing model parameter estimates and trajectory annotation. **A** KT distance transverse to the MPP for the 2 sisters (red, blue). (LLSM imaging of Ndc80-eGFP endogenously marked KTs). Additional data for this KT pair is given in Appendix Figure B6. **B** Trajectory annotation for the hidden K-fiber polymerisation states; probability *P* (*σ*_*t*_ |*x*_1:*T*_; *θ*) for being in each state at each time point as sampled via the backward algorithm (see Methods). **C** Probability of a directional switch initiated by the leading kinetochore (green), trailing kinetochore (grey), or a joint switch (red). Switching probability is assessed using the sampled hidden states and corresponds to a proportion of MCMC samples matching a particular pattern of states (e.g. LIDS [−+,−+,++,+−]) corresponding to a given switch type. **D-I** Marginal posterior distribution of the stated biophysical parameter for the trajectory data in **A**. In **H** the informative prior density plot for natural length *L* is plotted with red dashed line on top of posterior histograms. All other parameters have uninformative priors.

Inferred biophysical parameters for RPE1 cells are given in Table 1. We note that these are similar to previously inferred parameters in HeLA cells Armond et al. [2015a].

**Table 1:** Inferred biophysical parameter values of all sister KT-pairs using the vanilla model, Eq. (1) (based on Armond et al. [2015a]). The first column shows the median of posterior medians with interquartile ranges in brackets. The second column shows the median of the Posterior Interquartile Range (PIR), typically less than the IQR. Based on 26 cells and 798 KT pairs.

| Parameter | Median [IQR] | Median PIR |
| --- | --- | --- |
| $\tau$ | 673.7 [241.4] | 95.04 |
| $\kappa$ ($s^{-1}$) | 0.022 [0.018] | 0.018 |
| $\alpha$ ($s^{-1}$) | 0.009 [0.008] | 0.005 |
| $v_-$ ($\mu m/s$) | -0.041 [0.021] | 0.008 |
| $v_+$ ($\mu m/s$) | 0.010 [0.010] | 0.007 |
| $L$ ($\mu m$) | 0.808 [0.033] | 0.158 |
| $p_{incoh}$ | 0.760 [0.216] | 0.155 |
| $p_{coh}$ | 0.949 [0.027] | 0.026 |

### 2.4 Sister kinetochores exhibit substantial sister asymmetry in kinetochore forces

The two spindle poles are in fact not equal, respective spindle poles having a young and old centrosome. Centrosome age affects the stability of MTs from the associated spindle pole which imparts a half-spindle bias, both in positioning of unaligned KTs and missegregation, [Gasic et al., 2015]. Thus, assuming KT sister dynamics is symmetric is not justified. Our vanilla model assumes sister KTs and K-fibers are biophysically identical, *i*.*e*., have the same pulling and pushing forces. Here, we relax the assumption of sister symmetry in the K-fiber forces and their diffusive noise, *i*.*e*., the sisters can have distinct (unloaded) forces 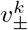 and noise (precisions *τ*^*k*^) in eq. (1), *k* = 1, 2, Figure 3C. Clearly *κ* is common. We do not consider asymmetry in *α* here, but note that asymmetry in *α* could arise from asymmetry in the PEF due to an asymmetric spindle, or asymmetry of the centralising flux. We confirm later that there is no half-cell bias in *v* _±_ supporting the assumption that the spindle poles are of equal strength. The full asymmetric model we consider is given by, (generalising model eq. (1))

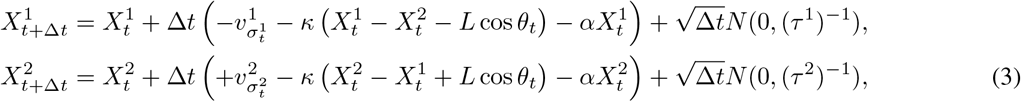

the hidden states 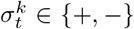, have the same dynamics as above. This model allows sisters to differ in 3 variables (*v* _±_, *τ*); we also consider the reduced models with only 1 or 2 parameters being different, giving 7 asymmetric models and the symmetric/vanilla model (eq. (1)); model relationship is shown in Figure 3E. A more complex model is deemed to be supported from the data if the Bayes factor *B >* 3.2, see Appendix C. We report results of the following 5 models:

- *M*_0_ no asymmetries, *i*.*e*., symmetric - vanilla model (eq. (1)).
- 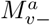 asymmetry only on *v*_−_
- 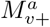 asymmetry only on *v*_+_
- 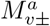 asymmetries on *v*_−_ and *v*_+_
- 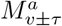 asymmetries on *v*_−_, *v*_+_ and *τ* with the superscript *a* denoting asymmetry models and the subscript which parameter(s) are asymmetric. The other models were rarely preferred, so not discussed further. The model likelihoods these 5 models are given in Appendix C.

We fitted the 4 asymmetric sister models to 798 sister pairs, across 26 cells. In RPE1 cells, 24.4% of sister kinetochores have significant asymmetry, with most, (45.1% of asymmetric pairs) preferring asymmetry in *v*_−_*v*_+_ whilst 88.7% have significant asymmetry in *v*_−_ (preferring either of the models 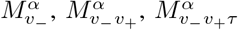), Figure 5A, Appendix

**Figure 5.**
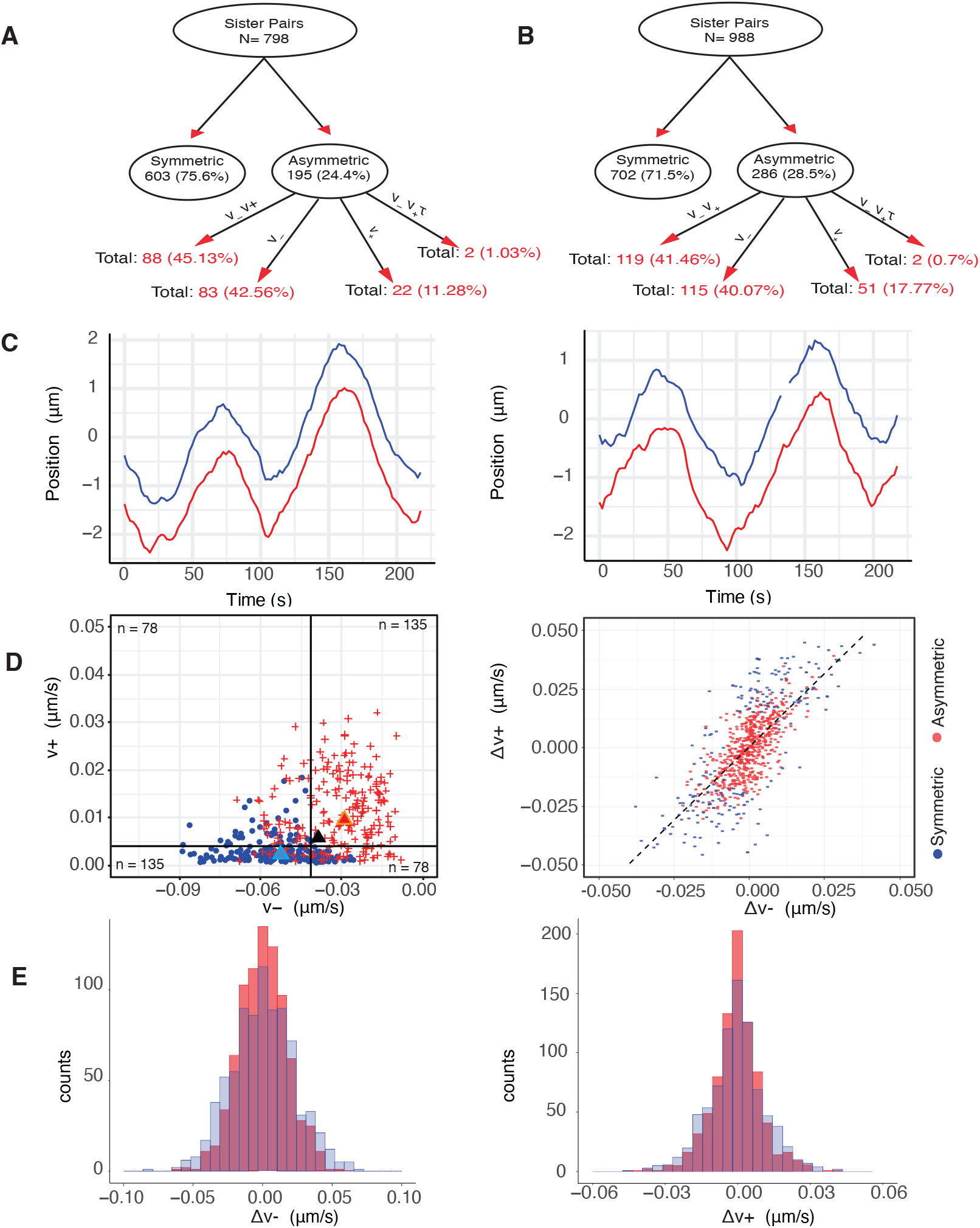
Asymmetry in sister kinetochores is a result of natural variation (noise). **A/B** Graphical representation of asymmetry model preference network with decomposition shown as percentage of parent node in DMSO (A) and Nocodazole (B) treated cells. **C** Typical symmetric (left) and asymmetric (right) trajectory, DMSO treated. No distinctive characteristics between the symmetric and asymmetric sister pairs. **D** (Left) Comparison of inferred *v*_−_ and *v*_+_ of asymmetric sister pairs (on asymmetric *v*_±_ model). For each pair, the stronger and weaker pulling sister are shown, blue and red respectively. Population medians are marked with triangle, while the black triangle shows the population median of the symmetric pairs. The number of KTs in each quadrant is shown in every quadrant corner. (Right) Scatter plot of sister difference in pulling force 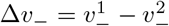 versus pushing force 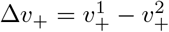, with symmetric and asymmetric sister pairs indicated in red and blue respectively, (inferred on the *v* _±_ Asymmetric pairs do not present any distinctive pattern. **E** (left) and of *v*_±_ asymmetric model). Histograms of the sister difference distribution, *i*.*e*., Δ*v*_−_ Δ*v*_+_ (right), when sisters are paired (red) and pairing is randomised (blue), (randomisation is restricted to a metaphase radial box to account for variation with respect to the metaphase plate distance, see Figure 7).

Table A1. However, (significantly) asymmetric sisters only show mild differences in biophysical parameters relative to symmetric sisters, Table 2, asymmetric and symmetric trajectories in fact appear similar, Figure 5C. Asymmetric sisters have a significantly different (larger) average pulling force (averaged over the two sisters) than symmetric sisters 9.3% (*p*_*MW*_ *<* 10^−7^, one-sided)^1^, whilst the centromeric spring constant, mid-plane centralising forces and natural length are 15.9 % (*p*_*MW*_ = 0.01), 34.3% (*p*_*MW*_ *<* 10^−7^) and 2.6% (*p*_*MW*_ *<* 10^−6^) larger. Since it is easier to detect differences when | *v*_−_| is larger, the asymmetric group would be expected to have a larger average *v*_−_. In all cells there is a fraction of (significantly) asymmetric sisters, see Appendix Figure B5A,B, while, there is also no half-spindle bias in the (significantly) asymmetric KTs, *p*_*Binom*_ = 0.88, *i*.*e*., cells exhibit equal numbers of stronger KTs to the right, respectively left, half-spindle, Appendix Figure B5C,D. Thus, kinetochore asymmetry is not due to a spindle asymmetry that can arise because of a younger and older centrosomes, Gasic et al. [2015].

**Table 2:** Biophysical parameters of symmetric and (significantly) asymmetric sister pairs. Sister 1 is the sister with the greater pulling force in an asymmetric pair (*i*.*e*., 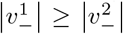). The first and third columns report the medians of posterior medians for symmetric and asymmetric sisters respectively; the population interquantile range is shown brackets. The average parameter confidence per trajectory is shown in the second and fourth columns, specifically the median of Posterior Interquantile Ranges (PIR) for symmetric and asymmetric sisters, respectively. Based on Mann-Whitney and Kolmogorov-Smirnov tests, the asymmetric posterior distributions of *κ*,*α, v*_−_ and *L* are significantly different than the symmetric posterior distributions, with size of effect (*i*.*e*., percentage difference of the medians) being 15.9%, 34.3%, 9.5% and 2.6%. Inference on asymmetric sisters is based on 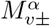.

|  | Symmetric |  | Asymmetric |  |
| --- | --- | --- | --- | --- |
|  | Median | PIR | Median | PIR |
| $\tau$ | 661.8 [265.4] | 95.055 | 700 [311.9] | 96.90 |
| $\kappa$ | 0.020 [0.018] | 0.017 | 0.023 [0.022] | 0.017 |
| $\alpha$ | 0.010 [0.008] | 0.005 | 0.013 [0.010] | 0.005 |
| $v_-^1$ | -0.042 [0.021] | 0.008 | -0.060 [0.020] | 0.009 |
| $v_-^2$ | -0.042 [0.021] | 0.009 | -0.032 [0.026] | 0.009 |
| $v_+^1$ | 0.010 [0.010] | 0.007 | 0.004 [0.005] | 0.007 |
| $v_+^2$ | 0.010 [0.010] | 0.007 | 0.015 [0.016] | 0.007 |
| $L$ | 0.806 [0.033] | 0.156 | 0.839 [0.067] | 0.155 |
| $p_{incoh}$ | 0.734 [0.216] | 0.173 | 0.766 [0.215] | 0.133 |
| $p_{coh}$ | 0.950 [0.027] | 0.026 | 0.952 [0.024] | 0.024 |

We examined if pushing and pulling forces are independent for individual KTs. There is a significant correlation between pushing (*v*_+_) and pulling (*v*_−_) forces in the asymmetric population, Figure 5D, specifically K-fibers that are stronger at pulling are typically weaker at pushing (*p*_*Binom*_ *<* 10^−8^, rejecting the null hypothesis that the strongest pulling sister (*v*_−_) had equal probabilities of being either the stronger or weaker pushing sister (*v*_+_)). Thus, pairs where a sister is both a stronger puller and pusher (respectively weaker puller and weaker pusher) are in a minority. The difference in sister forces, 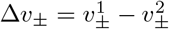, shows this trend on all KT pairs, Figure 5D, suggesting asymmetry may be a natural consequence of KT and K-fiber variation. Sister differences were in fact not significantly different to differences between any 2 randomly chosen KTs, (the force difference distributions were identical, *p*_*MW*_ = 0.986, *p*_*MW*_ = 0.621, for 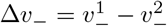 and 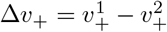, respectively, and *p*_*KS*_ = 0.056 and *p*_*KS*_ = 0.141 for Δ*v*_−_, Δ*v*_+_, respectively), Figure 5E. This also holds when we allow for the effects of location in the MPP (applying the same tests on 5 equally sized groups, based on their averaged location on the metaphase plate, see Section 2.6). The lowest p-value for the tests over the 5 classes was *p*_*MW/KS*_ = 0.385. This is further explored in Appendix Figure E1A.

We conclude that K-fiber forces are intrinsically variable and therefore sister differences (asymmetry) is a continuum. The significantly asymmetric sisters will therefore be those with the largest differences. Notably, the mechanism underlying this KT/K-fiber stochasticity introduces an inherent anti-correlation between pulling and pushing forces.

### 2.5 Symmetric and asymmetric sisters are organised transversely to the MPP

Our analysis reveals that K-fibers are intrinsically noisy, which results in an asymmetry in sister KTs in *v*_±_ and in the diffusive noise *τ*, Figure 5. Sister asymmetry in pulling forces should localise the sister pair off centre of the MPP plate, the K-fiber pulling force being the dominant force acting on the chromosome, [Armond et al., 2015a]. Specifically, a sister pair with a substantially stronger pulling sister will be pulled on average towards its pole, experiencing on average a higher centralising force away from that pole, the pair thus achieving a stable average off-centre position towards the stronger sister’s pole. This is in fact observed, significantly asymmetric sister pairs are localised on average towards their pole relative to symmetric sisters, Figure 6A. To ascertain if this repositioning is retained through anaphase, we back-tracked the KTs that descend to respective poles tracking the cluster back through metaphase. Within each poleward destined cluster, we can partition the KTs into 3 groups, firstly those that are part of a symmetric pair, secondly the stronger pulling sister of an asymmetric pair and finally the weaker pulling sister of an asymmetric pair. The stronger pulling sisters are positioned towards the pole in metaphase, Figure 6C, as expected since the sister centre is offset towards the stronger sister’s pole, an offset that is significant relative to the KTs from symmetric pairs, *p*_*MW/KS*_ *<* 10^−16^. This offset is in fact retained through into anaphase, Figure 6E. Similarly, the KTs that have the weaker pulling force of an asymmetric pair are positioned away from the pole to which they descend, *p*_*MW/KS*_ *<* 10^−14^, Figure 6C,E. An analysis of the KT position ranking within its cluster (ranked mean position towards the pole) also demonstrates the relative positioning of the weaker and stronger sisters of an asymmetric pair compared to kinetochores of a symmetric pair, Appendix Figure B17. The relative positioning is in fact fairly stable through metaphase with a small number of KTs changing position – what is surprising is that asymmetric KTs tend to be at the extremes, the majority in their correct position (rank), but a minority diametrically opposite. Our results demonstrate that there is an ordering along the spindle axis (MPP normal) of KTs that is a consequence of unequal K-fiber pulling forces in metaphase, an ordering that is retained through anaphase, Figure 6G. Thus, stronger KTs of a sister pair are positioned in the front of the cluster, weaker KTs at the back in metaphase and anaphase.

**Figure 6.**
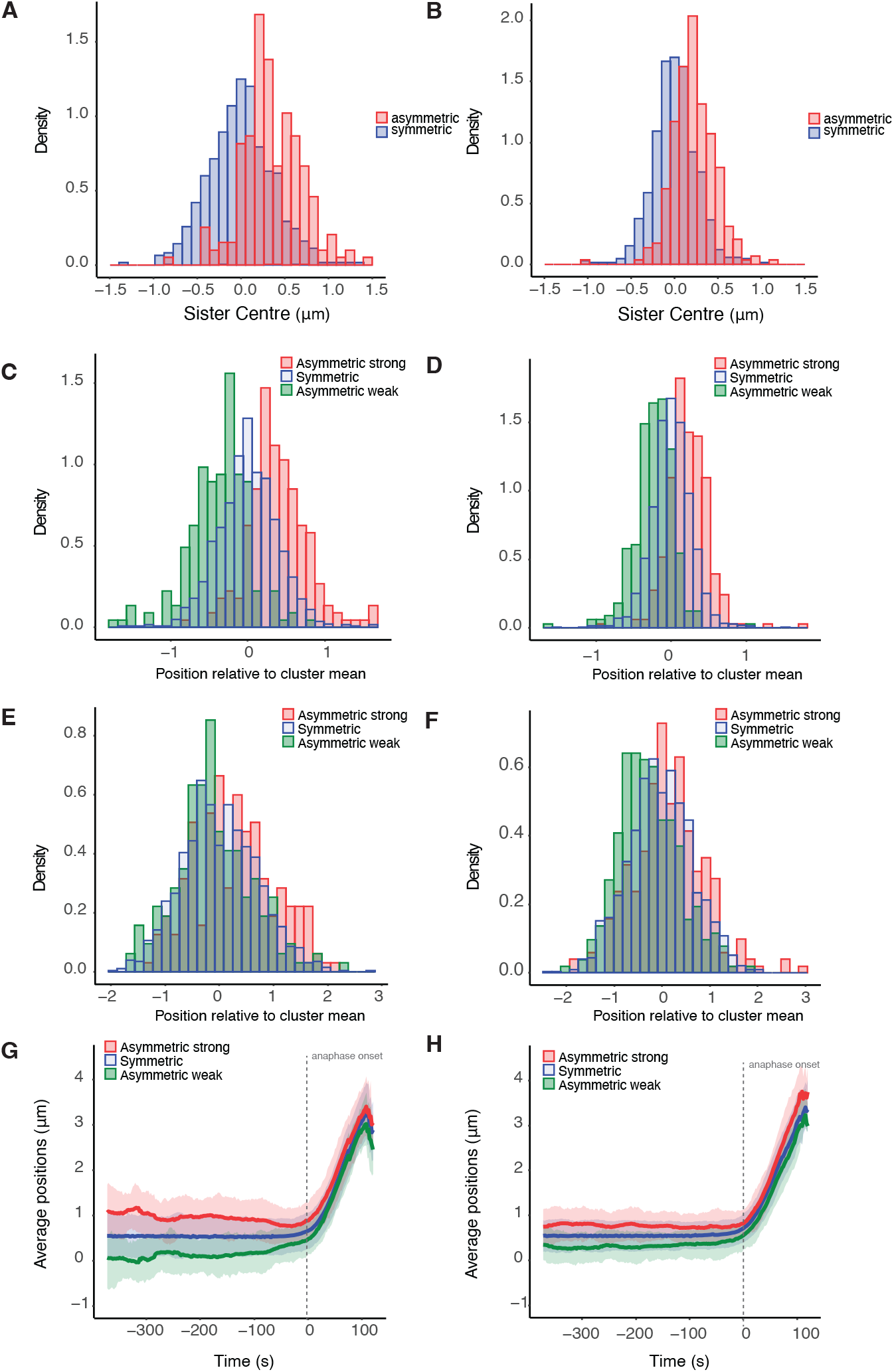
Symmetric and asymmetric kinetochores are organised transverse to the MPP. Transverse organisation of the MPP by pulling strength group for: left (**A**,**C**,**E**,**G**) DMSO (195 asymmetric and 603 symmetric sister pairs), right (**B**,**D**,**F**,**H**) nocodazole washout treated cells (286 asymmetric and 702 symmetric sister pairs). **A/B** KT pair average position (over sisters and time) normal to the MPP. For asymmetric KT pairs, the distance is measured towards the stronger pulling sisters’ pole. **C/D** Mean positions of sisters relative to their relevant cluster cluster, during metaphase. Stronger (weaker) asymmetric asymmetric sisters are positioned close (away) to their pole. **E/F** Mean positions of sisters relative to their cluster mean in anaphase. Cumulative distributions of asymmetric stronger pulling is significantly greater than cumulative symmetric distribution which in turn is statistically greater than asymmetric weaker pulling (*p*_*KS*_ *<* 10^−6^ (*p*_*KS*_ *<* 10^−4^), DMSO (nocodazole washout)). In addition, the median of asymmetric stronger pulling is significantly greater than the median of symmetric sisters which in turn is significantly greater than the median of asymmetric weaker sisters for both DMSO and nocodazole washout treated cells, (*p*_*MW*_ *<* 10^−5^, *p*_*MW*_ *<* 10^−10^). **G/H** Evolution of the average positions in time. Shaded envelope shows interquartile range for each group. *t* = 0 denotes the median anaphase onset per cell.

### 2.6 K-fiber parameters have substantial spatial variation across the metaphase plate

Previous reports suggest that KT dynamics varies with position in the MPP, in particular that (i) trajectories are more stochastic towards the periphery, [Jaqaman et al., 2010, Armond et al., 2015a] (ii) that the centralising force (PEF) increases towards the periphery of the MPP, [Armond et al., 2015a] and (iii) that sister pairs at the periphery exhibit higher swivel, [Smith et al., 2016]. To determine if there are trends in the biophysical parameters with distance *r* from the centre of the metaphase plate in RPE1 cells, we partitioned the population of KTs with respect to position, as in

Armond et al. [2015a], Figure 7. There are substantial trends with *r* in a number of parameters, specifically the pulling force *v*_−_ (*p*_*Corr/Kendall/Spearman*_ *<* 10^−16^, Figure 7C) and pushing force *v*_+_ (*p*_*Corr*_ *<* 10^−7^, *p*_*Kendall/Spearman*_ = 0.002, Figure 7D), the spring constant *κ* (*p*_*Corr*_ *<* 10^−8^, *p*_*Kendall/Spearman*_ *<* 10^−13^, Figure 7F) and precision *τ*, (*p*_*Corr/Kendall/Spearman*_ *<* 10^−16^, Figure 7H). These trends are also detectable in single cells, Appendix Figure B10. In contrast to previous reports (on HeLA cells), there was no trend in the centralising force, which was invariant with *r* (*p*_*Corr*_ *<* 0.48, *p*_*Kendall/Spearman*_ = 0.12), see Appendix Table A5 for a summary of p-values. In the previous study, Armond et al. [2015a], no trends in the other dynamic parameters were observed in contrast to those observed here in RPE1 cells. The coverage in that study was substantially lower than in our study, which may explain these differences; however, they could also reflect differences between RPE1 and HeLa cells. Chromosome size is known to increase towards the periphery, [Mosgöller et al., 1991, Booth et al., 2016], which could increase drag forces towards the periphery, whilst drag may also be dependent on MT density, that may vary laterally across the spindle. A change in the drag coefficient with *r* scales all the mechanical parameters as follows, generalising eq. (1):

**Figure 7.**
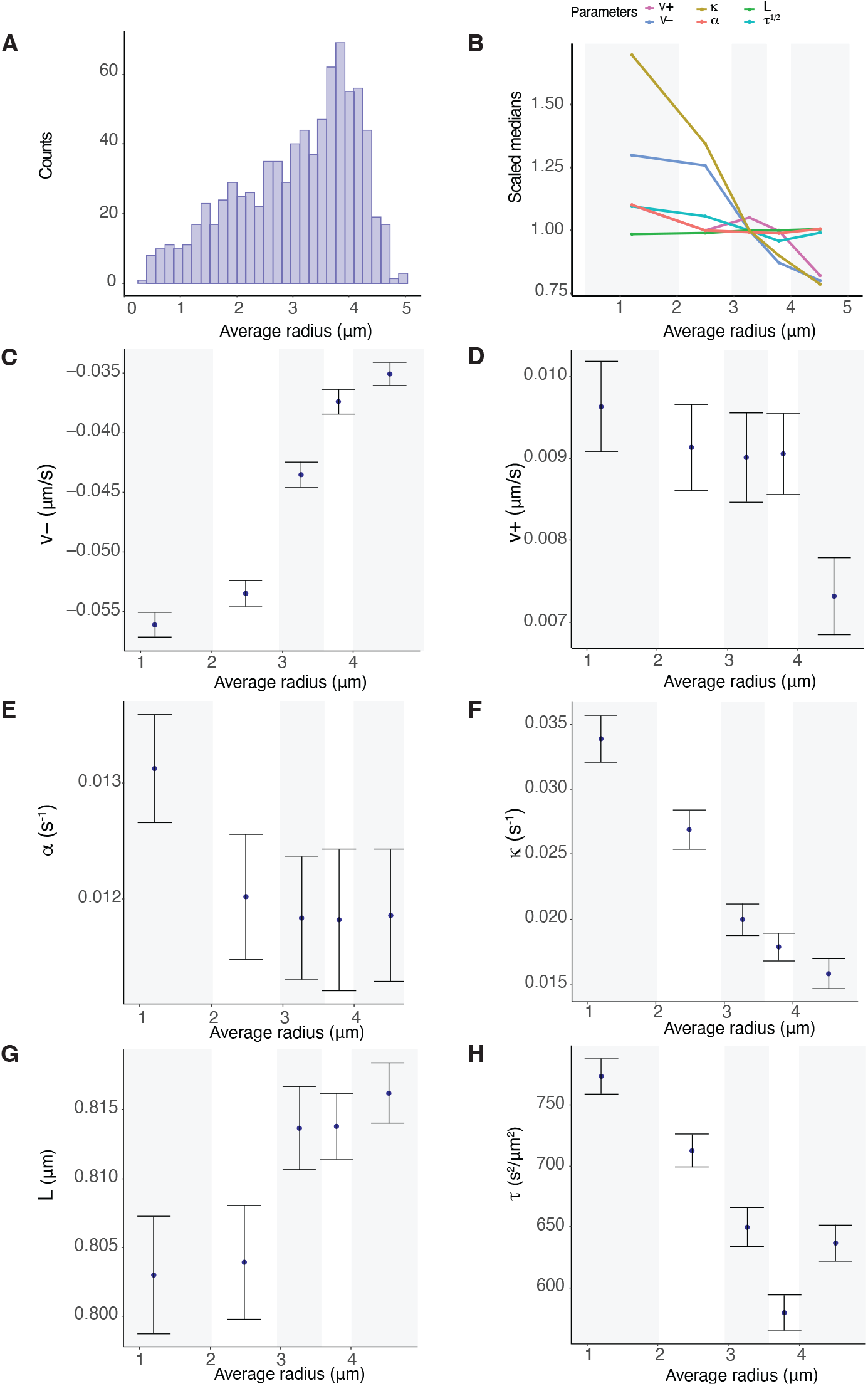
K-fiber pulling strength and centromeric spring stiffness decreases towards the periphery of the metaphase plate. **A** Location of kinetochore pairs within the metaphase plate across 26 cells. **B-H** Summary statistics (medians of posterior medians and standard error) of indicated biophysical parameters partitioned into 5 groups for the average radial position within the metaphase plate. Groups are equally sized, containing the same number of observations, with the five groups defined by their radial distances, *i*.*e*., [0, 2.03], (2.03, 2.95], (2.95, 3.58], (3.580, 4.01], (4.01, 5.03] respectively. **B** Summary of scaled spatial trends over all parameters. **C** Pulling forces *v*_−_, **D** Pushing forces *v*_+_, **E** Spring constant *κ*, **F** Centralising force, **G** precision *τ*, **H** Natural length of the centromeric chromatin spring connecting the kinetochore sisters *L*. Based on 798 KT pairs over 26 cells.

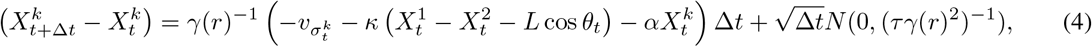

Thus, *v* _±_, *κ, α, τ* ^1*/*2^ would scale identically with *r*. The scalings are compared in Figure 7B. There is substantial variation in the relative reduction in the parameters (percentage difference peripheral to middle of MPP 38%, 25%, 9%, 53% and 2% of *v*_−_, *v*_+_, *α, κ* and *L* respectively), whilst an opposite trend is seen for *τ* ^1*/*2^, which increases by 10%, suggesting that solely changing drag across the MPP is not consistent with the data. The relative changes in the parameters are also not consistent with changing both drag and chromosome size across the MPP; a change in chromosome size would affect both the drag and the PEF.

Biological variation is expected both within cells, resulting in differences in the mechanical properties of individual KTs and K-fibers, and between cells, with the between cell variation expected to be substantially larger. However, the variation we see with respect to location in the metaphase plate in our pooled cell samples indicates that within-cell variation is in fact substantial since it is evident even on data pooled over cells. To ascertain the contributions to the variance from spatial location and biological variation between cells we performed a 2-way ANOVA on the median estimates per sister pair for each parameter, Table 3, with a 2-way grouping with respect to the 5 radial partitions and 26 cells. The interaction term (between radial location and cell) was always insignificant (in all parameters), so removed from the ANOVA analysis. In all parameters cell grouping was significant, whilst MPP location was significant in all except *α*. However, the variance contributions from location and cell showed high variability; the location variance of *v*_−_ is about double the cell variation, whilst the location variance of switching parameter *p*_*incoh*_ is also substantially larger indicating that the duration of directional switching is heterogeneous with location, Table 3. All other parameters have similar (or smaller) variance contributions from location compared to between cells (the natural length has a strong prior and the size of effect is small despite a significant difference between cells, whilst it also has negligible trend with *r*). The higher between cell variance of *α* suggests that the centralising force varies substantially between cells, whilst K-fibre pulling force and the centromeric spring varies substantially within cells, consistent with the strong trends seen in Figure 7. The relative (total) standard deviation (sd/mean) shows that variation of *v*_±_, *α, κ* is substantial (over 30%), clearly evident in the parameter estimates by cell, Appendix Figure B8.

**Table 3:**
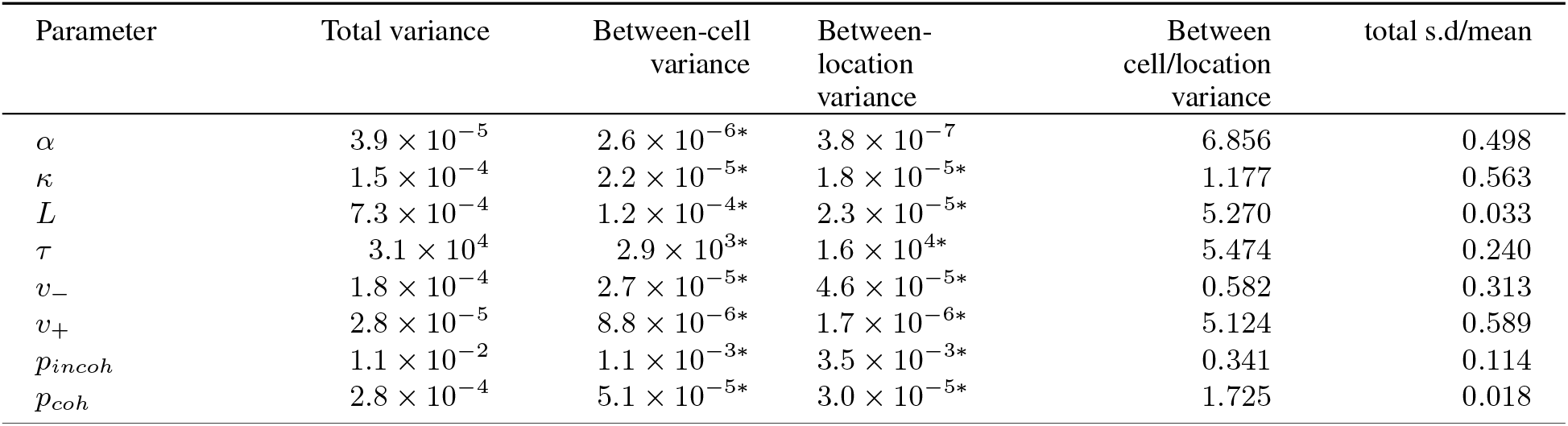
K-fiber force variation within cells is larger than between cells. Variance contributions of the biophysical parameters, (*n* = 383 KT pairs from *N* = 14 cells, imposing at least 3 sister pairs in each group), grouping by cell and location (5 radial boxes) within the metaphase plate. The second column denotes the within-cell variance (*i*.*e*., residuals), while the third and fourth columns report the within cell and location variance respectively. Parameter estimations use the model with asymmetry on *v*_−_ and *v*_+_; the posterior median *v*_−_ and *v*_+_ values are averaged over the two sisters. Stars denote that there is statistically significant difference (*α <* 0.01) between groups of this factor. The ratio of between cell to between location variance is shown in fifth column and the size of effect (ratio of population parameters’ standard deviation to the population parameters’ (absolute) mean) in the last column. Stars denote the statistically different ratio at *α <* 0.01. Caution: Due to low coverage of KT pairs per cell within each radius group, these results should be taken with caution as some of the ANOVA assumptions are violated. Results on 3 radial boxes give similar results, see Appendix Table A2.

| Parameter | Total variance | Between-cell variance | Between-location variance | Between cell/location variance | total s.d/mean |
| --- | --- | --- | --- | --- | --- |
| $\alpha$ | $3.9 \times 10^{-5}$ | $2.6 \times 10^{-6*}$ | $3.8 \times 10^{-7}$ | 6.856 | 0.498 |
| $\kappa$ | $1.5 \times 10^{-4}$ | $2.2 \times 10^{-5*}$ | $1.8 \times 10^{-5*}$ | 1.177 | 0.563 |
| $L$ | $7.3 \times 10^{-4}$ | $1.2 \times 10^{-4*}$ | $2.3 \times 10^{-5*}$ | 5.270 | 0.033 |
| $\tau$ | $3.1 \times 10^4$ | $2.9 \times 10^{3*}$ | $1.6 \times 10^{4*}$ | 5.474 | 0.240 |
| $v_-$ | $1.8 \times 10^{-4}$ | $2.7 \times 10^{-5*}$ | $4.6 \times 10^{-5*}$ | 0.582 | 0.313 |
| $v_+$ | $2.8 \times 10^{-5}$ | $8.8 \times 10^{-6*}$ | $1.7 \times 10^{-6*}$ | 5.124 | 0.589 |
| $p_{incoh}$ | $1.1 \times 10^{-2}$ | $1.1 \times 10^{-3*}$ | $3.5 \times 10^{-3*}$ | 0.341 | 0.114 |
| $p_{coh}$ | $2.8 \times 10^{-4}$ | $5.1 \times 10^{-5*}$ | $3.0 \times 10^{-5*}$ | 1.725 | 0.018 |

### 2.7 K-fiber mechanical parameters are time dependent and tune towards an anaphase ready state

The metaphase plate (MPP) undergoes time-dependent maturation in RPE1 cells, including a compaction of the width along the spindle axis as anaphase onset is approached, [Embacher et al., 2022] and Figure 2; a similar decrease in MPP width is observed in HeLa cells [Jaqaman et al., 2010]. We thus investigated whether the biophysical parameters change in time, potentially explaining MPP maturation. We modified the vanilla model, eq. (1), to have time dependent parameters; we assumed time dependence is exponential this preserving the parameters sign whilst it is also a good approximation for weak linear dependence. The full time dependent model has 5 time dependent parameters, Figure 3D,F, and given by,

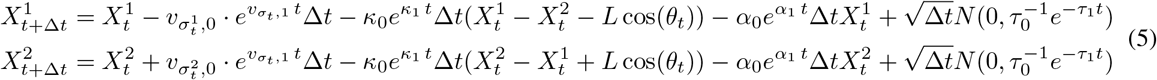

where we define for a time dependent parameter the constant term, with subscript 0, (value at the start of the movie), and rate of change parameter (on a log scale) with subscript 1. We don’t analyse time dependence in switching, as there are too few switching events per trajectory, or time dependence of the centromeric springs’ natural length as that is fixed from an accessory experiment (nocodazole washout) because of identifiability issues. We consider a nested sequence of models, Figure 3F. We label models by the parameters that are time dependent, for example 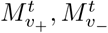 are the models with only *v*_+_, and *v*_−_ time dependent respectively (as eq. (5)), superscript *t* denoting time dependent parameter models.

#### 2.7.1 Inference of temporal dependence of mechanical parameters

We fitted five different models with a single time dependent mechanical parameter, 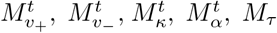, and four multiple time varying parameter models 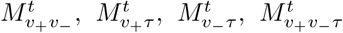, to each KT sister pair metaphase trajectory, nested models as shown in Figure 3F. We kept the same quality control criteria for trajectories as in the asymmetry model, *i*.*e*., restricted analysis to trajectories with at least 120 time points and less than 20% missing data in metaphase. We calculated the Bayes factors to determine the significance of the time dependence of mechanical forces (See Methods). We find that a substantial proportion (46.6%) of kinetochores demonstrate significant time dependence. Specifically, there is a proportion of sister pairs that show significant time dependence in *v*_−_, 20.2% (pooling the models with time dependence in *v*_−_, Table 4), *v*_+_, 15.8%, *τ*, 25.3%, *α*, 6.5% and *κ*, 1%. This suggests that the K-fibre pulling and pushing forces and the diffusive noise are time dependent. In contrast, few kinetochore pairs show significant time dependence in the centromeric spring strength *κ* or centralising force *α*, Table 4; therefore models with time dependent *κ* or *α* were not explored further. Through comparison of nested models, we determined which combinations of parameters are time dependent in each sister pair giving the model preference network, Figure 8A.

**Table 4:** Temporal dependence: Number of KT pairs that preferred each temporal model for the DMSO and nocodazole washout treated cells. In parenthesis, we quote the percentage of the sisters preferring the temporal model over the total number of sister pairs. The total number of pairs per treatment is shown in the column header.

| Preferred Model | DMSO (n=788) | Nocodazole (n=762) | Totals (n=1550) |
| --- | --- | --- | --- |
| Time invariant | 421 (53.4%) | 294 (38.6%) | 715 (46.1%) |
| $\alpha$ | 51 (6.5%) | 20 (2.6%) | 71 (4.6%) |
| $\kappa$ | 6 (1%) | 12 (1.6%) | 18/1550 (1.1%) |
| $v_-$ | 103 (13%) | 92 (12.1%) | 195 (12.6%) |
| $v_+$ | 40 (5%) | 110 (14.4%) | 150 (9.6%) |
| $v_-$, $v_+$ | 42 (5.3%) | 101 (13.3%) | 143 (9.2%) |
| $\tau$ | 93 (11.8%) | 78 (10.2%) | 171 (11%) |
| $v_- \tau$ | 15 (1.9%) | 27 (3.5%) | 42 (2.7%) |
| $v_+ \tau$ | 17 (2.1%) | 28 (3.7%) | 45 (2.9%) |
| $v_{\pm} \tau$ | 0 (0%) | 0 (0%) | 0 (0%) |

**Figure 8.**
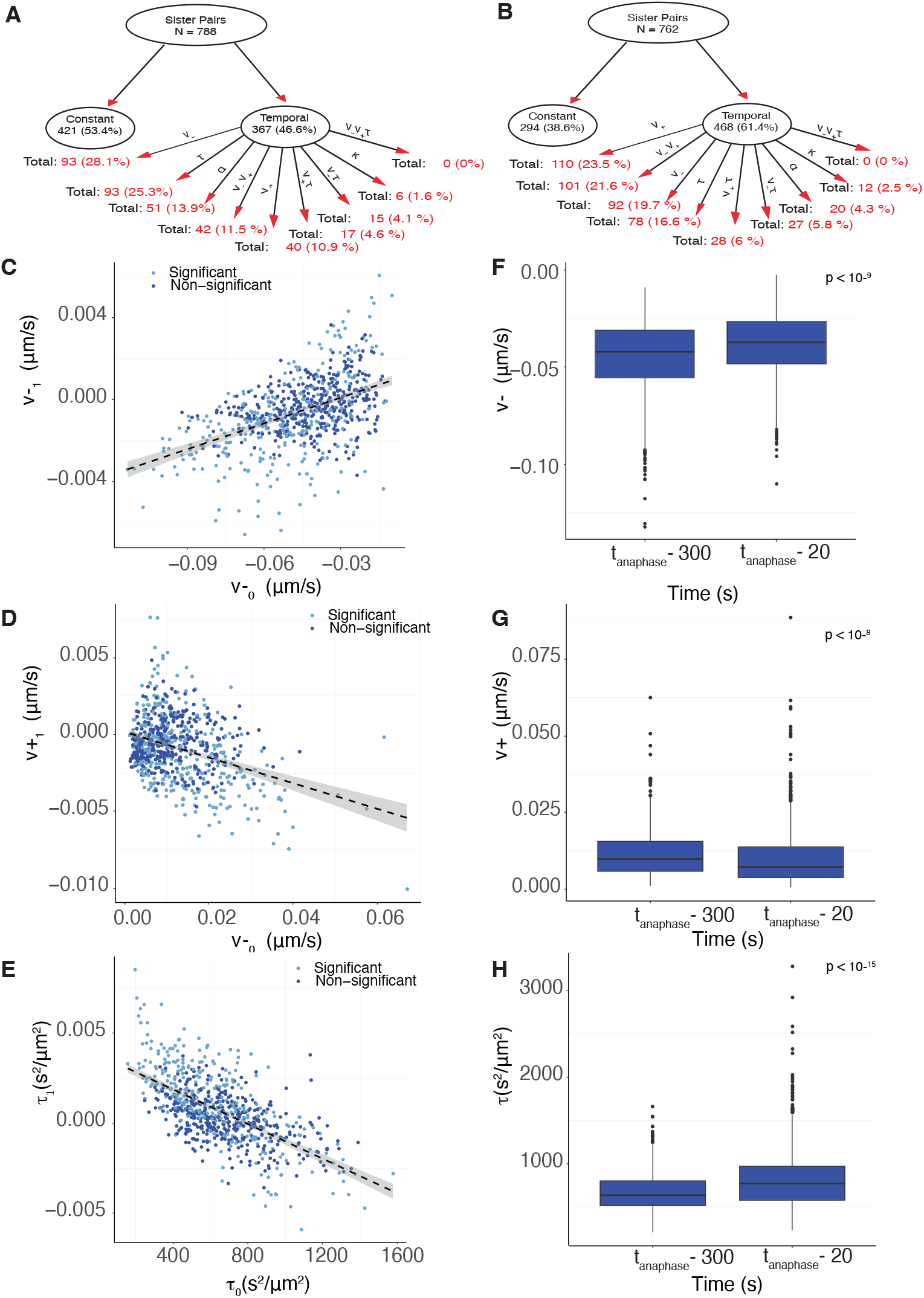
K-fiber forces decrease towards anaphase and are tuned towards an anaphase ready state. **A/B** Graphical representation of the temporal preference network with child node decomposition shown as percentage of parent DMSO (**A**) and nocodazole washout treated cells (**B**). **C/D/E** Scatter plot of posterior median of time dependence *p*_1_ versus initial parameter value *p*_0_ for **C** pushing forces, *v*_1,+_ versus *v*_0,+_; **D** pulling forces, *v*_1−,_ versus *v*_0, −_; **E** *τ*_1,+_ versus *τ*_0,+_ of DMSO treated cells. Light (dark) blue dots are KT pairs that have significant, respectively non-significant, time dependence. **F/G/H** Box-plots of posterior median parameter at the beginning and the end of the trajectory for **F** pushing forces *v*_+_; **G** pulling forces *v*; and **H** noise *τ* of DMSO treated cells. Parameters over time (mid-metaphase to late-metaphase comparison) are statistically different (*p*_*MW*_ *<* 10^−9^, *p*_*MW*_ *<* 10^−8^, *p*_*MW*_ *<* 10^−16^ for *v* _−_, *v*_+_, *τ* respectively). Finally, variances over time are statistically different with (*p*_*BF*_ *<* 10^−10^, *p*_*BF*_ *<* 0.02, *p*_*BF*_ *<* 10^−21^ for *v*_−_, *v*_+_, *τ* respectively). Parameters are inferred on the *M*_*v*± *τ*_ model.

#### 2.7.2. K-fiber dynamic parameters tune towards an anaphase ready state

To determine how the K-fibre biophysical properties vary in time, we analysed the posterior median of *v*_±,0_, *v*_±,1_ from model *M*_*v*±*τ*_ for all KTs. This model encompasses the main significant time dependence observed in our data, *i*.*e*., 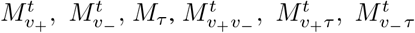 are nested models, Figure 3F. For trajectories that prefer a simpler nested model the inferred parameters were similar for the nested model, in particular the rate of change of any parameter without significant time dependence was zero in the nested model. There is significant correlation between *v* _− 1_ and *v*_− 0_, (*p*_*Corr*_ *<* 10^−16^), Figure 8C, and similarly for *v*_+_ and *τ* (*p*_*Corr*_ *<* 10^−16^ for both parameters), Figure 8D/E, with predominantly negative time dependent coefficients, 68.4 % with *v* _1_ *<* 0, and similarly 70% with *v*_+1_ *<* 0. Thus, the magnitude of both the pulling and pushing K-fibre forces typically decrease as metaphase proceeds, and K-fibers with higher magnitude pulling (or pushing) forces decrease faster, whilst the pulling, respectively pushing forces, of K-fibers with weak pulling, pushing are strengthened. This suggests the forces are being focused towards a specific value, *i*.*e*., K-fiber forces tune towards *an anaphase ready state*. This anaphase ready state corresponds to the time invariant K-fibers, *i*.*e*., when *v*_1−_ = *v*_1+_ = 0, giving *v*_−*\**_ = 0.033 microns s^−1^(*sd* = 0.009), *v*_+*_ = 0.003 microns s^−1^(*sd* = 0.001) using the line of regression, Table 5. Tuning is observed in the majority of cells, for instance the same positive correlation between *v* _−0_ and *v* _−1_ is present in 26 out of 26 individual cells, with the majority (24 of 26) having a negative (and thus physical) intercept on the *v* _−1_ = 0 axis, Appendix Figure B14. Tuning is also uniform across the MPP, Figure B13. The anaphase ready state has lower energy than at mid-metaphase, *i*.*e*.,, the forces from mid-metaphase reduce significantly over the 5 mins to anaphase, Figure 8F,G. This is consistent with the typically negative rate of change parameters, Figure 8. The KT pairs with significant time dependence are in fact those with initial values further from the anaphase ready state, Table 5, and thus have a larger rate of change, and a stronger time dependence. Their time dependence is therefore easier to detect.

**Table 5:** KT pairs with significant time dependence are those with initial values further from the anaphase ready state. Table shows the posterior median estimates of *v* _± 0_ and *v*_±_ for the the significant/non-significant time dependent KT pairs. Posterior distributions are statistically different, using one sided Kolmogorov-Smirnov (one-sided Mann-Whitney) tests. Anaphase ready state estimates from the line of regression of *v*_± 0_ against *v*_±_ are shown for comparison.

|  | Posterior Medians |  | Anaphase ready state | p-value |
| --- | --- | --- | --- | --- |
|  | Time dependent KTs | Time invariant KTs |  |  |
| $v_{-0}$ | -0.052 | -0.041 | $v_{-*} = -0.033$ | $< 10^{-8}$ ($< 10^{-7}$) |
| $v_{+0}$ | 0.012 | 0.009 | $v_{+*} = 0.003$ | $< 10^{-4}$ ($10^{-7}$) |

To further explore tuning, we estimated for each KT pair the forces *v* _±_(*t*) at 2 fixed reference time points, specifically at 300s prior to anaphase (in mid-metaphase) and 20s before anaphase onset (in late metaphase). The scatter plot, Appendix Figure B11, clearly shows the focusing effect, with a fixed point of −*v*_−_ = 0.04 microns/s (with standard error (se) = 0.016 microns/s), *v*_+_ = 0.01 microns/s (se = 0.009 microns/s), *i*.*e*., similar estimates for the K-fiber pulling, pushing strengths of the anaphase ready state as above. These results suggest the following temporal dynamics of *v*_−_,

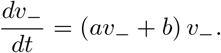

Specifically, Figure 8C implies that the average rate of change of log |*v*_−_|, *v*_1−_, is correlated with *v*_0−_, giving *a, b >* 0. This system has a (stable) fixed point at *v*_−_ = −*b/a*; it integrates to give the general time dependent solution:

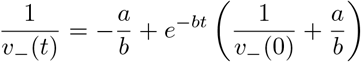

Turning this into a discrete map, *v*_−_(0) to *v*_−_(*t*) for a fixed *t*, as Appendix Figure B11, gives a stable fixed point of this map at −*b/a*.

To analyse time dependence in the diffusive noise, the posterior median of *τ*_0_ and *τ*_1_ from *M*_*v* ±*τ*_ were analysed. 125 out of 788 DMSO treated kinetochore pairs (15.9%) show significant time dependence in the noise (preferring any of 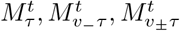). The noise precision of most of the kinetochore pairs (64.1%) increases which indicates that random (diffusive) noise in KT dynamics decreases over metaphase. There is a significant correlation (negative), (*p*_*Corr*_ *<* 10^−15^), between the time dependent coefficient (*τ*_1_) and the initial magnitude of noise (*τ*_0_), which suggests, for KT pairs with large noise at the beginning of the movie, that their noise typically diminishes during metaphase. Similarly, precision significantly increases from mid to late metaphase at the population level, (*p*_*MW/KS*_ *<* 10^−16^), Figure 8H. As shown Appendix Figure B11C, the noise precision is also tuned, *i*.*e*., the anaphase ready state has an associated level of diffusive noise.

### 2.8 KT heterogeneity and trends are robust to perturbation of the spindle assembly pathway

To determine to what extent KT heterogeneity is sensitive to congression dynamics and conditions where segregation is more error prone, we disassembled MTs with nocadozole. The spindle reforms upon washout with microtubule nucleation at both centrosomes and at KTs, [Tulu et al., 2006], and thus spindle assembly dynamics is different, has different initial conditions and possibly utilises a different pathway. Chromosomes congress and segregate on these spindles; however segregation errors due to merotelic attachment are more frequent, [Cimini et al., 2001, Worrall et al., 2018b], and there is a 3 fold increase in lazy kinetochores [Sen et al., 2021]. Lazy KTs are defined as KTs that significantly fall behind their cluster at any point during anaphase. At 4s time resolution, there were 0.26% lazy KTs in RPE1 cells, rising to 0.86% in nocodazole washout treated cells, Sen et al. [2021]. It is unknown which of the mechanisms that result in the normally high fidelity of segregation are disrupted under nocodazole washout treatment.

We analysed KT heterogeneity in metaphase of nocodazole washout treated cells (330nM conc, 2 hr treatment followed by washout as in Sen et al. [2021]), analysing 1240 KTs in 33 cells (out of 51 imaged cells using identical cell filtering requirements to previously). In these 33 cells detection and tracking performance was similar to cells cultured in DMSO; on average we obtained 89.9 KTs per frame, and 37.6 sister pairs per cell (quartiles Q1=35, Q3=41) where both sisters were tracked for at least 200 seconds (100 frames). The videos of nocodazole washout treated cells were typically longer than those of DMSO cultured cells, 237 frames versus 181 frames.

Metaphase was similar in nocodazole washout treated cells to DMSO cultured cells; the quasi-periodic oscillations had similar period and strength (ACF depth), with only weak variation across the MPP, Figure 9, and did not vary over the duration of metaphase, Appendix Figure B3. MPP maturation over the last 5 mins was substantially weaker in nocodazole washout although still significant, with weak reduction in KK and the MPP width over the last 5 mins of metaphase. This is possibly because the MPP width had already decreased by 5 mins before anaphase, Appendix Figure B3, which contrasts to the steady reduction seen in DMSO Figure 2. The average KK distance is marginally larger in nocodazole washout treated cells (1.135 microns compared to 1.104 microns, averaged over time and KTs), and there is a stronger spatial dependence across the MPP in nocodazole washout treated cells with a stronger increase in the KK distance towards the periphery, Figure 9A. The period and oscillation strength (ACF depth) were approximately constant across the MPP in both DMSO and nocodazole washout; the 20.2% greater depth of the ACF in the MPP centre compared to the periphery (DMSO) reflects the fact that oscillations are noisier towards the periphery.

**Figure 9.**
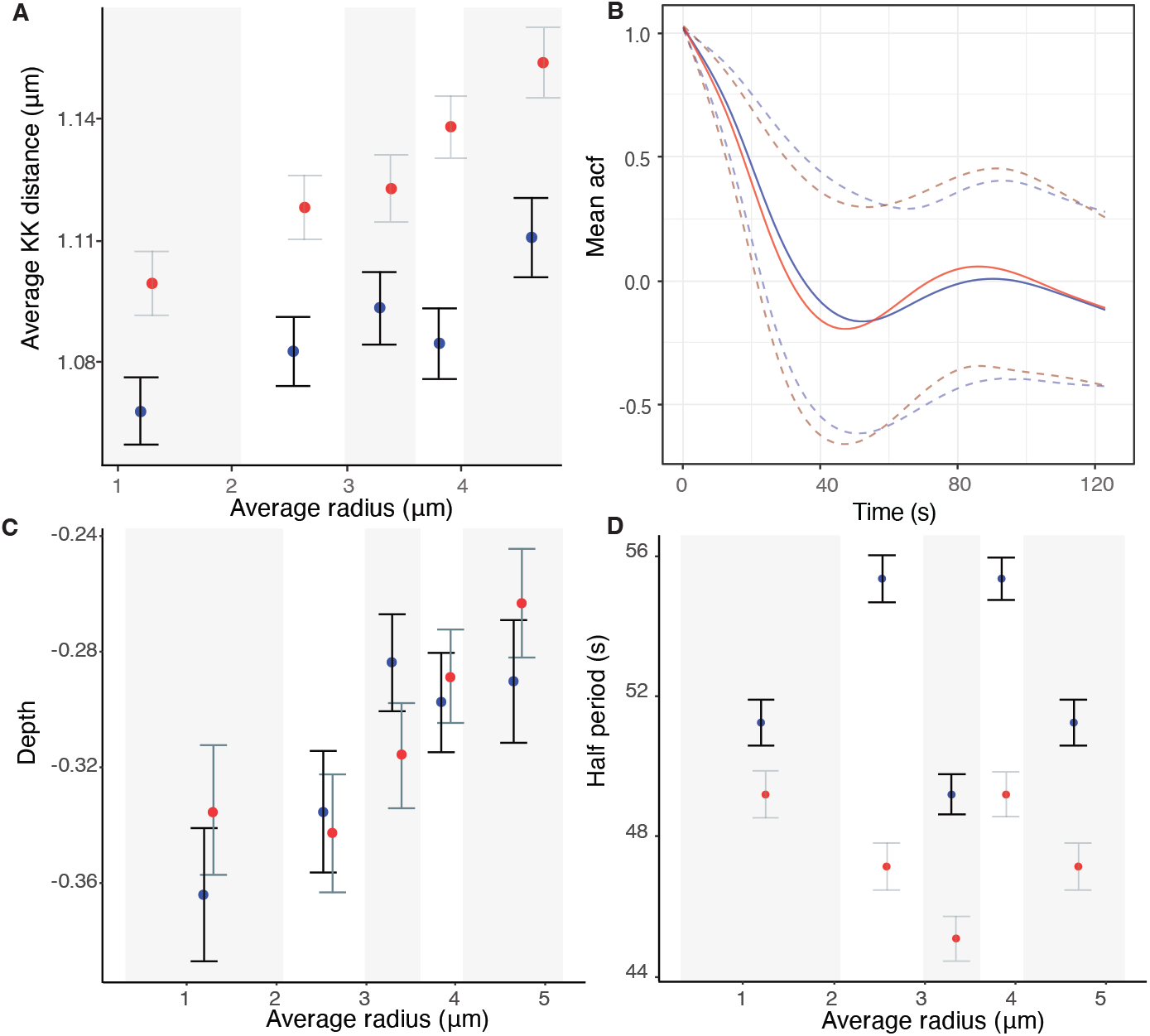
Quasi-periodic oscillation period and strength vary weakly across the MPP and are robust to perturbation. **A** Sister-sister separation (KK distance) with position in the metaphase plate. **B** Mean ACF for DMSO (blue) and nocodazole washout (red) treated cells. Dashed lines denote the 5% and 95% percentile. **C** Average depth (ACF minimum) and **D** oscillation period for sister pairs across the metaphase plate. ACFs for each spatial partition are given in Figure B25.

We next determined if the heterogeneity in the biophysical parameters in time and space observed in DMSO are present in nocodazole washout treated cells. Although most biophysical parameters were substantially changed in nocodazole washout treatment, heterogeneity trends were all present and essentially identical,

1. **Biophysical parameters**. Nocodazole washout treated cells have on average weaker K-fibers, specifically the magnitude of *v*_+_, *v*_−_ were reduced by 37.5% (*p*_*MW*_ *<* 10^−15^), and 5.8% (*p*_*MW*_ *<* 10^−4^) respectively, Appendix Table A7. They also have significantly reduced diffusive noise, (*p*_*MW*_ *<* 10^−15^, precision increased by 20%), a higher centralising force (*p*_*MW*_ *<* 10^−10^, *α* increased by 15%) and the spring constant is reduced by 40% (*p*_*MW*_ *<* 10^−9^). The natural length is as expected the same as in DMSO, since the same informative prior is used. These data suggests that fewer MTs are bundled into K-fibers, leaving a higher density of spindle MTs that would explain the increase in the strength of the centralising force.
2. **Spatial trends with the metaphase plate radius** were reproduced similar to DMSO, Figures 10 and B9. In nocodazole washout treated cells, K-fibre pushing (*v*_+_), pulling forces (*v*_−_), the spring constant (*κ*) and diffusive noise (*τ*) were reduced by 14%, 34%, 65%, 19% from middle to periphery of the metaphase plate (in nocodazole washout treated cells). Finally, unlike DMSO, we observed a statistically significant reduction of 15% of the centralising force parameter *α* towards the periphery, (*p*_*Corr*_, *p*_*Kendall/Spearman*_ *<* 10^−3^).
3. **Sister asymmetry** was similar with 28.5% having significant asymmetry, Figure 5B. There was an identical anti-correlation between the strength of pulling and pushing forces, Figure 5, and Appendix Figure B15, and sister differences were identical to differences between random KTs. The lateral organisation transverse to the MPP into strong/weak asymmetric and symmetric KT groups was also reproduced, Figure 6B,D,F,H.
4. **Temporal trends** were very similar, Figure 8B, Table 4, in fact there was a reduction in the preference for the time invariant model (53.4% DMSO to 46.1% nocodazole washout), likely due to the typically longer time series and the reduced diffusive noise under nocodazole washout, both increasing the ability to detect temporal variation. Significant temporal dependence was primarily seen in *v*_±_, *τ* as before, with preferences for temporal dependence in *v* _−_(24.5%), *v*_+_ (21.7%), *τ* (16.6%), whilst preference for temporal dependence in the other parameters is infrequent, Table 4. The correlation between the rate parameters and the initial parameter value was similar, Appendix Figure B16 suggesting that K-fibers in nocodazole washout treated cells also tune towards an anaphase ready state. The anaphase ready state has slightly lower pulling force (magnitude) and pushing forces, (−0.02, 0.003) compared to (−0.04, 0.01) microns/s in DMSO, with this difference being statistically different (*p*_*MW/Ztest*_ *<* 10^−20^). As before the K-fibre reduction in pulling/pushing force magnitudes over the last 5 mins of metaphase is significant (*p*_*MW/KS*_ *<* 10^−16^, *p*_*MW/KS*_ *<* 10^−9^ for *v*_−_, *v*_+_ respectively). There is little increase in the precision over that period, Appendix Figure B12 and Appendix Figure B16, whereas the precision increased in DMSO, Figure 8. This is possibly explained by the higher precision on all inferred models in nocodazole washout treated cells, Figure 10, Appendix Figure B12 and Appendix Table A4, possibly limiting any further increase. We confirmed that this higher precision isn’t due to differences in movie length (reducing the length of movies had little effect on mean (median) estimates) as expected.

**Figure 10.**
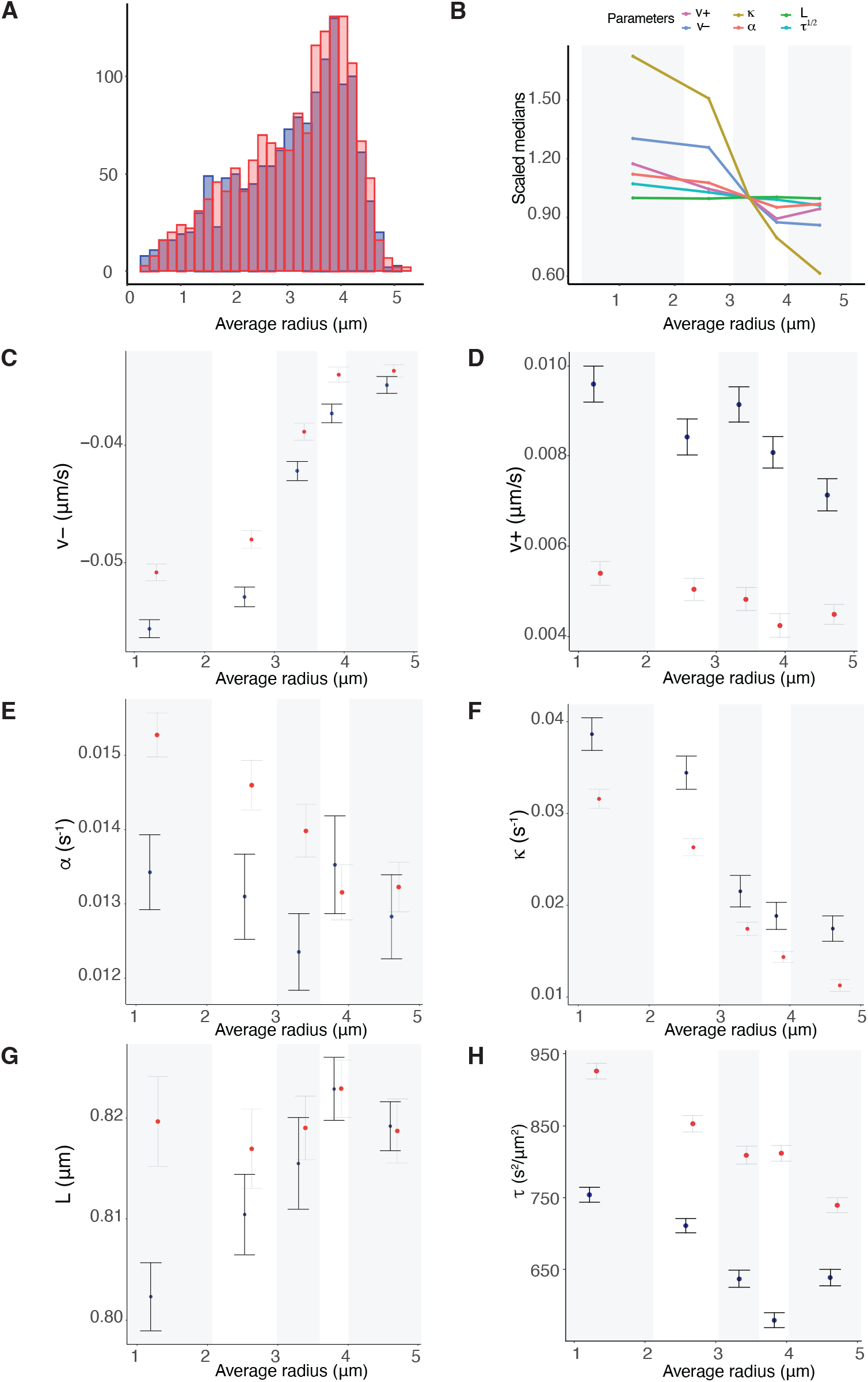
Comparison of spatial mechanical parameter trends across the metaphase plate for DMSO (blue) and nocodazole washout (red) treated cells. **A** Location of DMSO (blue) and nocodazole washout (red) treated kinetochore pairs within the metaphase plate, based on 798 and 984 kinetochore pairs respectively, over 59 cells. **B-F** Summary statistics of indicated biophysical parameters partitioned into 5 groups for the average radial position within the metaphase plate. Groups are equally sized, containing the same number of observations (DMSO and nocodazole washout treated cells), with the five groups defined by their radial distances, *i*.*e*., [0, 2.07], (2.07, 2.97], (2.97, 3.59], (3.59, 4.01], (4.01, 5.20] respectively. Hence, the groups’ margins are not identical to groups of Figure 7. **B** Summary of scaled spatial trends over all parameters for nocodazole washout treated cells, in reference to Figure 7. **C** Pulling forces *v*_−_, **D** Pushing forces *v*_+_, **E** Centralising force, **F** Spring constant *κ*, **G** natural length of the centromeric chromatin spring connecting the kinetochore sisters *L*, **H** precision *τ*. Spatial trends for coherent and incoherent probabilities (*p*_*coh*_, *p*_*incoh*_) can be found in Appendix Figure B9.

The heterogeneity in space, time and through random variation is thus robust to variation in spindle assembly dynamics, even under conditions with a higher chromosome mis-segregation rate, suggesting that this heterogeneity is intrinsic to mitosis, either tolerated or through design.

### 2.9 Oscillations are classified into 5 clusters varying in quality and strength and represent a distinct type of heterogeneity

Finally, we analysed the most obvious aspect of within cell variability – the fact that typical cells have a mixture of both good and poor oscillating KT pairs. Metaphase oscillations are robust to variation in spindle assembly, and are a result of K-fiber switching control processes [Civelekoglu-Scholey and Cimini, 2014, Banigan et al., 2015] and thus dependent on biophysical parameters. Thus, we address if the biophysical heterogeneity we documented above explains the variation in oscillation quality and strength, or if it represents another aspect of heterogeniety.

To analyse metaphase oscillation variability we clustered KT pair trajectories based in the autocorrelation function (ACF) of the sister mid-point (mean of the sister *x* coordinates) using hierarchical clustering with time warping, [Murtagh and Contreras, 2012], (implemented in the “ tsclust” R package Montero and Vilar [2014]). We pooled the DMSO and nocodazole washout treated cell datasets to enable comparisons between treatments. 5 clusters were deemed most appropriate, Figure 11A based on the clustering dendrogram, Figure 11B, with cluster signatures that were fairly consistent across different clustering methods. We confirmed that there was consistency in the ACF of individual pairs within a cluster, Figure 11E-I. The clusters had distinct average ACFs, Figure 11A, suggesting the following 5 phenotypes: *strong oscillators with short period* (ACF local maximum 89 s), *strong oscillators with longer period* (ACF local maximum 116 s), *weak oscillators with long period* (1*/*2 period of 60s), *weak and noisy oscillators* (1*/*2 period of 66s) and finally, *poor oscillators/non-oscillators*. Examples of trajectories in each category are shown in Figure B19. The oscillatory category proportions are similar in DMSO and nocodazole washout treated cells Figure 11C, with a slightly higher proportion of poor oscillators in DMSO treated cells, *i*.*e*., 35.8%, compared to 27.3% in nocodazole treated cells, Table A8 and Figure 11C. Cells varied substantially in the proportion of poor oscillators, Figure 11D, with 13.63% (6.45%) to 63.16% (46.88%) in DMSO (nocodazole washout) treated cells. Thus, some cells have poorer KT population oscillations than others, although each cell had a mixture of oscillators, Figures B21 and B22.

**Figure 11.**
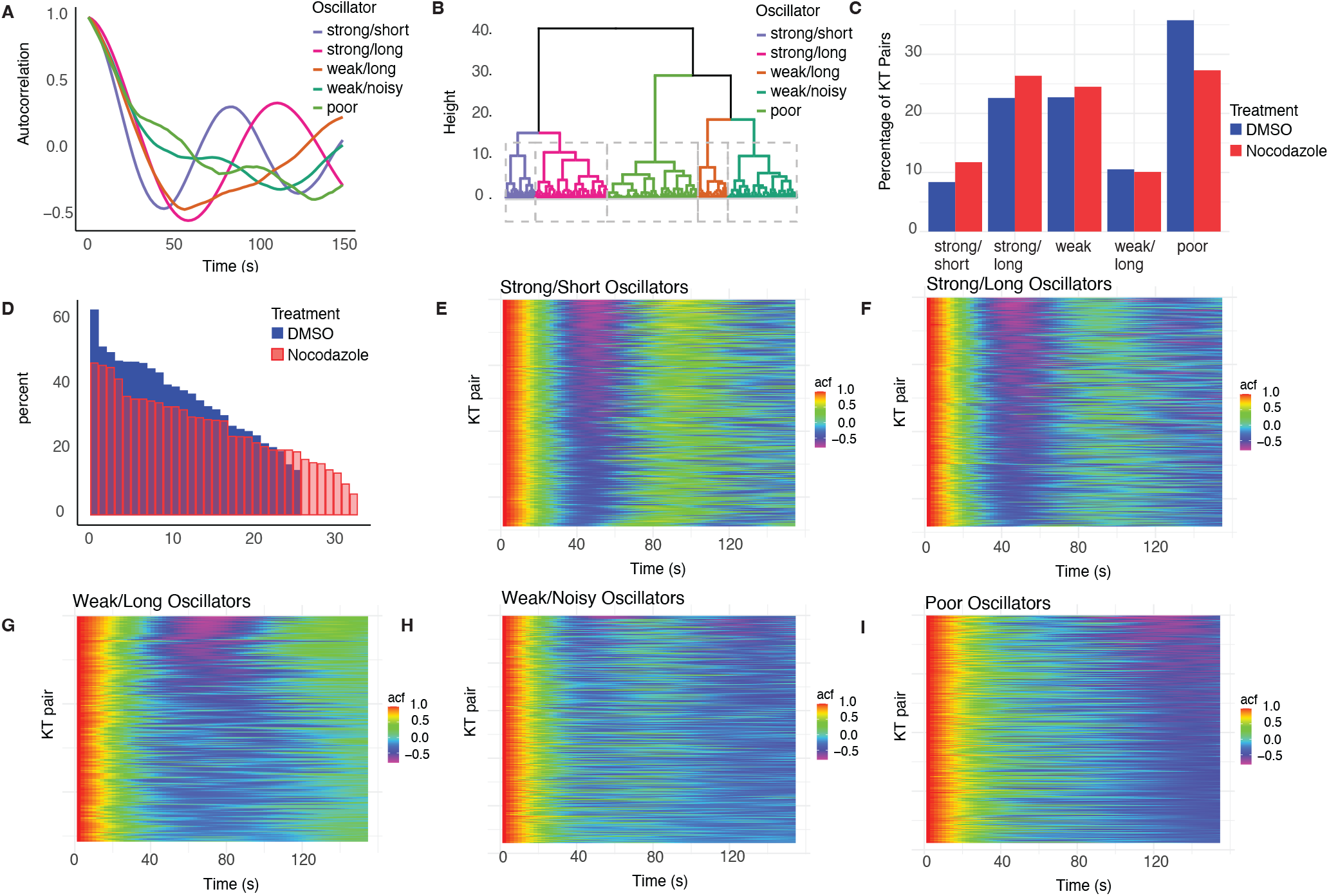
Oscillator classification into 5 clusters-strong oscillators with short period, strong oscillators with long period, weak oscillators with very long period, weak and noisy oscillators and poor/no oscillators. **A** Centroid ACF plots for each of the 5 clusters. Typical trajectories are plotted in Figure B19.**B** Clustering dendogram (tree plot) which displays the hierarchy of clusters created by hierarchical clustering. It shows which clusters were merged at what distance/similarity level (height). **C** Cluster proportions for DMSO (blue) and nocodazole washout (red) treated KT-pairs dataset. **D** Proportion of poor oscillators in each cell, for DMSO (blue) and nocodazole washout treatment (red) datasets (cells ordered by decreasing proportion of poor oscillators). **E-I** ACF of individual KT-pairs in each cluster, shown in pseudocolour. KT pairs stacked by row.

To determine whether there are mechanical differences between our oscillation categories, we analysed the biophysical parameters of sister pairs in each cluster. Strong oscillators have higher K-fiber pulling forces, higher |*v*_−_|, and directional switching events are longer (higher *p*_*incoh*_) compared to weak and poor oscillators, Figures 12 and B24; the median parameter estimations of the asymmetric 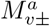 model were statistically significant different between the strong versus poor oscillators in most parameters, *v*_−_, *p*_*incoh*_ the strongest, Table A9. There is a marginal decrease in the strength of the centromeric spring (*κ*) and in the strength of the centralising forces from strong to poor oscillators. This contrasts with oscillation amplitude that is reported to be dependent on the PEF, Ke et al. 2009]. The period of the strong oscillator clusters is reflected in a higher probability of remaining in the coherent state in the long period cluster, *i*.*e*., *p*_*coh*_ is higher.

**Figure 12.**
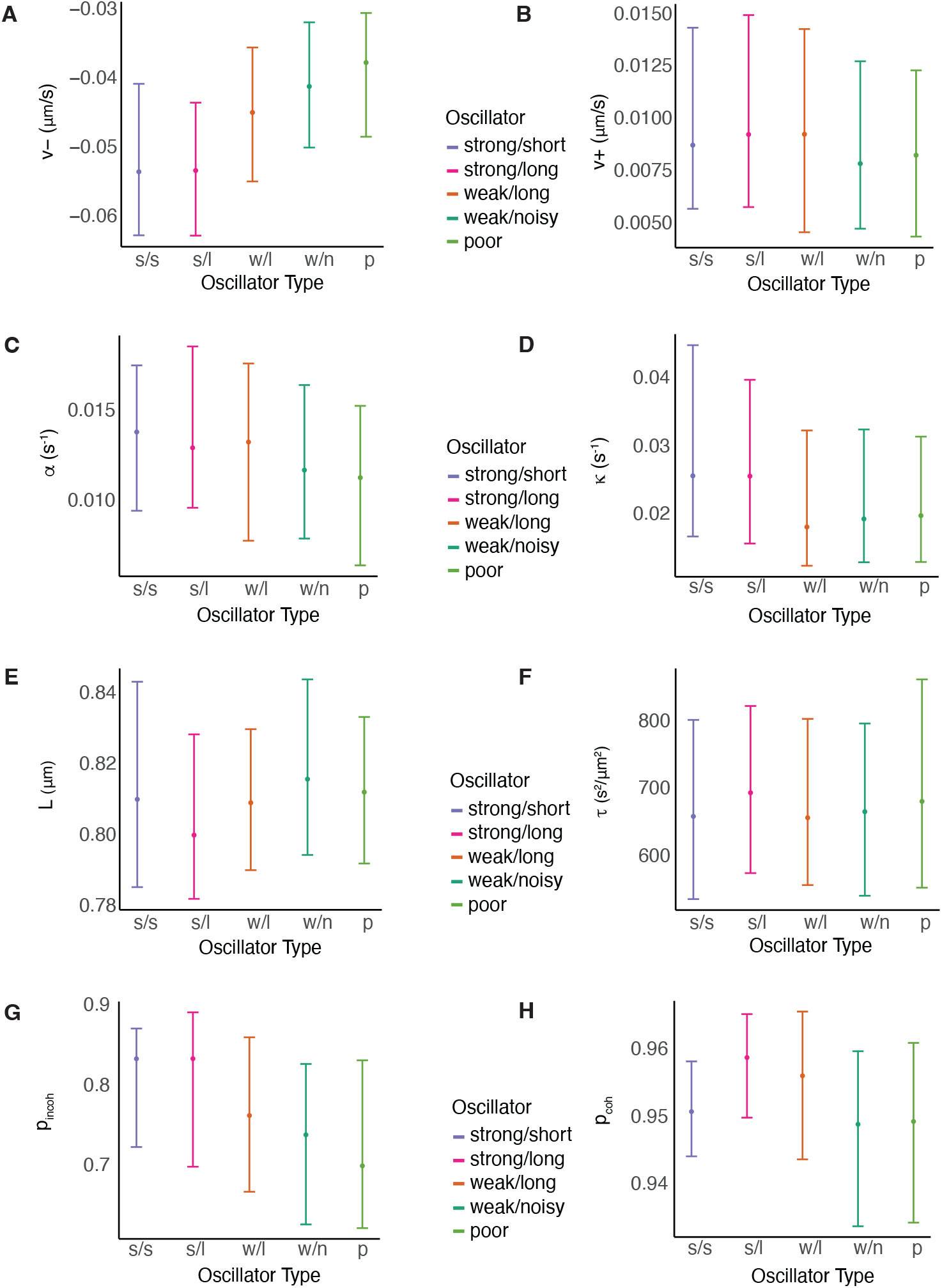
Biophysical parameters with oscillation quality (DMSO). Parameter estimates for the asymmetric 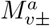 model, are consistent across different types of oscillators in nocodazole washout treated cells, except *v*_−_ and the probability of remaining incoherent between frames (*p*_*incoh*_), where we observe a decreasing trend as the quality of oscillation deteriorates. Correponding data for nocodazole washout treated cells is in Appendix Figure B21.

Oscillator type profile changed across the MPP, with the proportion of strong oscillators decreasing towards the periphery, Appendix Table A8, Figure B20, as previously reported in other cell types [Civelekoglu-Scholey et al., 2013a, Armond et al., 2015a]. This spatial trend may explain the change in biophysical parameters from strong to poor oscillators. We partitioned the KT pairs into 5 radial partitions, Figure B23, and analysed *p*_*incoh*_ and *v*_−_ variation by partition and cluster. This demonstrated that indeed the pulling forces and the directional switching time decreased towards the periphery in each cluster, Figure 10, reflecting trends within the MPP of the KT population as a whole Figures 10 and B9. However, within each spatial partition the strong oscillators have stronger pulling forces and longer directional switching times (higher *p*_*incoh*_); in fact the directional switching time was significantly different in most partitions for both DMSO and nocodazole washout treatments, Appendix Table A10. An ANOVA analysis on *p*_*incoh*_ with oscillator cluster (poor and strong only) and spatial partition indicates that 20% of variation is attributed to *r*, and 10% to oscillation, with both radial distance and oscillations being significant, (*p* = 10^−16^). Similarly, for *v* _−_16% is attributed to oscillation and 18.4% is attributed to *r*, (*p* = 10^−16^). There was no correlation between oscillation quality and asymmetry - the fraction of (significant) asymmetric pairs was not significantly different for each oscillation category, *p*_*CMH*_ = 0.41 for DMSO treated cells and *p*_*CMH*_ = 0.10 for nocodazole washout treated data (test accounts for the distance from the centre of the metaphase plate). Thus, oscillation quality variability is distinct from heterogeneity in the biophysical parameters.

## 3 Discussion

In this work we developed a data driven modeling framework for the study and analysis of KT dynamics at the cell level, revealing substantial heterogeneity in KT dynamics within a cell, Figure 13. Specifically, we observed both temporal organisation, with maturation of KT dynamics in time (*v* _±_, *τ*), thereby tuning system dynamics towards an anaphase ready state, and spatial organisation, with the MPP organised both within the MPP (*v*_±_, *κ, τ*) and transverse to the MPP (caused by random sister asymmetry in *v*_−_). Metaphase oscillations are also heterogeneous, in part reflecting biophysical parameter trends within the MPP, but also representing a 4^*th*^ dimension of heterogeneity through noise in directional switching control. Perturbing the spindle self-assembly pathway (nocodazole washout) gave similar results indicating that the processes governing heterogeneity and spatial-temporal trends are intrinsic to mitosis.

**Figure 13.**
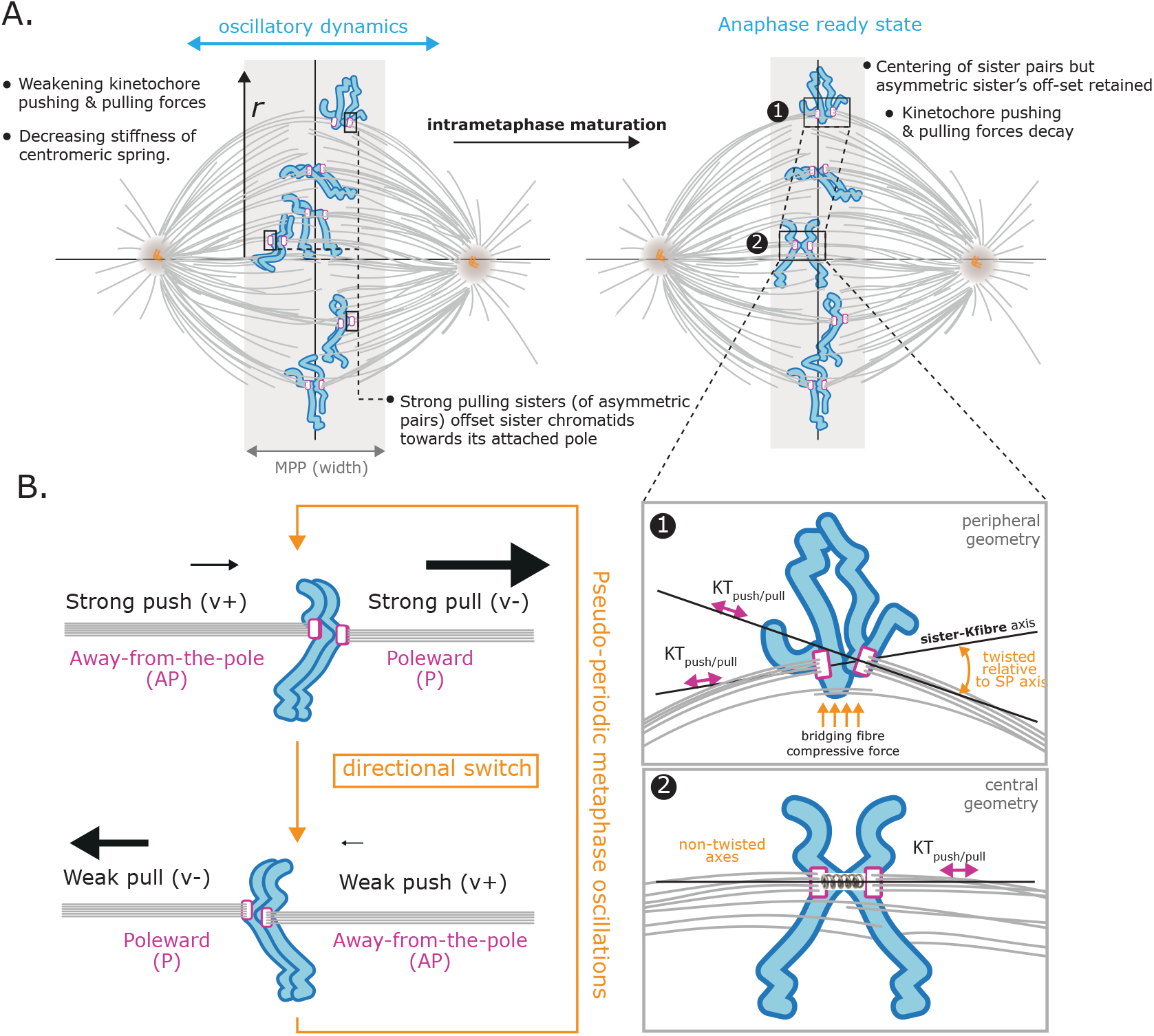
Organisation of the KT population within (radially) and transverse (laterally) to the MPP. **A** Left. Schematic of the spindle showing microtubules, bridging fibers, KTs (magenta) and chromosome arms (cyan). K-fiber pulling and pushing forces decay with radial distance (*r*), while the centromeric spring stiffness decreases. Strong pulling sisters of asymmetric pairs offset the sister pair towards its pole. Right. Late metaphase organisation after intrametaphase temporal tuning. The anaphase ready state has more centred chromosomes, and the K-fibre pulling and pushing forces typically decay. Sister chromatid offset is retained, but decreased. Annotated chromosomes at the periphery, (1), and central, (2), are shown in detail (lower panel), emphasising how the different local spindle geometry impacts attachment force orientation. **B** Schematic representation of forces in an asymmetric sister pair through the directional switching cycle (orange) of metaphase oscillations. Magnitude of the force is indicated by size of arrow. Schematic shows the inverse correlation between pulling and pushing strength of a K-fibre, whilst pulling forces are substantially larger than pushing forces.

The majority of the parameter variation we report can be related to the K-fibers, Figure 13. Specifically, we observed (i) strong trends in the K-fiber force strength across the MPP, with both the pulling and pushing forces decreasing towards the periphery (38% and 25% respectively), (ii) asymmetry in the sister K-fiber pulling forces that results in transverse KT organisation across the MPP, and (iii) a decay of the pulling and pushing forces in the last 5 mins of metaphase. MT polymerisation in K-fibers will likely contribute substantial active noise, [Gnesotto et al., 2018], thus the spatial and temporal variation in diffusive noise probably reflects changes in MT dynamics and/or cohesion of K-fibers, rather than a change in chromosome diffusion coefficients. What causes this spatial and temporal variation in K-fiber dynamics is unknown. We hypothesise that the observed temporal intrametaphase changes in K-fiber dynamics are due to regulatory processes that prepares the system for anaphase, primarily reducing the (kinetic) energy in the system. Intrametaphase molecular changes in the KT or K-fibers that could underpin this process have not been reported, more likely reflecting the fact that quantification is difficult given the need to accurately time metaphase progress on a time scale of minutes with a resolution of seconds. In contrast, differences between prometaphase and metaphase have been reported, Harasymiw et al. [2019], Embacher et al. [2022]. As to spatial heterogeneity, there are known mechanisms that could generate such trends, specifically,

- The spindle is not a homogeneous environment with an increase of K-fiber and bridging fiber curvature towards the periphery, [Vukušić et al., 2017, Kajtez et al., 2016, Miles et al., 2022], thereby changing the geometry of the KT/K-fiber interface, whilst bridging fibers are also under much higher compressive forces at the periphery compared to central K-fibers, [Pavin and Tolić, 2020]. The spatial dependence of the K-fiber properties (*v*_±_, *τ*) within the MPP may therefore be an adaptation to this changing geometry.
- Chromosome size and centromere size varies between chromosomes, and in particular, large chromosomes tend to be localised towards the periphery, [Mosgöller et al., 1991, Booth et al., 2016]. Both centromere differences between chromosomes [Dumont et al., 2020] and KT size [Drpic et al., 2018] have been implicated in biasing chromosome segregation errors, whilst chromosomes 1 and 2 (the 2 largest chromosomes) have a higher mis-segregation rate [Worrall et al., 2018b]; these biases may be a result of a mechanical dependence on size and thus related to the within MPP trends we observe.
- Our asymmetric models indicate that random variation is responsible for KT heterogeneity within a cell, particularly in *v*_±_, *τ*, with variation in *v*_−_ between sisters giving rise to lateral organisation of KTs in the MPP, Figure 13. This random variation in K-fiber properties could be a result of outer KT assembly at nuclear envelope breakdown (NEBD), [Navarro and Cheeseman, 2021], or self-assembly of the K-fibers and spindle during prometaphase, which will introduce variation in K-fiber composition, bundle coherence, [Armond et al., 2015b], retrograde flux and/or control processes. Since there is an inherent anti-correlation in K-fibre pulling and pushing forces, Figure 5B, bundle size (MT count) is unlikely to be responsible as this would be expected to increase both pushing and pulling forces. Variation in K-fiber flux has the correct signature, *i*.*e*., a decrease in flux increases pushing forces and decreases pulling forces. Further investigation is needed to determine both the cause of asymmetry and MT bundle dynamics generally, and if the associated lateral organisation of the MPP has consequences for segregation efficacy for affected chromosomes.

We also addressed if the spatial organisation in the K-fiber pulling forces in metaphase is preserved into anaphase. We fitted a metaphase-anaphase model to the trajectories, Appendix D and compared the K-fiber pulling force in metaphase with that in anaphase. There is a weak correlation between *v*_−_ and the anaphase speed, *ρ* = 0.174, *p*_*Corr*_ *<* 10^−15^, Appendix Figure D3F, whilst spatial trends in the pulling forces within the MPP are lost in anaphase, Appendix Figure D3C, reminiscent of the anaphase speed governor, [Anjur-Dietrich et al., 2021]. In fact, the anaphase speed of a KT pair had a higher correlation with its anaphase onset time, *ρ*_*DMSO*_ = 0.247 (*p*_*Corr*_ *<* 10^−9^) in DMSO, *ρ*_*noc*_ = 0.125 (*p*_*Corr*_ *<* 10^−3^) in nocodazole washout, (with *ρ*_*noc*_ *< ρ*_*DMSO*_, *p*_*Ztest*_ = 0.004). The anaphase speeds were similar in DMSO and nocodazole washout (*p*_*MW*_ = 0.083), which contrasts to the lower pulling speed in nocodazole washout *p*_*MW*_ *<* 10^−4^; pulling speed |*v*_−_| decreased 5.85% and anaphase speed decreased 2.8% in nocodazole. This change in heterogeneity dependence likely reflects the fact that K-fibers are regulated by different mechanisms and the KTs are in different states in metaphase and anaphase, KTs undergoing substantial phosphorylation at anaphase onset [Su et al., 2016, Vázquez-Novelle et al., 2014].

We observed spatial trends in the centromeric spring constant *κ*, the stiffness decreasing 53% from the centre to the periphery of the MPP. Our model assumed a linear spring, although nonlinearity has been reported, stiffness increasing with extension, [Stephens et al., 2013, Harasymiw et al., 2019]. However, since the spring extension increases towards the periphery, Figures 9 and 13, nonlinearity is not the cause of trends observed in the spring constant with radial distance *r*, in fact it would give an opposite trend. Centromere size could potentially affect the spring stiffness but data on centromere size variation within the MPP isn’t available, and the change in *κ* over the MPP is substantial suggesting it arises from a direct dependence on *r*, for instance from the change in the spindle geometry. The inferred spring constant in all our models, and in other studies such as Harasymiw et al. [2019], is an effective spring constant, so this must be considered as a possible cause. The centromeric spring models the connecting chromatic material between the sister chromatids, a composite material with likely anisotropic elasticity. There are two clear effects that might affect the effective spring constant upon moving towards the periphery. Firstly, a geometrical effect, with K-fibers not being aligned with the spindle axis, Figure 13; increasing the sister-sister distance would then also comprise a shear and torque. Secondly, chromosome arms take up a different geometry towards the periphery, being pushed outwards relative to the KTs, thereby generating a torque. How this anisotropy in the centromeric spring affects the effective spring constant is unknown.

We identified two sources of random variation that impacts KT dynamics, random variation in K-fiber strength and random variation in directional switching parameters, the latter giving rise to variation in oscillator quality. Random variation in sister K-fiber strength, which was identical to differences between random KTs, resulted in lateral organisation of the MPP. In fact KT clusters that descend to respective poles in anaphase are organised laterally with weak pulling KTs on the inside, and strong pulling KTs on the outside, Figure 6, Figure 13. Crucially, this ordering was preserved from mid to late metaphase in both DMSO and nocodazole washout, Appendix Figure B17D,E, Appendix Figure B18D,E, suggesting that the tuning towards the anaphase ready state does not affect poleward bias of these groups. Tuning was apparent in the descending clusters, with a statistically significant reduction in spread from mid to late metaphase, *i*.*e*., 0.468 to 0.414 microns in DMSO and from 0.351 to 0.321 in nocodazole washout (*p*_*MW*_ *<* 10^−15^, one-sided for both treatments). This lateral organisation is also weakly preserved through to anaphase, Appendix Figure B17F, Appendix Figure B18F. Oscillations are also heterogeneous within cells, section 2.9, with the proportion of poor oscillators varying substantially between cells. Strong oscillators have higher K-fiber pulling forces, higher |*v* _−_|, and directional switching events take longer (higher *p*_*incoh*_) compared to weak and poor oscillators, Figures 12 and B24. As previously reported, Civelekoglu-Scholey et al. [2013b], Armond et al. [2015a], oscillation quality deteriorates towards the MPP periphery, Appendix Figure B20, and thus although these trends partially mirror that of location in the MPP, Appendix Figure B9, Figure B23A,C, they are not fully explained by location. Thus, oscillation quality is another dimension to KT heterogeneity, distinct from spatial and temporal trends, and sister asymmetry.

Our analysis suggests that metaphase dynamics are robust to changes in the spindle environment. Specifically, despite the substantial variation in K-fibre length and composition across the MPP, Novak et al. [2018], qualitatively similar dynamics occurs throughout the plate indicating mechanism robustness, in fact the strength of oscillations and the period are essentially invariant across the MPP, Figure 9. This robustness is of course already well documented; quasi-periodic oscillations being observed in different human cell lines, [Jaqaman et al., 2010, Armond et al., 2015a], are robust to a range of perturbations, including decreasing flux [Guerreiro et al., 2021] and perturbed K-fiber polymerisation [Jaqaman et al., 2010], and conserved across species. We examined robustness to spindle assembly dynamics using nocodazole washout treatment; oscillation quality was similar Figure 11C and we saw practically identical trends in biophysical parameters in space (transverse and within the MPP) and in time (intrametaphase tuning), both effects also being observed in individual cells, Appendix Figure B15. Of note in nocodazole washout, centralising forces were stronger, K-fibers weaker (lower |*v*_±_|), diffusive noise reduced and the anaphase ready state had lower energy (with lower |*v*_±*_|). We hypothesise that K-fiber bundle is smaller in nocodazole washout, with fewer MTs, so are weaker but more coherent, thus reducing active noise contributions to the inferred diffusion. Correspondingly the density of MTs is higher in the spindle explaining the higher mid-point centralising force.

We did observe distinct changes in anaphase dynamics under nocodazole washout treatment, suggestive of a decreased segregation efficacy. Firstly, on the descending clusters - there was a substantial increase in average cluster spread from 0.345 to 0.610 microns (*i*.*e*., 77% increase) within the first 120s from anaphase onset, Figure D1B, whilst DMSO cluster size was invariant to a 17.5% tolerance (minimum to maximum mean cluster size). Secondly, we observed a higher frequency of transient direction reversal events, [Skibbens et al., 1993], Appendix Figure D1. KTs with reversals had stronger pulling K-fibers, Appendix Table A12, Appendix Figures D6 and D7, consistent with a higher proportion of strong oscillators having reversals, Appendix Table A11. Whether such reversals impact segregation efficacy is unknown.

Our data driven analysis suggests that human KT dynamics analysis should incorporate sister asymmetry and temporal variation. In particular, the following model is suggested (sister *k*),

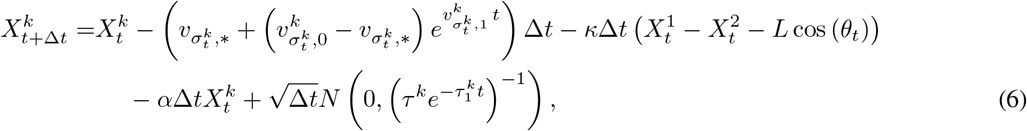

where *v*_± *_, is the anaphase ready state. Time resolution, spatial resolution and signal to noise will impact the ability to infer all these processes. Further work is required to confirm support for this joint asymmetric, temporal model, and determine whether it is identifiable, and the requirements on time and spatial resolution to do so. This is a model for individual KT pairs. There is also substantial variation in biophysical parameters with metaphase plate position, although we have not determined the functional form of this variation. This could be inferred using Gaussian process modelling.

Our results are dependent on our models capturing the key mechanical processes driving KT dynamics. Our data driven modelling approach has demonstrated that fundamental properties of KT biophysical behaviour can be extracted, so although we have used simple models that match the data complexity they have powerful knowledge generating capability. Further improvements are possible, both overcoming the simplifying assumptions of the current models and incorporating additional processes into the biophysical models to improve their realism, although additional relevant datasets will be required to parametrise those additional processes. Our models are in 1D, with K-fiber forces assumed to lie along the axis. In practice, there is slight curvature of the trajectories in metaphase as they track the spindle geometry, so *v*_±_ may be underestimated towards the periphery. The impact of bridging fibers on KT dynamics is also unknown; acute removal of PRC1, which weakens such fibers, does lead to changes in sister kinetochore tension and orientation, [Jagrić et al., 2021]. Extending the models and analysis to 3D and incorporating bridging fibers, would thus capture these key difference between peripheral and central KTs. Our models are in terms of composite forces, specifically *v*_±_ are the K-fiber pushing/anti-poleward, pulling/poleward forces respectively, comprising (de)polymerisation at the MT plus ends and retrograde flux (depolymerisation at the spindle pole). *α* models a linear centralising force comprising the PEF, flux driven centralisation Risteski et al. [2022] and regulation of K-fiber plus end dynamics by HURP Drpic et al. [2018]. Deconvolving these composite forces, thus establishing their relative contributions will required additional experiments that perturb individual processes. Also, the model for drag forces may also be too simplistic since the spindle behaves as a visco-elastic anisotropic material, [Shimamoto et al., 2011], whilst MT bundles exhibit high transverse drag Nazockdast et al. [2017], Assessing the impact of this visco-elasticity, and the spindle structure more generally on KT dynamics would require rheology experiments in association with KT tracking. Interactions between kinetochore pairs, [Vladimirou et al., 2013] can also be incorporated into the models.

## Acknowledgements

We gratefully acknowledge the initial work and software development by Jonathan Harrison.C.K., A.V.I, A.D. were supported by BBSRC (BB/R009503/1), AD and ADM supported by Wellcome Discovery Award (226605/Z/22/Z) and ADM supported by Wellcome Investigator Award (106151/Z/14/Z)

## 4 Methods and Materials

### Code

The trajectory data and original code used to produce the results reported in this work are available at Github, https://github.com/ckoki21/MetaAnaDynamics.git. Any additional information required to re-analyze the data reported in this paper is available from the lead contact upon request.

### Cell culture and generation of cell lines

Immortalized (hTERT) diploid human retinal pigment epithelial (RPE1) cell line (MC191), expressing endogenously tagged Ndc80-eGFP, was generated by CRISPR-Cas9 gene editing, [Roscioli et al., 2020]. hTERT-RPE1 cells were grown in DMEM/F-12 medium containing 10% fetal bovine serum (FBS), 2 mM L-glutamine, 100 U/ml penicillin and 100 mg/ml streptomycin (full growth medium); and were maintained at 37ºC with 5% CO_2_ in a humidified incubator.

### Live cell imaging by lattice light sheet microscope

The lattice light sheet microscope (LLSM), [Chen et al., 2014], used in this study was manufactured by 3i (https://www.intelligent-imaging.com). Cells were seeded on 5 mm radius glass coverslips one day before imaging. On the imaging day, each coverslip was transferred to the LLSM bath filled with CO_2_-independent L15 medium, where live imaging takes place. All imaged cells entered anaphase, which is a suitable proxy for a lack of phototoxicity effects, [Jaqaman et al., 2010]. The LLSM light path was aligned at the beginning of every imaging session by performing beam alignment, dye alignment and bead alignment, followed by the acquisition of a bead image (at 488 nm channel) for measuring the experimental point spread function (PSF). This PSF image is later used for the deconvolution of images. 3D time-lapse images (movies) of Ndc80-eGFP were acquired at 488nm channel using 1% laser power, 20 ms exposure time/z-plane, 75 z-planes, 307 nm z-step and 0.5 s laser off time, which results in 2 s/z-stack time/frame. Acquired movies were de-skewed and cropped in XYZ and time, using Slidebook software in order to reduce the file size. Cropped movies were then saved as OME-TIFF files in ImageJ.

### Tracking

Kinetochore tracking is performed using the software package KiT v3.0. The tracking algorithm proceeds by detecting candidate spots via a constant false alarm rate (CFAR) detection method, [Daniyan et al., 2024], to set a KT-wise dynamic threshold on per image frame in a movie. Candidate spot locations are refined by fitting a Gaussian mixture model. Spot locations are linked between frames by solving a linear assignment problem, with motion propagation via a Kalman filter. Tracked kinetochores are paired by solving another linear assignment problem. Sister kinetochore pairing used spatial and temporal trajectory data. Sister kinetochores are closer (*d*_*ij*_, average distance) and exhibit less distance variation (*v*_*ij*_, distance variance) than non sisters. In the presence of a metaphase plate, their connecting vector aligns with the plate’s normal (*α*_*ij*_, average angle). Pairing costs were calculated as *d*_*ij*_ ×*v*_*ij*_ without a plate or *d*_*ij*_ ×*v*_*ij*_ ×*α*_*ij*_ with a plate. Trajectories had to overlap for at least 10 frames, and the configuration with the minimum global cost was selected.

A quality control filter was applied to ensure good sister pair coverage per cell. Specifically, at least 30 sister pairs had to be tracked for 75% of the movie length, see Appendix C1.

The code to perform kinetochore tracking is available from https://github.com/cmcb-warwick/KiT, and this software includes a graphical user interface (GUI) for ease of use.

### Statistical Tests

The statistical tests used in this study can be found in Appendix C, Table C3.

## Appendix

### A Additional tables

**Table A1:** Support for asymmetry models in DMSO and nocodazole washout treatment. Numbers of KT pairs that prefer each asymmetric model divided into strength categories. The strength of asymmetry is based on the categorisation of the strength of evidence of asymmetry over the symmetric model, using the Bayes Factors, Kass and Raftery [1995b], *i*.*e*., substantial, strong, decisive. At least substantial evidence is shown (Substantial+), the sum of the previous 3 categories. We observe that asymmetric sisters are predominately asymmetric on *v*_−_.

| Preferred Model | $v_-, v_+, \tau$ | $v_-, v_+$ | $v_-$ | $v_+$ | Totals |
| --- | --- | --- | --- | --- | --- |
| **DMSO** |  |  |  |  |  |
| Strength of Asymmetry |  |  |  |  |  |
| Substantial | 1 | 24 | 36 | 13 | 74 |
| Strong | 1 | 27 | 32 | 8 | 68 |
| Decisive | 0 | 37 | 15 | 1 | 53 |
| Substantial+ | 2 (1.03%) | 88 (45.13%) | 83 (42.56%) | 22 (11.28%) | 195 |
| **Nocodazole washout** |  |  |  |  |  |
| Strength of Asymmetry |  |  |  |  |  |
| Substantial | 0 | 23 | 35 | 29 | 87 |
| Strong | 0 | 39 | 33 | 15 | 87 |
| Decisive | 2 | 56 | 47 | 7 | 112 |
| Substantial+ | 2 (0.7%) | 118 (41.46%) | 115 (40.07%) | 51 (17.77%) | 286 |

**Table A2:** K-fiber force variation within cells is larger than between cells. Variance contributions of the biophysical parameters, (*n* = 635 KT pairs from *N* = 24 cells to allow for at least 3 sister pairs in each group and remove outliers), grouping by cell and location (3 radius group classes) within the metaphase plate. The second column denotes the within-cell variance (*i*.*e*., residuals), while the third and fourth columns report the within cell and location variance respectively. Parameter estimations use the model with asymmetry on *v*_−_ and *v*_+_; the posterior median *v*_−_ and *v*_+_ values are averaged over the two sisters. Stars denote that there is statistically significant difference (*α <* 0.01) between groups of this factor. The ratio of the between cell to the between location variance is shown in fifth column and the size of effect (ratio of population parameters’ standard deviation to the population parameters’ (abs) mean) in last column. Stars denote the statistically different ratio at *α <* 0.01. Caution: Due to low coverage of KT pairs per cell within each radius group, these results should be taken with caution as some of the ANOVA assumptions are violated.

| Parameter | Within variance | Between-cell variance | Between-location variance | Between cell/location variance | sd/mean |
| --- | --- | --- | --- | --- | --- |
| $\alpha$ | $4.0 \times 10^{-5}$ | $2.3 \times 10^{-6*}$ | $4.6 \times 10^{-8}$ | 50.256 | 0.508 |
| $\kappa$ | $2.1 \times 10^{-4}$ | $3.6 \times 10^{-5*}$ | $2.9 \times 10^{-5*}$ | 1.256 | 0.578 |
| $L$ | $9.0 \times 10^{-4}$ | $1.1 \times 10^{-4*}$ | $9.9 \times 10^{-6*}$ | 11.416 | 0.037 |
| $\tau$ | $3.2 \times 10^4$ | $1.9 \times 10^4*$ | $3.1 \times 10^3*$ | 5.987 | 0.260 |
| $v_-$ | $1.8 \times 10^{-4}$ | $2.3 \times 10^{-5*}$ | $4.1 \times 10^{-5*}$ | 0.569 | 0.303 |
| $v_+$ | $3.4 \times 10^{-5}$ | $9.7 \times 10^{-6*}$ | $9.0 \times 10^{-7*}$ | 10.723 | 0.562 |
| $p_{coh}$ | $2.3 \times 10^{-4}$ | $3.5 \times 10^{-5*}$ | $2.4 \times 10^{-5*}$ | 1.451 | 0.279 |
| $p_{incoh}$ | $1.2 \times 10^{-2}$ | $1.7 \times 10^{-3*}$ | $3.0 \times 10^{-3*}$ | 0.582 | 0.149 |

**Table A3:**
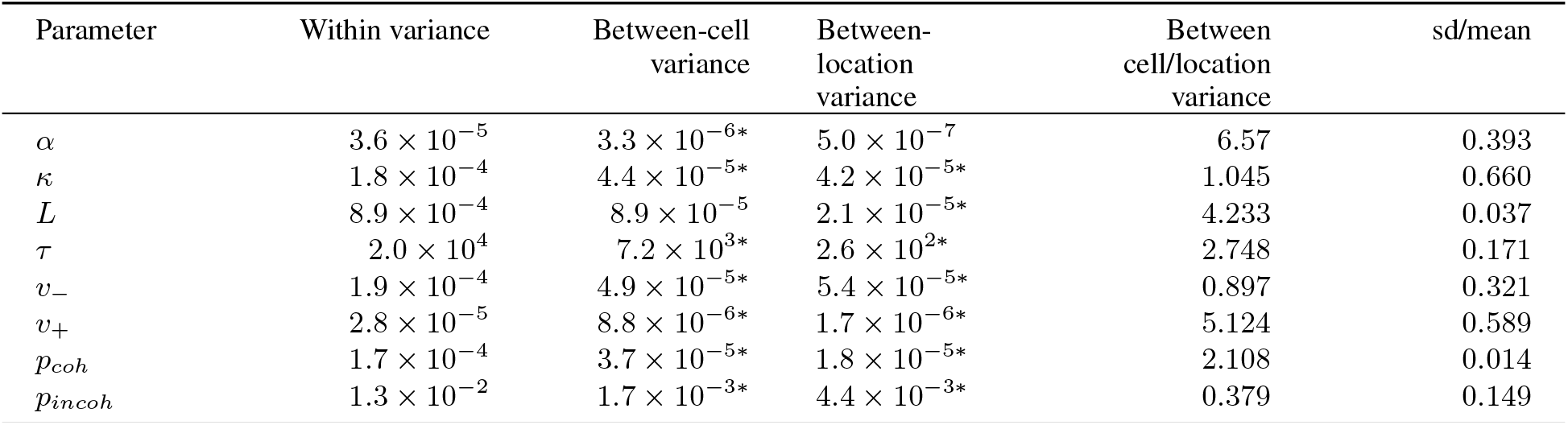
Variance contributions of the biophysical parameters, (*n* = 290 KT pairs from *N* = 11 cells to allow for at least 3 sister pairs in each group), grouping by cell and location (3 radius group classes) within the metaphase plate in nocodazole washout treated cells. The second column denotes the within-cell variance (i.e., residuals), while the third and fourth columns report the within cell and location variance respectively. Parameter estimations use the model with asymmetry on *v*_−_ and *v*_+_; the posterior median *v*_−_ and *v*_+_ values are averaged over the two sisters. Stars denote that there is statistically significant difference (*α <* 0.01) between groups of this factor. The ratio of the between cell to the between location variance is shown in fifth column and the size of effect (ratio of population parameters’ standard deviation to the population parameters’ (abs) mean) in last column. Stars denote the statistically different ratio at *α <* 0.01. Caution: Due to low coverage of KT pairs per cell within each radius group, these results should be taken with caution as some of the ANOVA assumptions are violated.

**Table A4:** Summary statistics of symmetric and asymmetric sisters of nocodazole washout treated cells. We label sister 1 as the sister with the greater pulling force, *i*.*e*., 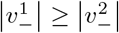. The first and third columns reports the medians of posterior medians for symmetric and asymmetric sisters respectively, while brackets report the population interquartile range. The second and fourth columns report the median of Posterior Interquartile Ranges (PIR) for symmetric and asymmetric sisters, respectively. Based on Mann-Whitney and Kolmogorov-Smirnov tests, the all asymmetric posterior distributions but *τ* are significantly different that the symmetric posterior distributions, with size of effect (*i*.*e*., percentage difference of the medians) being 15.7%, 26.4% and 9.5%, 20.6%, 2.6% for *κ, α, v*_−_, *v*_+_, *L* respectively. Note that asymmetric parameter values on *v*_−_ and *v*_+_ are averaged over the two sisters.

|  | Symmetric |  | Asymmetric |  |
| --- | --- | --- | --- | --- |
|  | Median | PIR | Median | PIR |
| $\alpha$ | 0.012 [0.009] | 0.005 | 0.016 [0.009] | 0.005 |
| $\kappa$ | 0.017 [0.017] | 0.013 | 0.019 [0.024] | 0.013 |
| $L$ | 0.813 [0.042] | 0.176 | 0.840 [0.077] | 0.156 |
| $\tau$ | 830.6 [265.34] | 86.42 | 824.1 [229.1] | 83.77 |
| $v_-^1$ | -0.039 [0.020] | 0.006 | -0.053 [0.018] | 0.009 |
| $v_-^2$ | -0.039 [0.020] | 0.005 | -0.030 [0.021] | 0.009 |
| $v_+^1$ | 0.006 [0.007] | 0.005 | 0.002 [0.002] | 0.007 |
| $v_+^2$ | 0.006 [0.07] | 0.005 | 0.016 [0.009] | 0.007 |
| $p_{icoh}$ | 0.777 [0.220] | 0.120 | 0.803 [0.226] | 0.133 |
| $p_{coh}$ | 0.947 [0.0208] | 0.021 | 0.952 [0.019] | 0.020 |

**Table A5:** Spatial biophysical parameter trends across the metaphase plate: p-values of Pearson’s, Kendall’s and Spearman’s tests for the presence of a trend, in 798 kinetochore pairs over 26 DMSO treated cells.

| Preferred Model | Pearson | Kendal | Spearman |
| --- | --- | --- | --- |
| $\alpha$ | $4.8 \times 10^{-1}$ | $1.2 \times 10^{-1}$ | $1.2 \times 10^{-1}$ |
| $\kappa$ | $2.2 \times 10^{-16}$ | $2.2 \times 10^{-16}$ | $2.2 \times 10^{-16}$ |
| $L$ | $9.8 \times 10^{-1}$ | $1.2 \times 10^{-3}$ | $1.2 \times 10^{-3}$ |
| $\tau$ | $1.9 \times 10^{-15}$ | $6.5 \times 10^{-15}$ | $9.4 \times 10^{-15}$ |
| $v_-$ | $2.2 \times 10^{-16}$ | $2.2 \times 10^{-16}$ | $2.2 \times 10^{-16}$ |
| $v_+$ | $7.8 \times 10^{-7}$ | $2.7 \times 10^{-7}$ | $2.5 \times 10^{-7}$ |

**Table A6:** Spatial biophysical parameter trends across the metaphase plate for a single cell: p-values of Pearson’s test for the presence of a trend, in 798 kinetochore pairs over 26 DMSO treated cells, and for 36 KT pairs in one DMSO treated cell.

| Preferred Model | Population | Cell |
| --- | --- | --- |
| $\alpha$ | $4.8 \times 10^{-1}$ | $6.5 \times 10^{-1}$ |
| $\kappa$ | $2.2 \times 10^{-16}$ | $4.5 \times 10^{-5}$ |
| $L$ | $9.8 \times 10^{-1}$ | $1.9 \times 10^{-1}$ |
| $\tau$ | $1.9 \times 10^{-5}$ | $2.3 \times 10^{-15}$ |
| $v_-$ | $2.2 \times 10^{-16}$ | $2.2 \times 10^{-5}$ |
| $v_+$ | $7.8 \times 10^{-7}$ | $7.5 \times 10^{-1}$ |

**Table A7:** Comparison of medians for DMSO and nocodazole washout treated cells across model parameters. The relative change column quantifies the percentage difference from DMSO to nocodazole washout cells. Negative percentages indicate a reduction in the nocodazole condition.

| Parameter | DMSO | Nocodazole washout | Relative Change |
| --- | --- | --- | --- |
| $\alpha$ | 0.0131 | 0.0148 | 12.9% |
| $\kappa$ | 0.0277 | 0.0226 | −18.4% |
| $L$ | 0.820 | 0.841 | 2.56% |
| $\tau$ | 703 | 844 | 20.0% |
| $v_-$ | −0.0446 | −0.0420 | −5.85% |
| $v_+$ | 0.0100 | 0.00625 | −37.5% |
| $p_{coh}$ | 0.937 | 0.944 | 0.75% |
| $p_{incoh}$ | 0.741 | 0.766 | 3.37% |

**Table A8:** Oscillations deteriorate towards the MPP periphery. Percentages indicate the proportion of each oscillator type in the far (*r* ≥ 3.3*µm*) versus near (*r <* 3.3*µm*) distance from the metaphase plate origin. There are significant differences in both DMSO treated cells (*p*_Pearson_ = 5.48 ×10^−8^) and nocodazole washout treated cells (*p*_Pearson_ = 0.00044).

| Cluster | DMSO | | Total ($n = 836$) |
| --- | --- | --- | --- |
| | $r < 3.3 \mu m, n = 418$ | $r \geq 3.3 \mu m, n = 418$ | |
| strong/short | 40 (9.6%) | 30 (7.2%) | 70 (8.4%) |
| strong/long | 125 (29.9%) | 64 (15.3%) | 189 (22.6%) |
| weak/long | 52 (12.4%) | 36 (8.6%) | 88 (10.5%) |
| weak/noisy | 78 (18.7%) | 112 (26.8%) | 190 (22.7%) |
| poor | 123 (29.4%) | 176 (42.1%) | 299 (35.8%) |

Table A8: **Oscillations deteriorate towards the MPP periphery.**
| Cluster | Nocodazole washout | | Total ($n = 971$) |
| --- | --- | --- | --- |
| | $r < 3.3 \mu m, n = 485$ | $r \geq 3.3 \mu m, n = 486$ | |
| strong/short | 69 (14.2%) | 45 (9.3%) | 114 (11.7%) |
| strong/long | 147 (30.3%) | 109 (22.5%) | 256 (26.4%) |
| weak/long | 46 (9.5%) | 52 (10.7%) | 98 (10.1%) |
| weak/noisy | 96 (19.8%) | 142 (29.3%) | 238 (24.5%) |
| poor | 128 (26.3%) | 137 (28.2%) | 265 (27.3%) |

**Table A9:** Strong and poor oscillators have distinctly different biophysical parameters. Hypothesis tests for 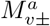 model parameters comparing strong and poor/not oscillators using the Mann-Whitney test. We used *n* = 257 (strong oscillators) and *n* = 359 (poor/not oscillators) KT-pair estimates for DMSO-treated cells, and *n* = 294 (strong oscillators) and *n* = 244 (poor/not oscillators) for nocodazole washout treated cells. P-values are adjusted using Benjamini-Hochberg to control for 16 multiple comparisons (8 variables × 2 conditions). Statistically significant tests (*p <* 0.05) are marked with *.

| Variable (medians) | $p$-value | BH-adjusted $p$ |
| --- | --- | --- |
| DMSO |  |  |
| $\alpha$ | $5.75 \times 10^{-6}$ | $1.15 \times 10^{-5*}$ |
| $\kappa$ | $7.70 \times 10^{-5}$ | $1.23 \times 10^{-4*}$ |
| $L$ | $1.79 \times 10^{-2}$ | $2.05 \times 10^{-2*}$ |
| $\tau$ | $9.29 \times 10^{-1}$ | $9.29 \times 10^{-1}$ |
| $v_-$ | $7.20 \times 10^{-22}$ | $5.76 \times 10^{-21*}$ |
| $v_+$ | $4.07 \times 10^{-3}$ | $5.43 \times 10^{-3*}$ |
| $p_{coh}$ | $1.80 \times 10^{-6}$ | $4.80 \times 10^{-6*}$ |
| $p_{incoh}$ | $1.25 \times 10^{-13}$ | $5.00 \times 10^{-13*}$ |
| Nocodazole washout |  |  |
| $\alpha$ | $6.62 \times 10^{-11}$ | $2.65 \times 10^{-10*}$ |
| $\kappa$ | $2.57 \times 10^{-1}$ | $2.94 \times 10^{-1}$ |
| $L$ | $8.11 \times 10^{-1}$ | $8.11 \times 10^{-1}$ |
| $\tau$ | $6.70 \times 10^{-2}$ | $1.07 \times 10^{-1}$ |
| $v_-$ | $1.22 \times 10^{-15}$ | $9.76 \times 10^{-15*}$ |
| $v_+$ | $7.34 \times 10^{-2}$ | $9.79 \times 10^{-2}$ |
| $p_{coh}$ | $3.94 \times 10^{-5}$ | $7.88 \times 10^{-5*}$ |
| $p_{incoh}$ | $9.39 \times 10^{-9}$ | $2.51 \times 10^{-8*}$ |

**Table A10:** Strong oscillators are more prevalent towards the centre of the metaphase plate, and have a significantly larger directional switching time. Comparison of *p*_*incoh*_ distributions between strong and poor oscillators across different radial positions from the metaphase plate intervals in DMSO and Nocodazole conditions. The radius intervals represent quintiles of average radius; *n*_*strong*_ and *n*_*poor*_ indicate the number of observations in each group. p-values are from Wilcoxon tests comparing distributions of *p*_*incoh*_ between strong and poor oscillators within each radius group, with adjusted p-values controlling for multiple testing.

| DMSO |  |  |  |  |
| --- | --- | --- | --- | --- |
| Radius Interval ($\mu m$) | $n_{strong}$ | $n_{poor}$ | p-value | p-value (adj.) |
| [0.40, 1.99] | 65 | 46 | 0.02 | 0.04* |
| [2.02, 2.86] | 63 | 47 | 0.13 | 0.13 |
| [2.90, 3.58] | 53 | 57 | $1.56 \times 10^{-4}$ | $7.81 \times 10^{-4*}$ |
| [3.58, 4.00] | 41 | 69 | $3.00 \times 10^{-3}$ | $7.51 \times 10^{-3*}$ |
| [4.00, 4.99] | 35 | 75 | 0.04 | 0.05* |
| Nocodazole |  |  |  |  |
| [0.32, 2.06] | 77 | 44 | 0.09 | 0.11 |
| [2.06, 2.87] | 76 | 45 | 0.18 | 0.18 |
| [2.87, 3.50] | 79 | 42 | $3.20 \times 10^{-3}$ | $7.99 \times 10^{-3*}$ |
| [3.51, 4.01] | 67 | 53 | 0.01 | 0.02* |
| [4.02, 4.92] | 60 | 60 | $2.48 \times 10^{-3}$ | $7.99 \times 10^{-3*}$ |

**Table A11:** Trajectories with reversal events are more frequent in strong oscillators. This table summarises the number of pairs in each oscillator type cluster (in the pulled dataset), the number of trajectories with reversal events per oscillator type and the relevant proportion. One-tailed two-proportion z-tests for reversal proportions showed that the reversal proportions are higher in strong oscillators. Note that this is obvious when comparing (a) Strong (Short + Long) vs Weak (Noisy + Long): *p*_*Ztest*_ = 0.0029 and (b) Strong (Short + Long) vs Weak + Poor: *p*_*Ztest*_ = 1.6 ×10^−5^ oscillators. Both tests show significantly higher proportions of reversals in Strong oscillators.

| Oscillators | $n_{total}$ | $n_{reversals}$ | Proportion of Reversals |
| --- | --- | --- | --- |
| Strong/Short | 164 | 70 | 0.427 |
| Strong/Long | 399 | 155 | 0.388 |
| Weak/Long | 167 | 50 | 0.299 |
| Weak/Noisy | 371 | 122 | 0.329 |
| Poor | 496 | 135 | 0.272 |

**Table A12:** Summary of results comparing parameter distributions between of the trajectories without reversal events in the anaphase and with reversal events in DMSO and nocodazole washout treated cells. Medians are reported for each group, and p-values (Mann-Whitney tests) less than 0.05 are marked with ^*^.

| DMSO |  |  |  |
| --- | --- | --- | --- |
| Parameter | Median (No reversals) | Median (Reversals) | p-value |
| $\alpha$ | 0.0109 | 0.0109 | 0.8274 |
| $\kappa$ | 0.0121 | 0.0138 | 0.1356 |
| $L$ | 0.8027 | 0.7953 | 0.0147* |
| $\tau$ | 567.29 | 558.55 | 0.2693 |
| $v_-$ | 0.0372 | 0.0421 | 0.0011* |
| $v_+$ | 0.0089 | 0.0101 | 0.0045* |
| $p_{coh}$ | 0.9565 | 0.9569 | 0.2887 |
| $p_{incoh}$ | 0.7414 | 0.7677 | 0.2710 |
| Nocodazole |  |  |  |
| $\alpha$ | 0.0131 | 0.0134 | 0.2939 |
| $\kappa$ | 0.0102 | 0.0160 | $5.1 \times 10^{-6}$* |
| $L$ | 0.8081 | 0.8146 | 0.0037* |
| $\tau$ | 707.93 | 663.50 | $6.0 \times 10^{-4}$* |
| $v_-$ | 0.0357 | 0.0420 | $1.7 \times 10^{-9}$* |
| $v_+$ | 0.0057 | 0.0056 | 0.5134 |
| $p_{coh}$ | 0.9564 | 0.9519 | 0.0052* |
| $p_{incoh}$ | 0.7561 | 0.8311 | $8.3 \times 10^{-10}$* |

**Figure B1:**
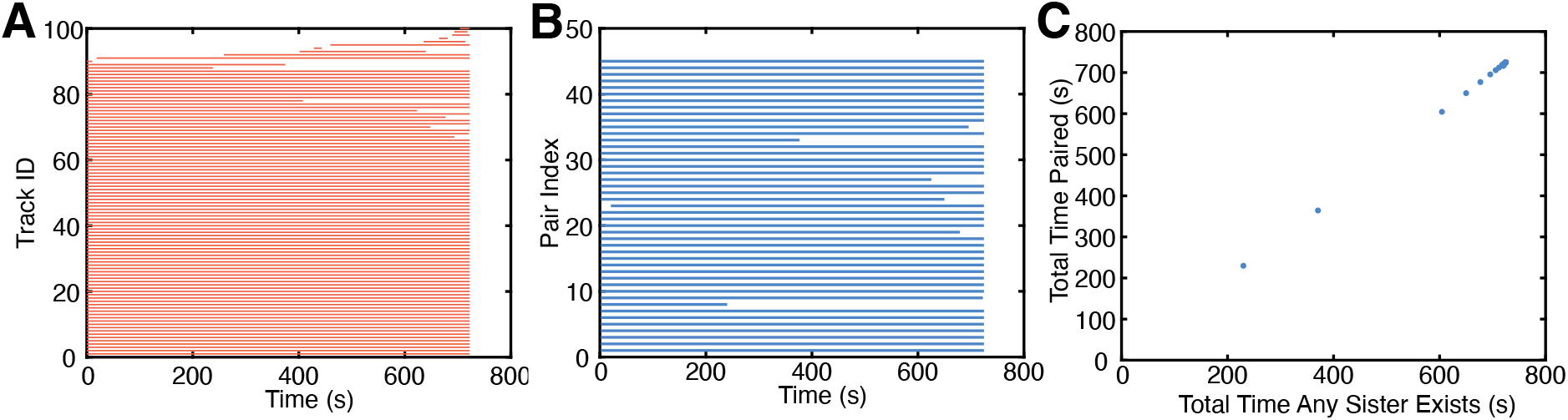
Near-complete tracking of kinetochores through metaphase and anaphase in a human RPE1 cell (tracks are shown in Figure 1). **A** Tracklet plot showing detected KT tracks, tracing the track through time; 82 tracks run through the entire movie, (3 missing time points are allowed, not necessarily consecutive). Because of track breakage, an individual kinetochore could be tracked more than once, resulting in more than a total of 92 tracks. **B** Pairlet plot showing the times in a movie that a sister pair (by pair index) exists (and is paired). **C** Total time a sister pair is paired against total time either sister exists.

**Figure B2:**
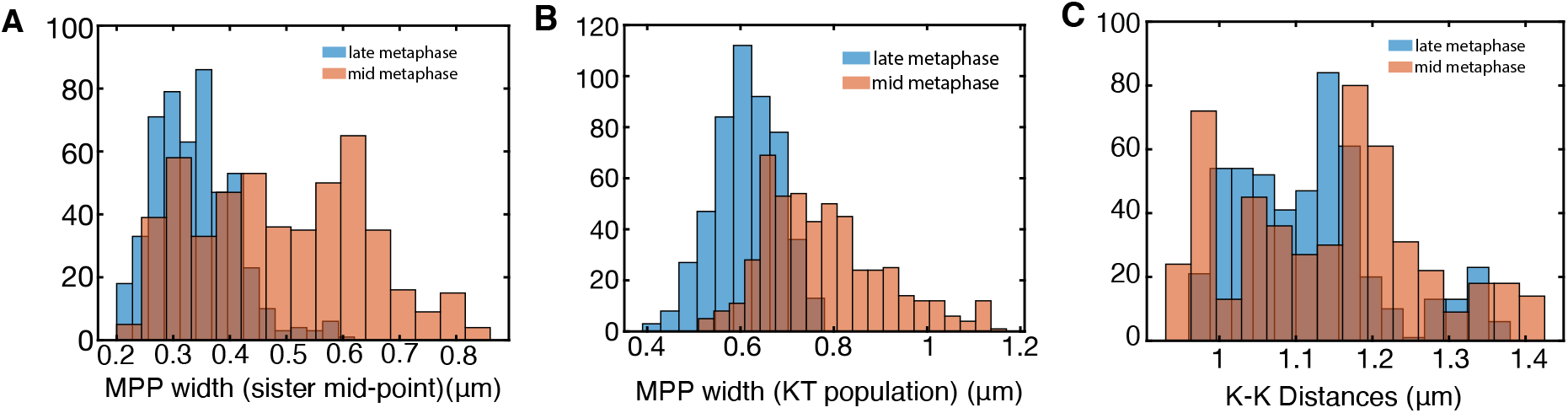
Quantification of intrametaphase maturation. **A** MPP width as measured by paired sister mid-point width (smallest eigenvalue of the covariance matrix of kinetochore mid-points). MPP statistics: **B** Metaphase plate (MPP) width as measured by the covariance matrix of the KT population (smallest eigenvalue). **C** Kinetochore-Kinetochore (KK) distance. Comparison of mid (orange) and late (blue) metaphase as shown in A-C relative to the time before anaphase onset. Data include only cells with at least 30 sisters both tracked for 75% of movie. Mid and late metaphase had significantly different MPP width (both measures), and KK distance (*p*_*MW*_ *<* 10^−3^).

**Figure B3:**
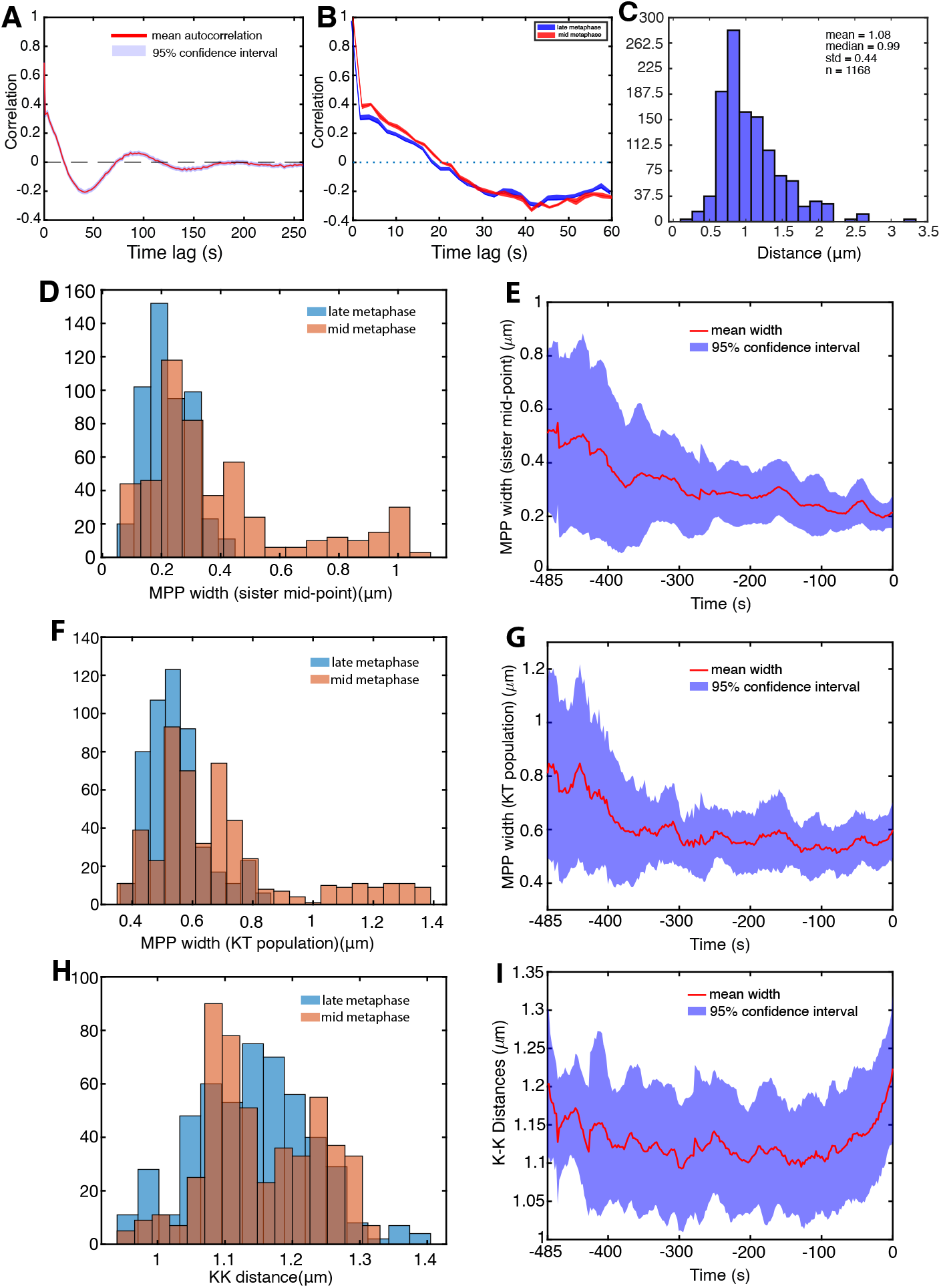
Quantification of intrametaphase maturation in nocodazole washout treated cells. **A** Autocorrelation plot showing the temporal correlation of metaphase oscillations. **B** Autocorrelation of metaphase oscillations in late (red) and mid (blue) metaphase. **C** Sister kinetochore (KK) distance pooled over all KT pairs and time. **D/E** MPP width as measured by paired sister mid-point width (smallest eigenvalue of the covariance matrix of kinetochore mid-points). MPP statistics: **F/G** Metaphase plate (MPP) width as measured by the covariance matrix of the KT population (smallest eigenvalue). **H/I** KK distance. Comparison of mid (orange) and late (blue) metaphase in **D**,**F**,**H**, and time-series (time before anaphase onset) in **E**,**G**,**I**, mean (red) and standard deviation (blue). Data are based on cells having at least 30 sisters both tracked for 75% of the movie (31 cells in total). Early and mid-metaphase median estimates of MPP in D and F where significantly different (minimum *p*_*MW*_ *<* 10^−30^, while the KK-distance in early an mid-metaphase remained the same, *p*_*MW*_ = 0.026.

**Figure B4:**
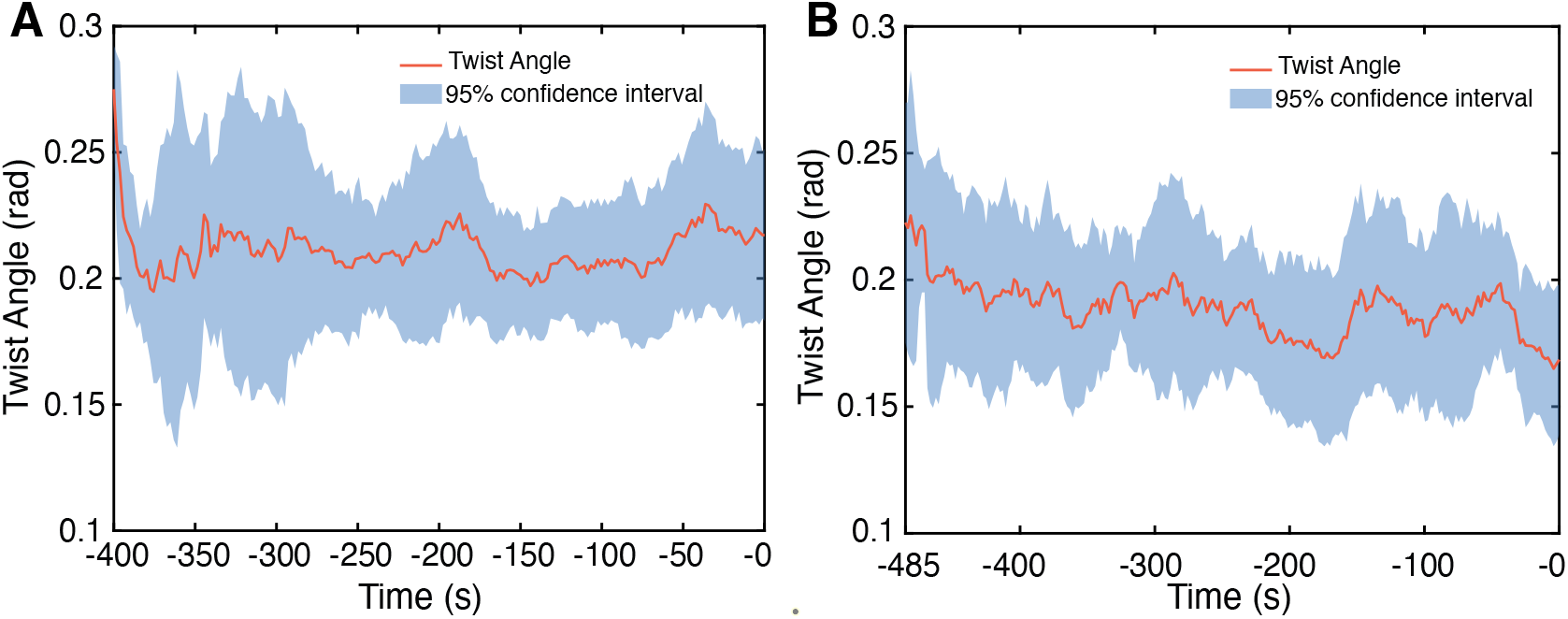
Mean twist angle (in radians) in **A** DMSO **B** nocodazole washout treated data. Shaded blue areas denote the 95% confidence intervals. Time 0 denotes the anaphase onset.

**Figure B5:**
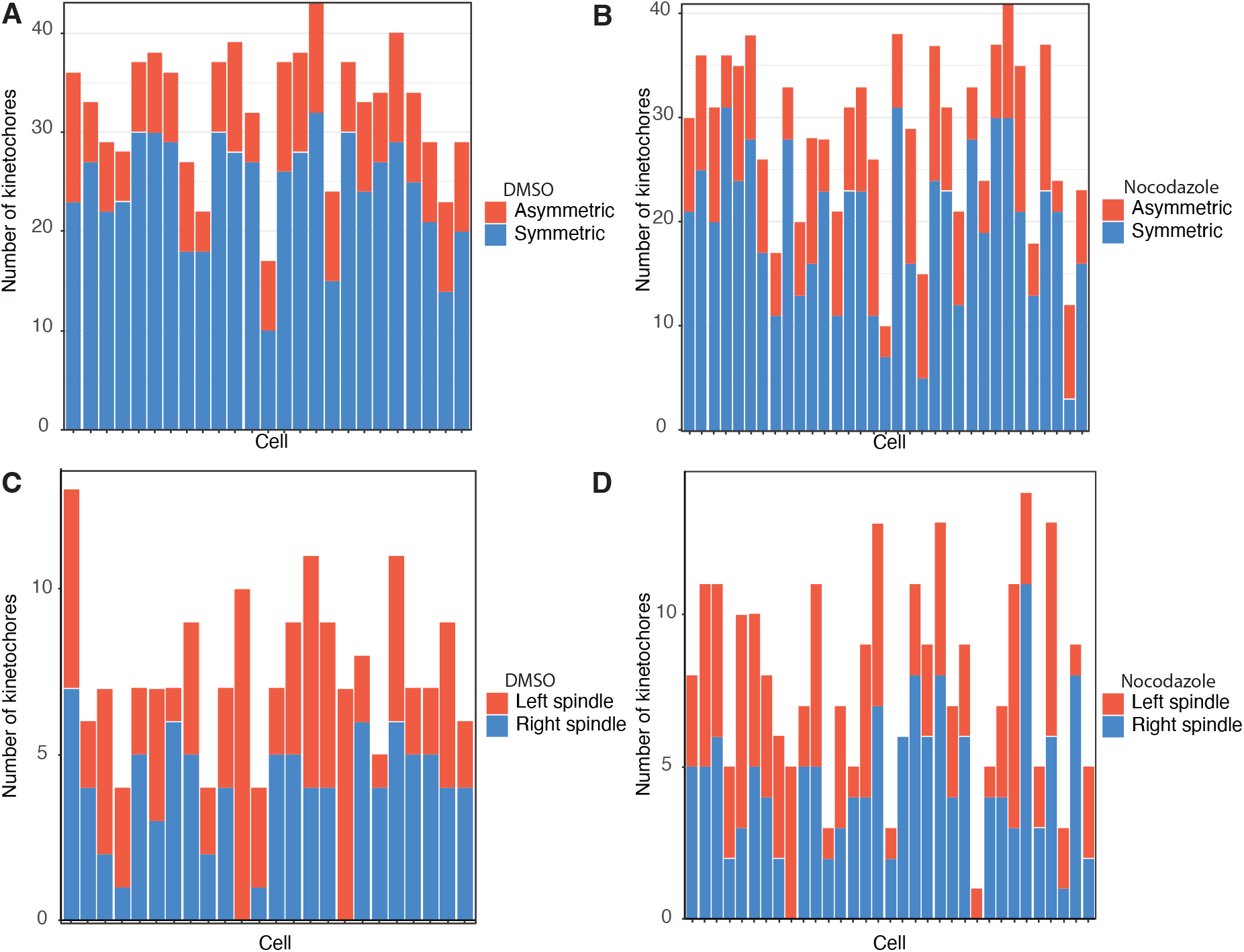
All cells present a significant fraction of asymmetric sisters and no half-spindle bias is observed. **A/B** Symmetric (blue) and significantly asymmetric (red) sister pairs within a cell in **A** DMSO and **B** nocodazole washout treated cells. **C/D** Number of stronger pulling asymmetric sisters positioned in the left (blue) and right (red) half-spindle for **C** DMSO and **D** nocodazole washout treated cells. Stronger pulling asymmetric sisters proportions are not statistically different to *p* = 0.5 in left and right spindle with *p*_*Binom*_ = 0.88 for DMSO and *p*_*Binom*_ = 0.10 for nocodazole washout treated cells.

**Figure B6:**
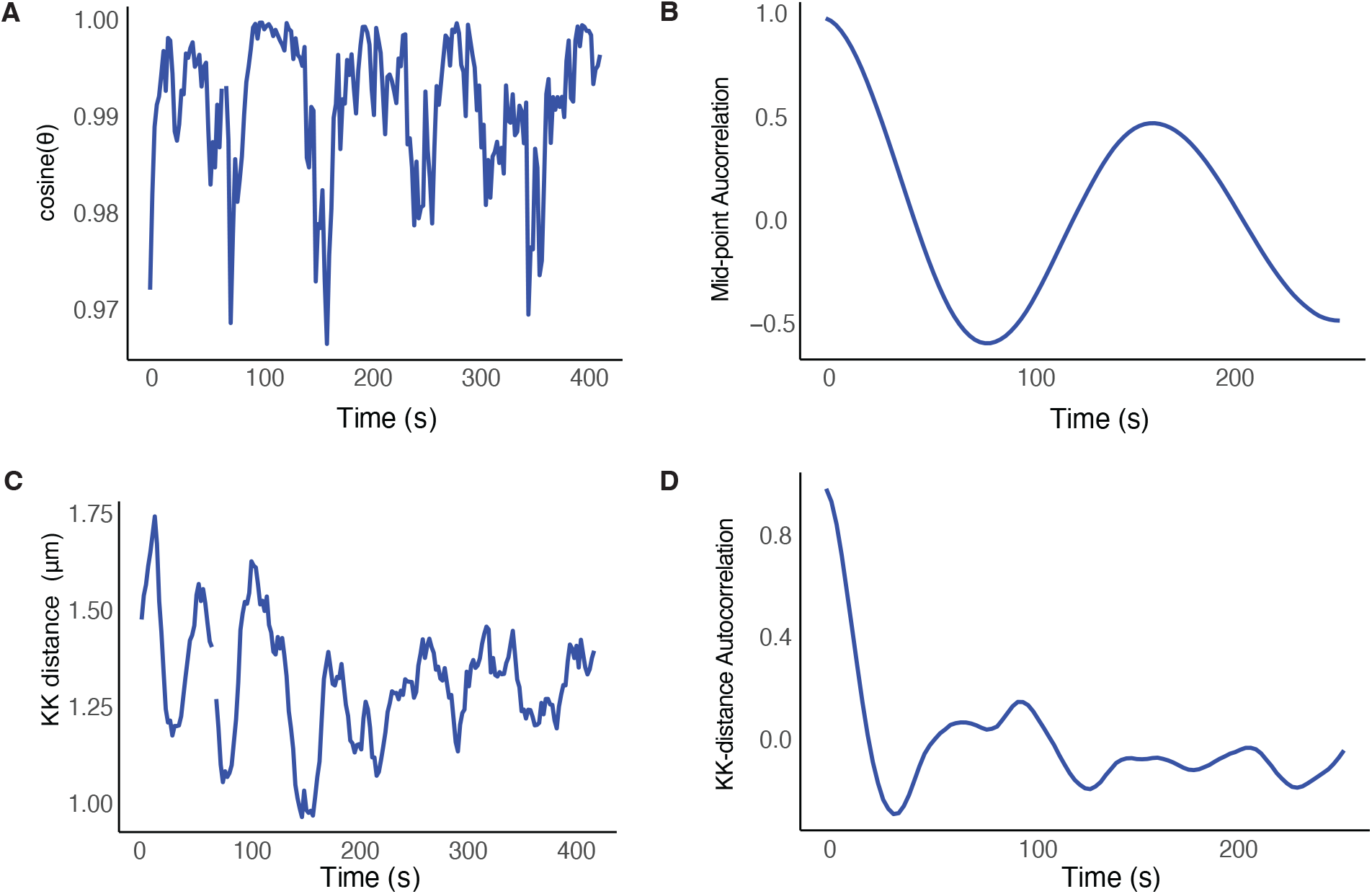
Additional figures for the KT-pair of Figure 4A. **A** Time series of *cosine*(*θ*_*t*_), where *θ*_*t*_ is the angle between the normal to the MPP and the sister to sister vector at time *t*. **B** ACF of sister pair midpoint, confirming that the pair is a good oscillator. **C** Intersister (KK) distance time series. **D** ACF of KK distance (breathing oscillation)

**Figure B7:**
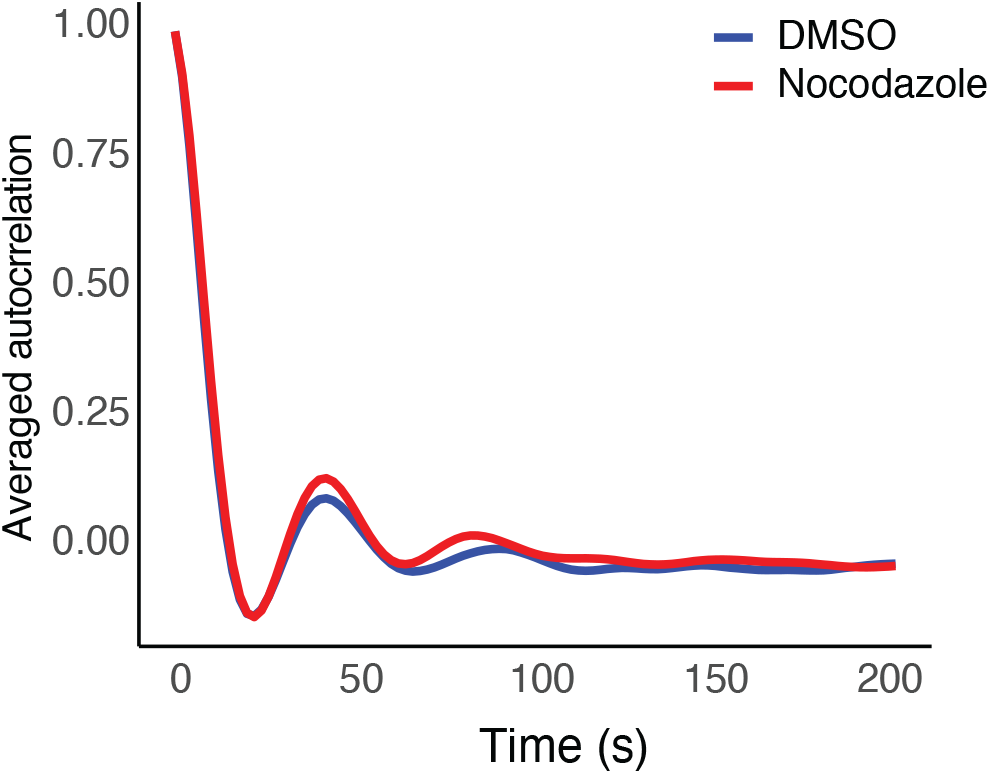
Autocorrelation function of Kinetochore to Kinetochore (KK) sister distance (breathing oscillation): averaged ACF of the distance between sister kinetochores, *i*.*e*., the Kinetochore to Kinetochore (KK) distance for DMSO (blue) and nocodazole washout (red) treated cells. Data based on 31 DMSO cells, 33 nocodazole washout cells.

**Figure B8:**
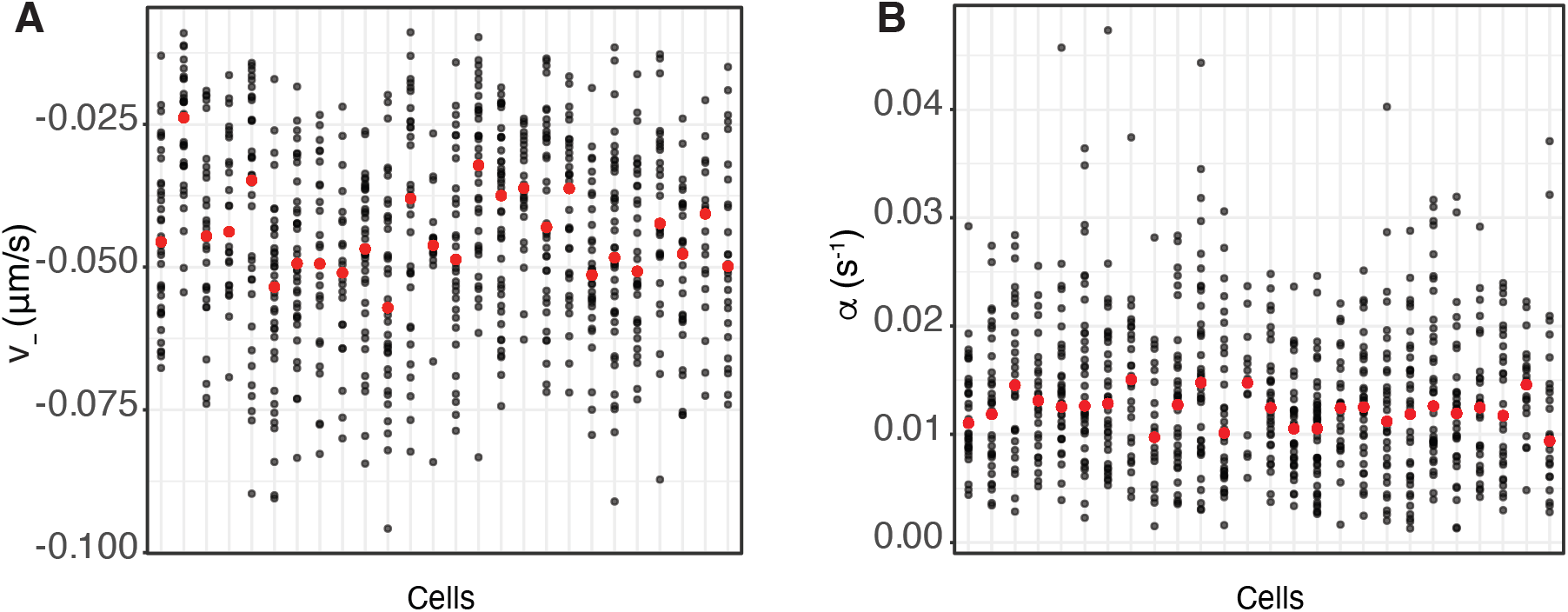
Posterior median estimations of **A** *v*_−_ and **B** *α* shown with respect to cell (x-axis). Red dot denotes the median value of the posterior estimations per cell. Between cell variation is evident in both plots.

**Figure B9:**
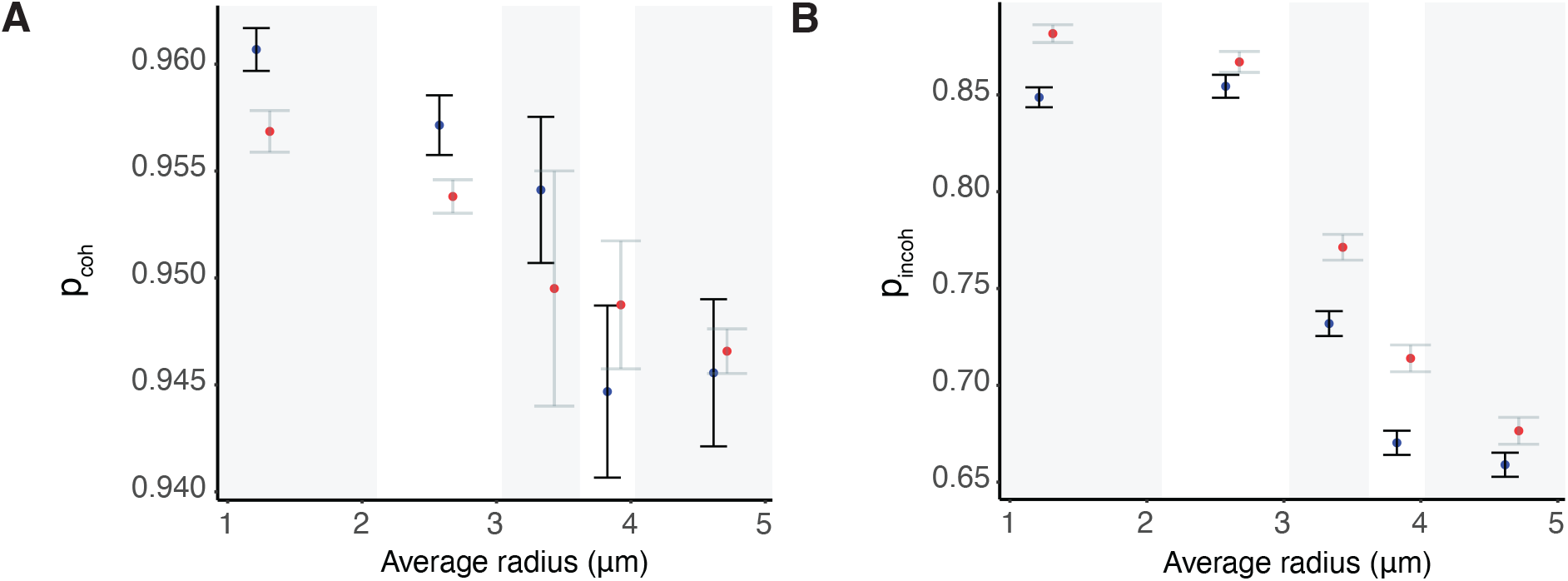
Spatial mechanical trends across the metaphase plate for a DMSO (blue) and nocodazole washout (red) treated cells, complimentary to Figure 10. We report the median estimated values and their interquartile ranges. Groups are equally sized, containing the same number of observations, with the five groups defined by their radial distances, *i*.*e*., [0, 2.03], (2.03, 2.95], (2.95, 3.58], (3.580, 4.01], (4.01, 5.03] respectively. **A** Coherent (*i*.*e*., when hidden states are either +− or −+) probabilities *p*_*coh*_. **B** Incoherent (*i*.*e*., when hidden states are either ++ or −−) probabilities *p*_*incoh*_.

**Figure B10:**
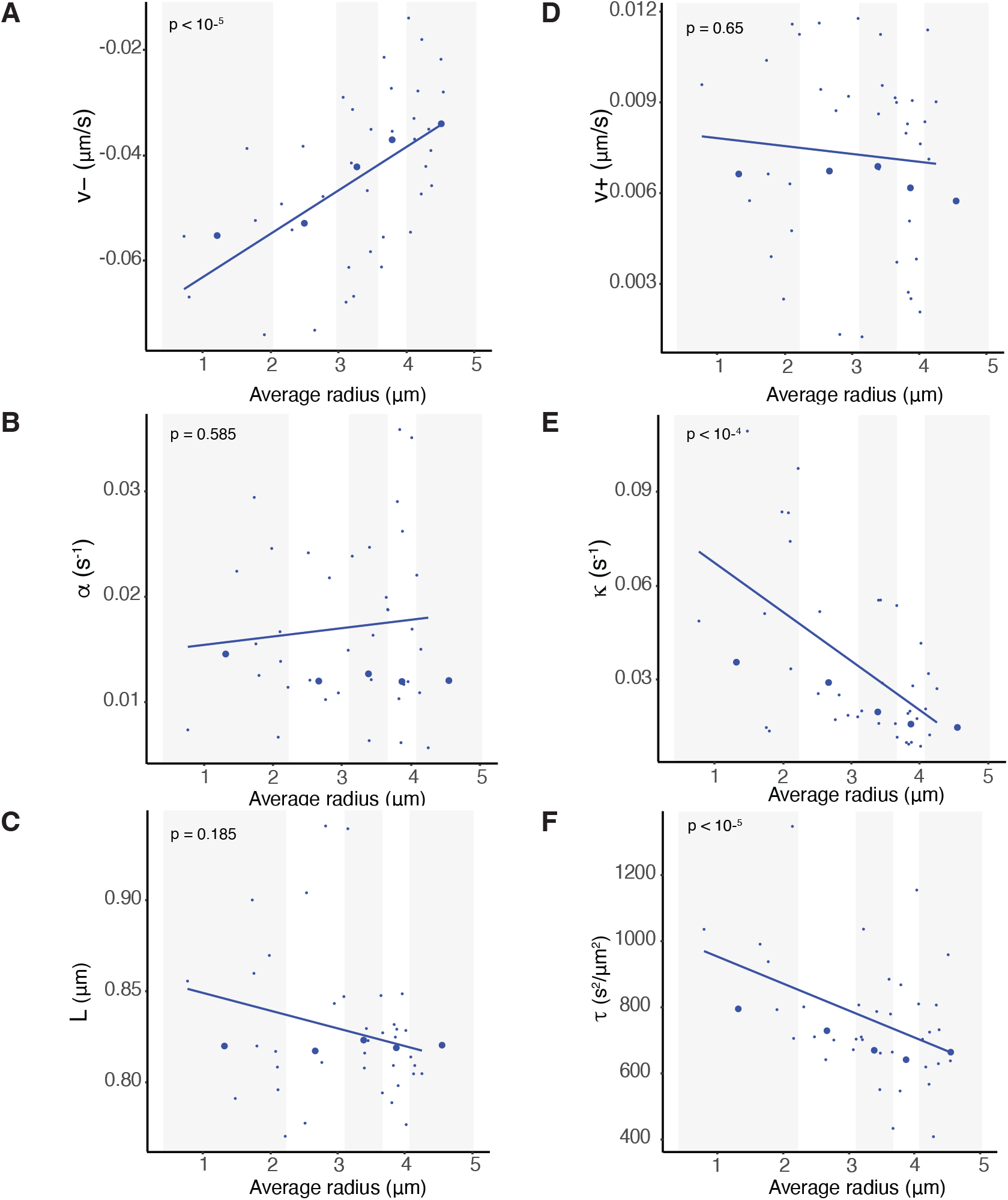
Spatial biophysical parameter trends across the metaphase plate for a single (DMSO) cell. General population trends are evident even on a cell basis. Summary statistics of indicated biophysical parameters partitioned into 5 groups for the average radial position within the metaphase plate. Groups are equally sized, containing the same number of observations, with the five groups defined by their radial distances, *i*.*e*., [0, 2.03], (2.03, 2.95], (2.95, 3.58], (3.580, 4.01], (4.01, 5.03] respectively. **A** Pulling forces *v*_−_, **B** Pushing forces *v*_+_, **C** Spring constant *κ*, **D** Centralising forces, **E** precision *τ*, **F** Natural length of the centromeric chromatin spring connecting the kinetochore sisters *L*.

**Figure B11:**
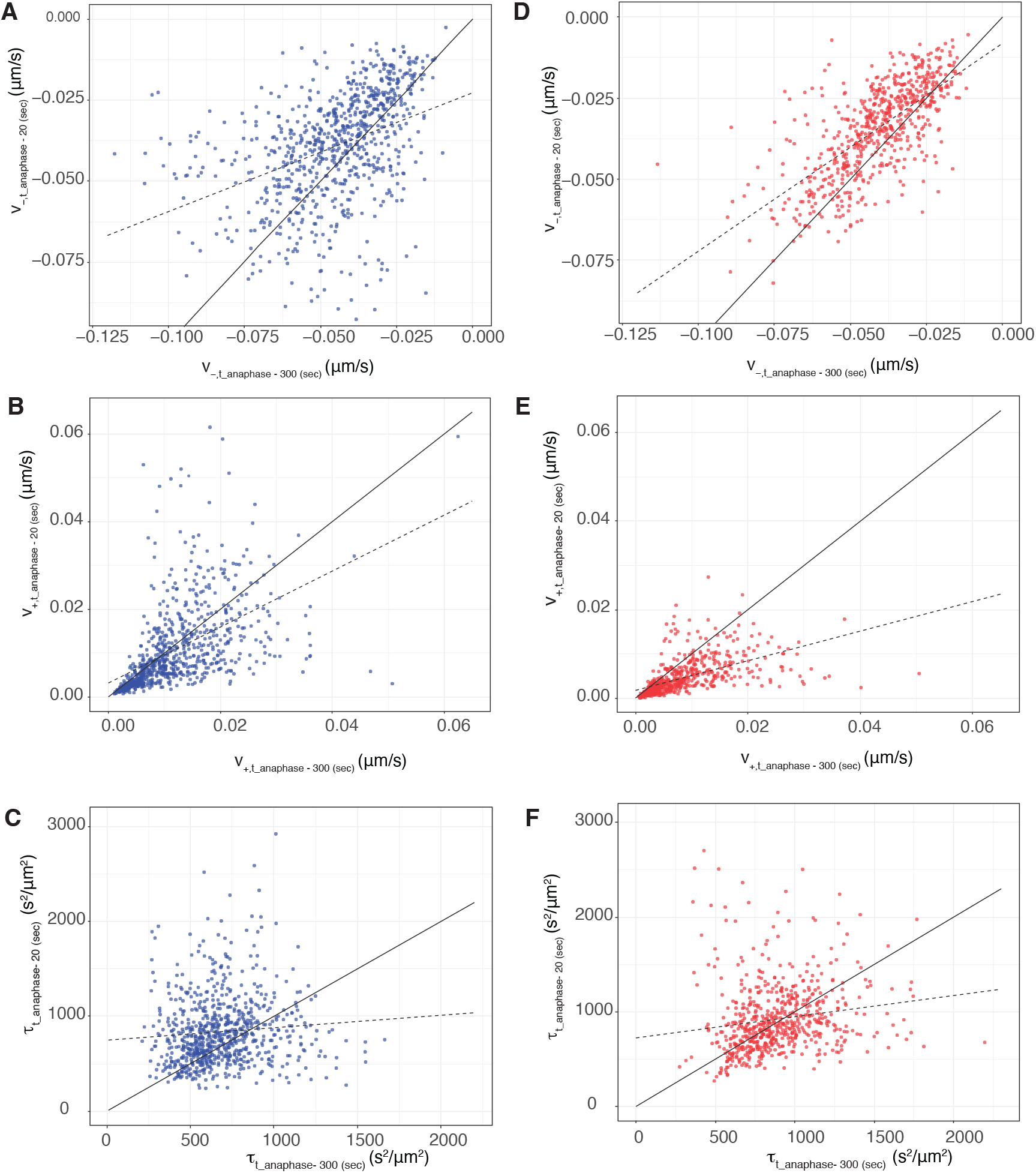
Maturation towards an anaphase ready state. Temporal changes of speed and noise across the last 5 mins of metaphase. Estimated parameters of the model with time dependent parameters *v*_−_, *v*_+_ and *τ* for **A/B/C** DMSO (blue) **D/E/F** nocodazole washout (red) treated cells, respectively. Estimations are based on the median posterior parameters and calculated at two different time points at 300 seconds and 30 seconds prior to anaphase, *i*.*e*., *t* = *t*_*anaphase*_ − 20 (y-axis) and *t* = *t*_*anaphase*_ − 300 (x-axis). The discrete map comprising the line of regression and the 1:1 diagonal indicate the existence of a stable fixed point at *v*_−_ = − 0.04 (*v* _−_ = − 0.02), *v*_+_ = 0.01 (*v*_+_ = 0.003) and *τ* = 849 (*τ* = 923) in DMSO (nocodazole washout) treated cells.

**Figure B12:**
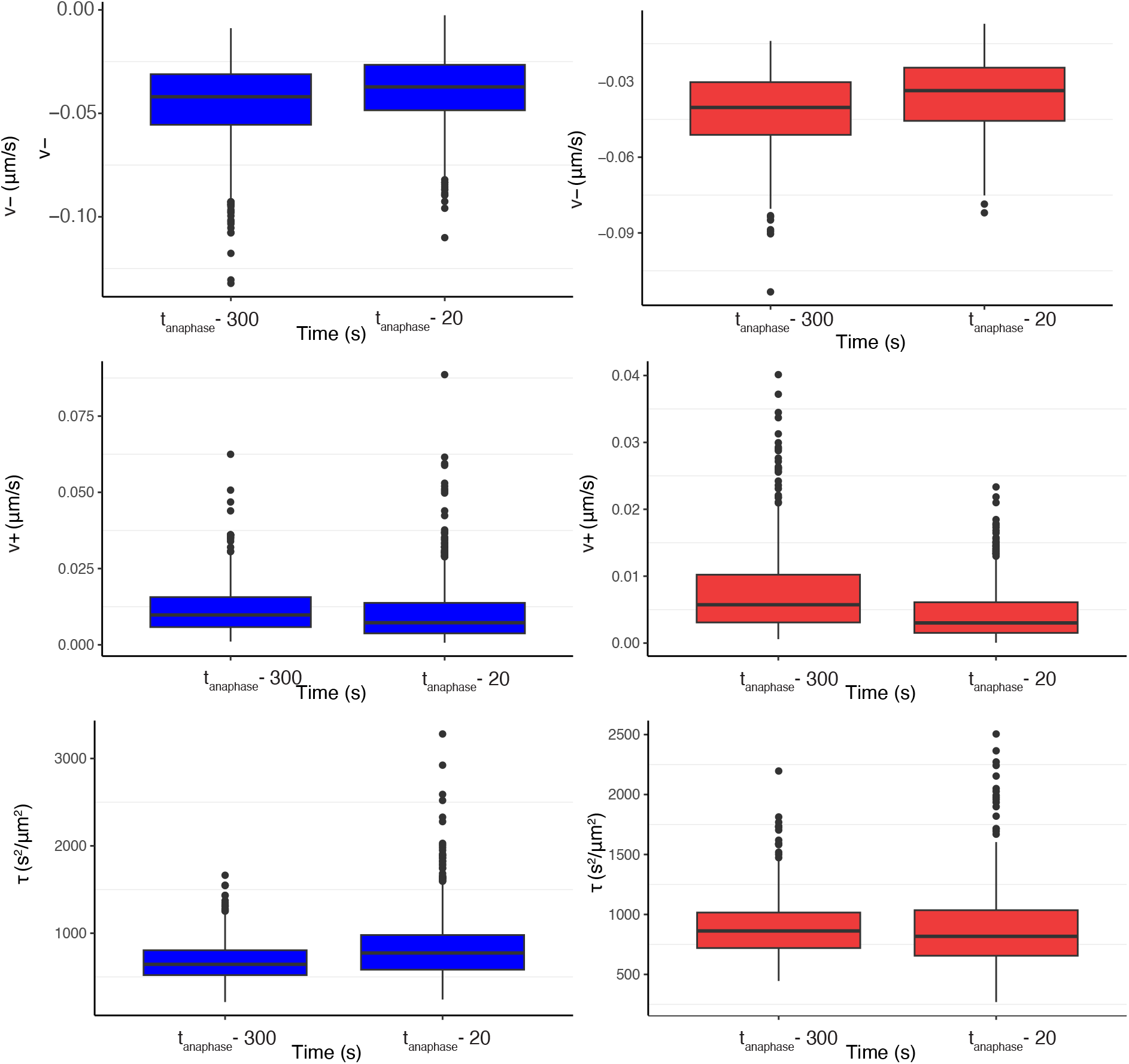
Maturation of dynamic parameters from mid-metaphase (300 s before anaphase) and just before anaphase onset (20s before anaphase). Box plots of the posterior estimation of *v*_−_(*t*) = *v*_−0_ exp {*v*_−1_*t*}, *v*_+_(*t*) = *v*_+0_ exp {*v*_+1_*t*}, *τ* (*t*) = *τ*_0_ exp {*τ*_1_*t*}, evaluated at time *t* set to 300 seconds and 20 seconds prior to anaphase, *i*.*e*., *t* = *t*_*anaphase*_ − 300 and *t* = *t*_*anaphase*_ − 20. DMSO (blue) and nocodazole washout (red) treated cells.

**Figure B13:**
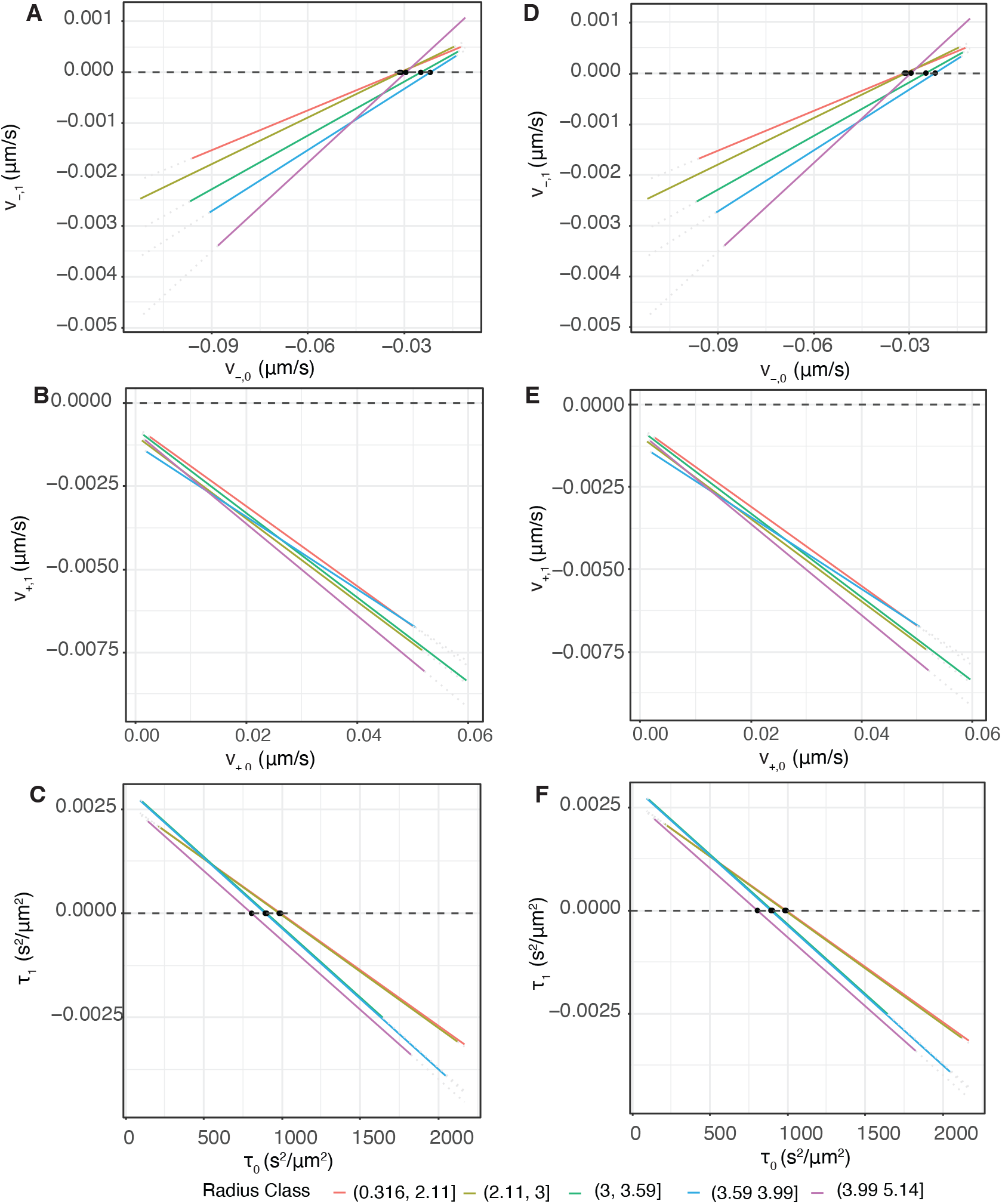
Tuning is consistent across the MPP. Plots of the regression lines for the KT pairs belonging in each of the 5 radius classes (with the five groups defined by their radial distances, *i*.*e*., [0, 2.11], (2.113], (3, 3.59], (3.59, 3.99], (3.99, 5.14] respectively) estimates of initial *p*_0_ with time dependent parameter *p*_1_. **A/D** Pulling forces *v*_−_ in DMSO, nocodazole washout treated cells, respectively. **B/E** Pushing forces *v*_+_ in DMSO, nocodazole washout treated cells, respectively.**C/F** Noise *τ* in DMSO, nocodazole washout treated cells, respectively.

**Figure B14:**
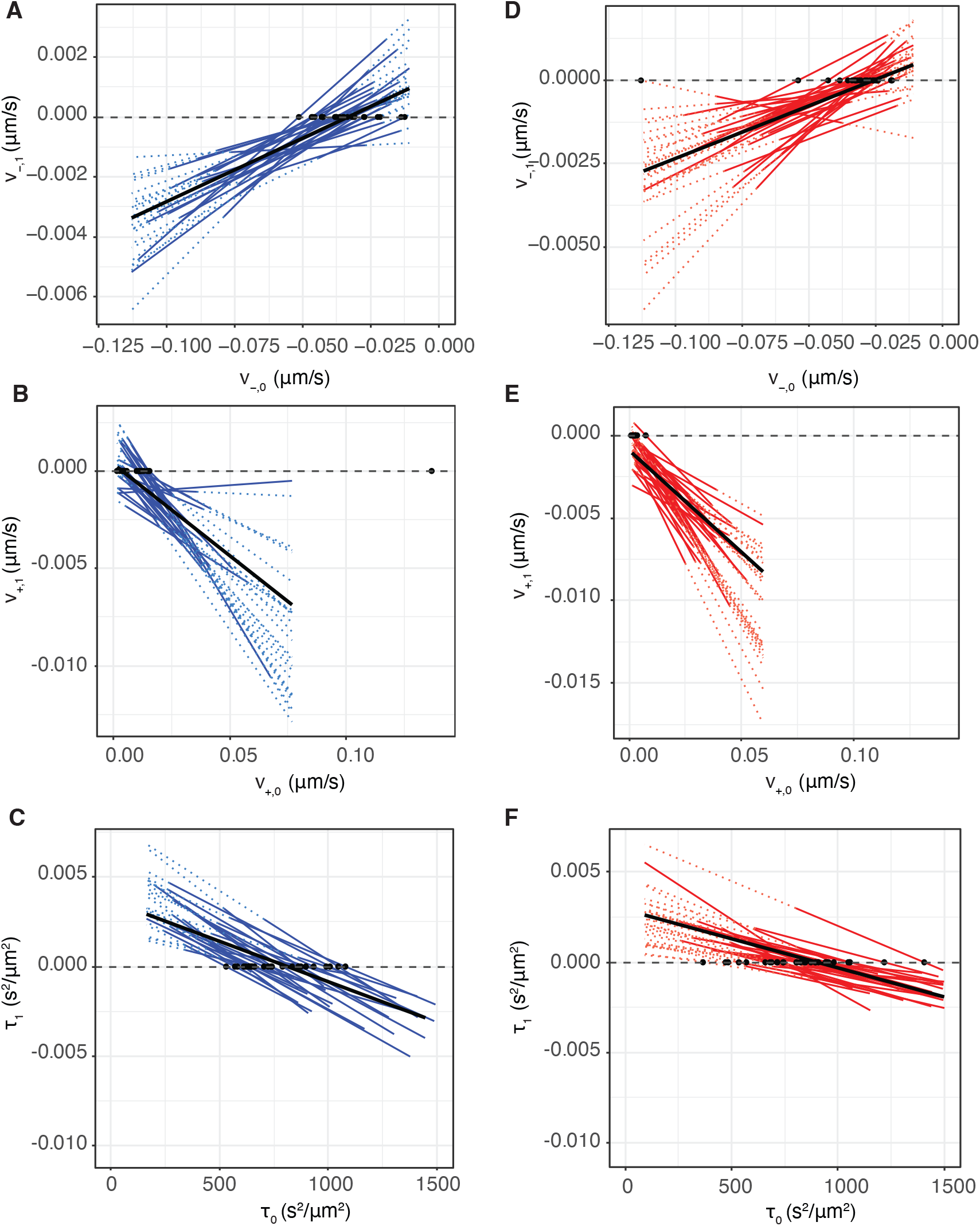
Tuning occurs in individual cells. Plots of the regression lines for individual cells of KT pair estimates of initial *p*_0_ with time dependent parameter *p*_1_. **A/D** Pulling forces *v*_−_ in DMSO, nocodazole washout treated cells. 92.3% and 83% of cells have physical (negative) anaphase ready state estimates (intercept of regression line with *v* _−1_ = 0, **B/E** pushing forces *v*_+_ in DMSO, nocodazole washout treated cells, 73.1% and 67.7% of cells have physical (positive) anaphase ready state, **C/F** precision *τ*, in DMSO, nocodazole washout treated cells,respectively, both datasets have 100% positive intercept at *τ*_1_ = 0.

**Figure B15:**
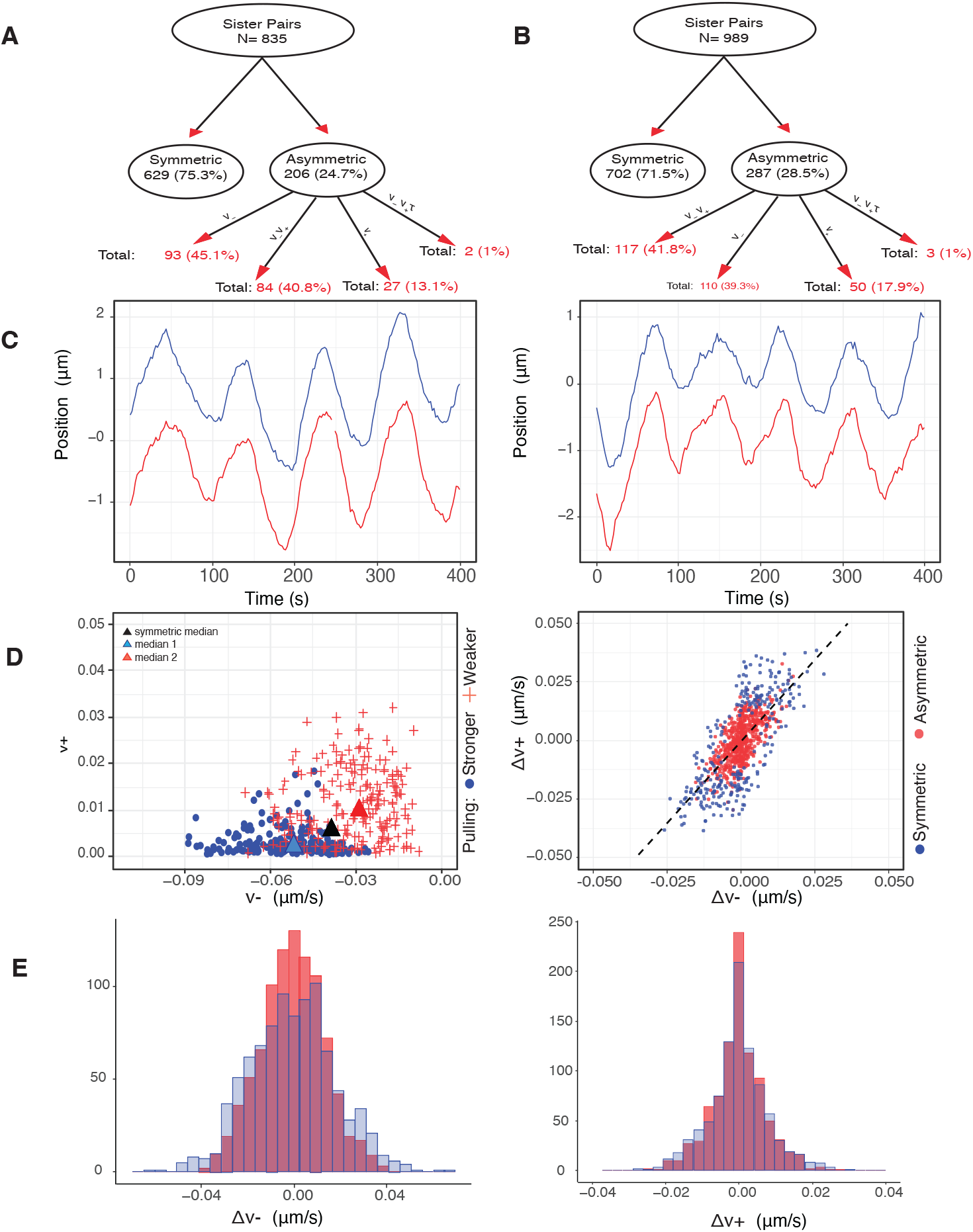
Asymmetry in sister kinetochores is unaffected by nocodazole washout treatment. **A/B** Graphical representation of asymmetry model preference network in DMSO (A) and nocodazole washout (B) treated cells. **C** Typical symmetric (left) and asymmetric (right) trajectory. No distinctive characteristics between the symmetric and asymmetric sister pairs, nocodazole washout treated. **D** (LEFT) Comparison of inferred *v*_−_ and *v*+, using the asymmetric on *v* _−_, *v*_+_ model, over the nocodazole washout asymmetric pairs. Distinction of sisters within a pair is based on their pulling force strength. Stronger and weaker pulling sisters are denoted with blue and red. Population medians are marked with triangle, while black triangle shows the estimated population median of the symmetric pairs.(RIGHT) Direct comparison of 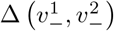 versus 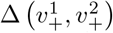 of symmetric and asymmetric nocodazole washout treated sister pairs, in red and blue respectively, when inferring them with the *v*_−_, *v*_+_ model. Asymmetric pairs do not present any distinctive pattern. **E** Histograms of the nocodazole washout sister difference distribution, *i*.*e*., 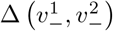 (left) and of 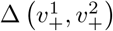 (right), when accounting for the sister pairing (red) and disturbing randomizing the pairing (blue), while accounting for the spatial metaphase plate distance.

**Figure B16:**
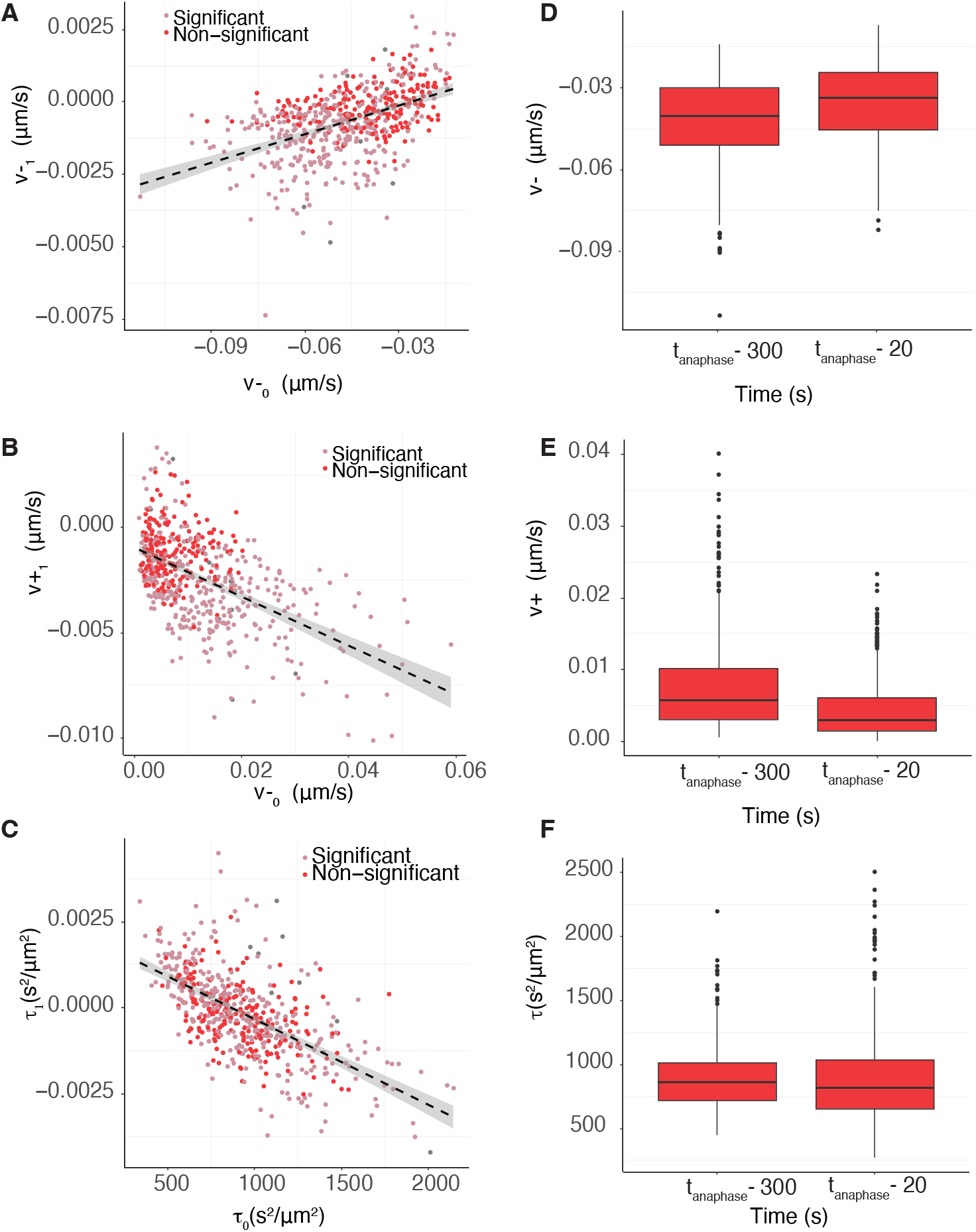
Mechanical tuning in nocodazole washout treated cells. **A/B/C** Scatter plot of posterior median of time dependence *p*_1_ versus initial parameter value *p*_0_ for **A** pushing forces, *v*_1,+_ versus *v*_0,+_; **B** pulling forces, *v*_1,_ versus *v*_0, −_; **C** *τ*_1,+_ versus *τ*_0,+_. **D/E/F** Boxplots of posterior median parameter at the beginning and the end of the trajectory for **D** pushing forces *v*_+_; **E** pulling forces *v* _−_; and **F** noise *τ*. Parameters *v* _−_, *v*_+_ are statistically different over time (mid-metaphase to late-metaphase comparison), (*p*_*MW*_ *<* 10^−16^, *p*_*MW*_ *<* 10^−9^, *p*_*MW*_ *<* 10^−16^ for *v* _−_, *v*_+_, respectively), while *τ* is not found to be statistically different over time. Finally, variances over time are statistically different with (*p*_*BF*_ *<* 10^−17^, *p*_*BF*_ *<* 10^−35^, *p*_*BF*_ *<* 10^−3^ for *v*_−_, *v*_+_, *τ* respectively). Parameters are inferred on the *M*_*v*± *τ*_ model.

**Figure B17:**
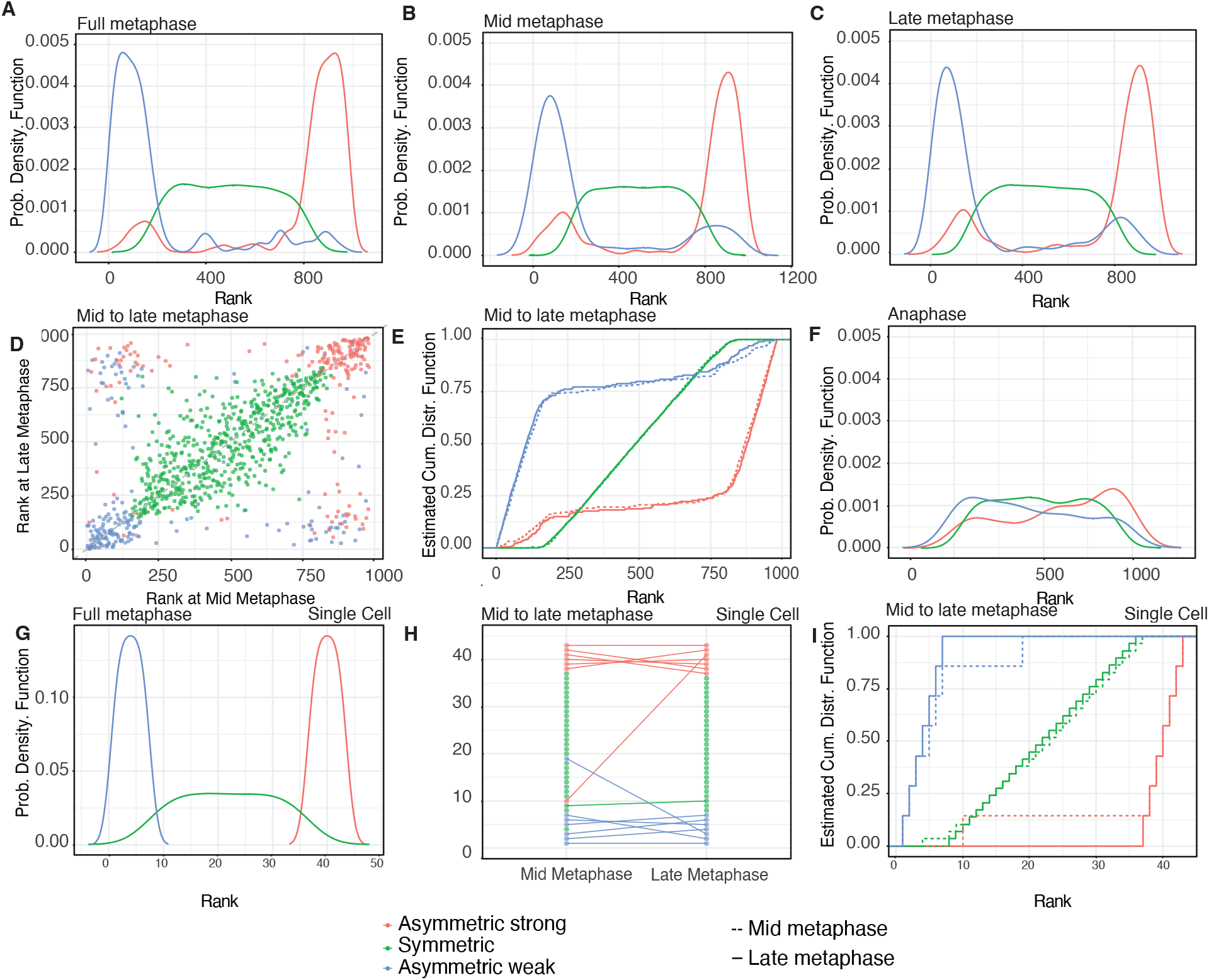
A rank analysis of the lateral organisation of the MPP in DMSO cultured cells. **A/B/C/F** Estimated probability density function of the ranks of averaged positions (from the centre of the metaphase plate) of DMSO treated kinetochores over different periods. **A** full metaphase (185 frames), **B** mid metaphase (80 frames), **C** late metaphase (80 frames) and **F** anaphase (80 frames). **D/E** Change of rank of the averaged positions from mid to late metaphase, for DMSO treated cells, averaged over 50 frames. **D** Scatter plot, **E** estimated cumulative distribution. **G** Estimated probability density function of the ranks of the averaged positions (from the centre of the metaphase plate) of kinetochores over the whole metaphase (185 frames) for a single cell. **H** Rank changes of the positions of kinetochores in a cell, from mid to late metaphase. Lines connecting points from mid to late metaphase indicate the rank of a particular sister KT at these two periods, for the same cell as in G. **I** estimated cumulative distribution of rank changes from mid to late metaphase for the same cell as in G. Red colour denotes the stronger pulling asymmetric sisters, blue the weaker pulling asymmetric sisters and green the symmetric sisters.

**Figure B18:**
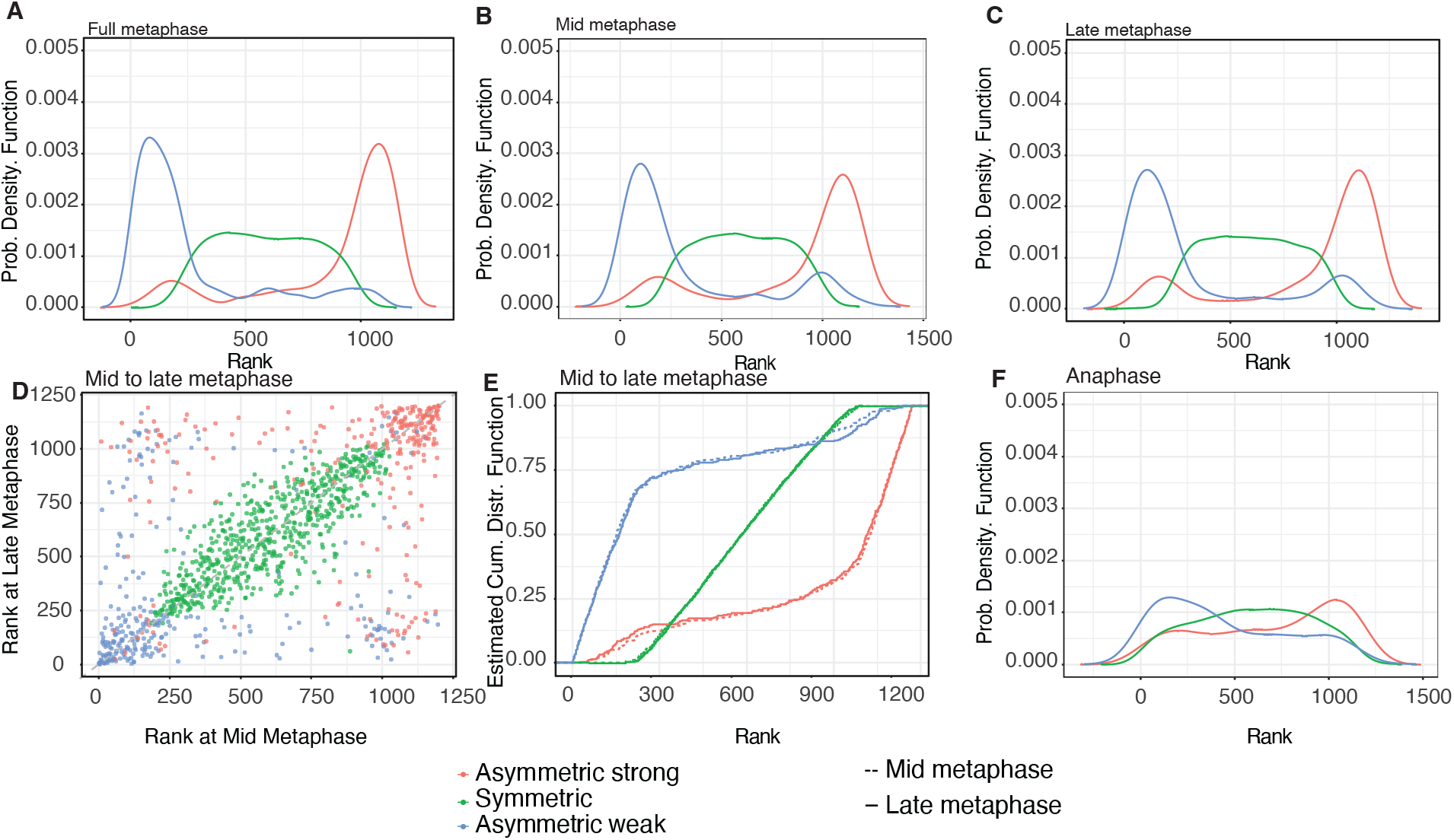
A rank analysis of the lateral organisation of the MPP in nocodazole washout treated cells. **A/B/C/F** Estimated probability density function of the ranks of averaged positions (from the centre of the metaphase plate) of nocodazole washout kinetochores over different periods. **A** full metaphase (185 frames), **B** mid metaphase (80 frames), **C** late metaphase (80 frames) and **F** anaphase (80 frames). **D/E** Change of rank of the averaged positions from mid to late metaphase, averaged over 50 frames. **D** Scatter plot, **E** estimated cumulative distribution. Red colour denotes the stronger pulling asymmetric sisters, blue the weaker pulling asymmetric sisters and green the symmetric sisters.

**Figure B19:**
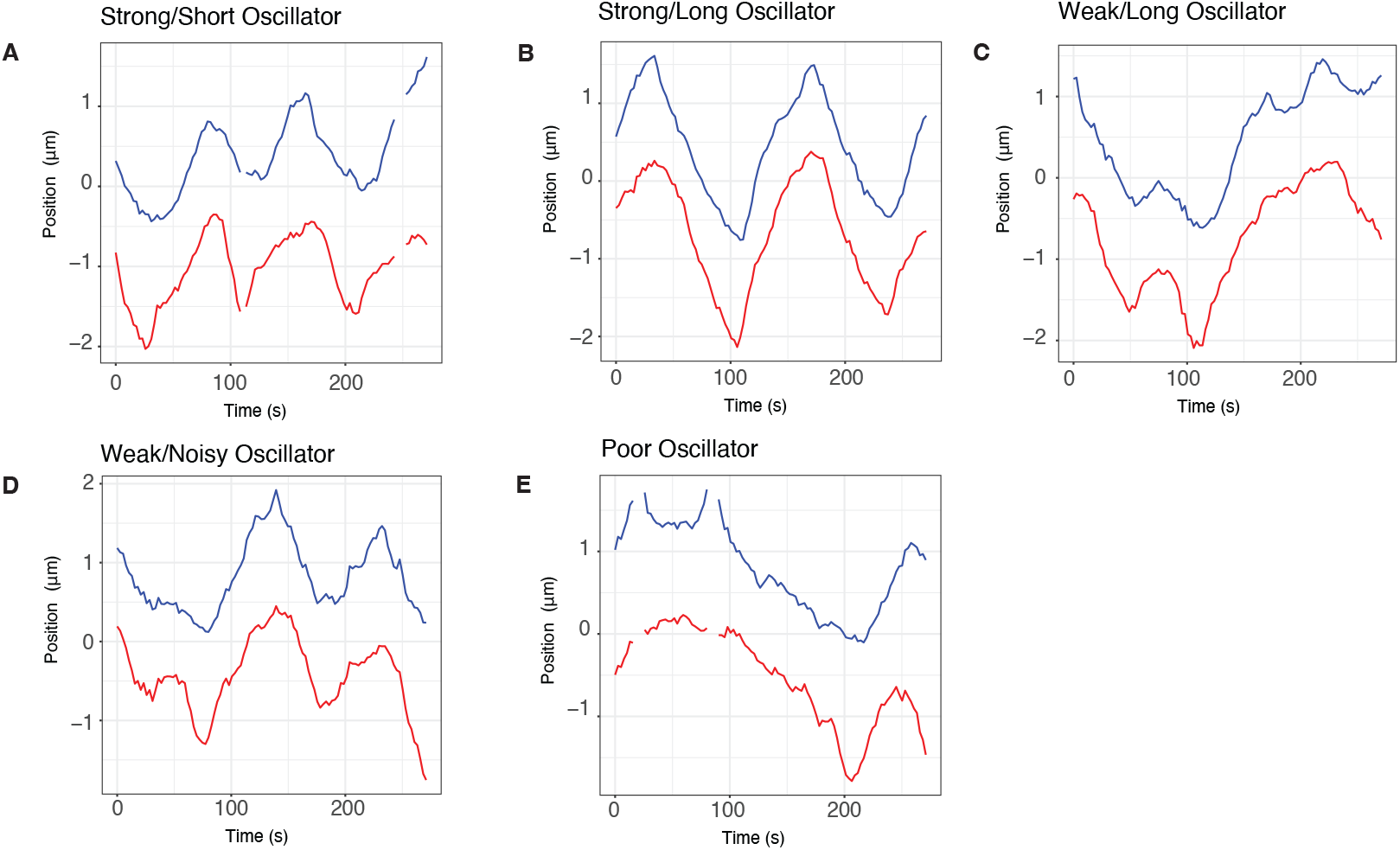
Examples of KT pair trajectories for each of the five oscillator categories: strong oscillators with short period, strong oscillators with long period, weak oscillators with long period, weak and noisy oscillators and poor oscillators/non-oscillating KT pairs.

**Figure B20:**
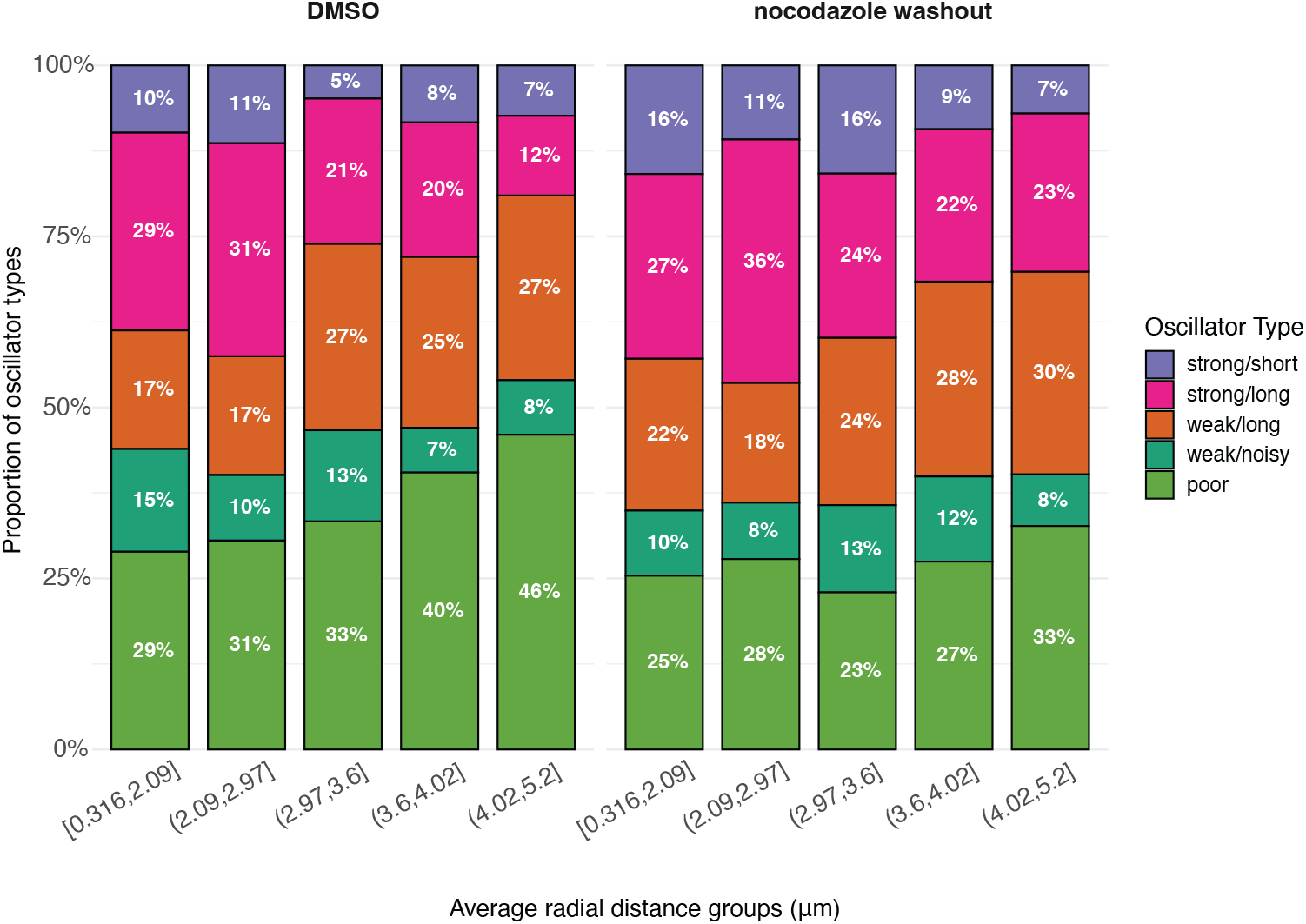
Oscillator type profile shifts from strong to poor as distance from the centre of the MPP increases. Proportion of KT pairs in the 5 oscillator clusters for the shown radial partitions.

**Figure B21:**
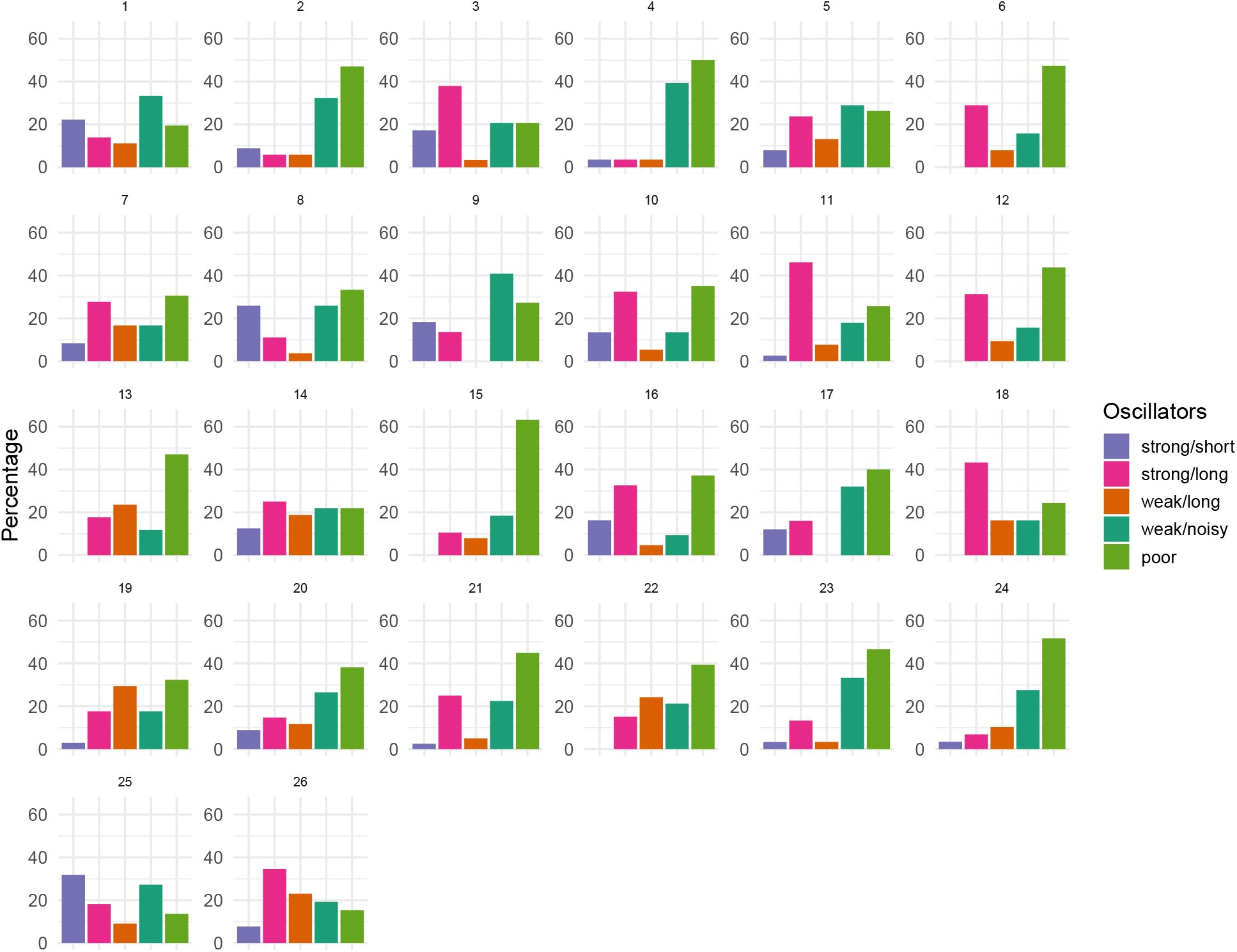
Different oscillator types across each of the 26 DMSO treated cells.

**Figure B22:**
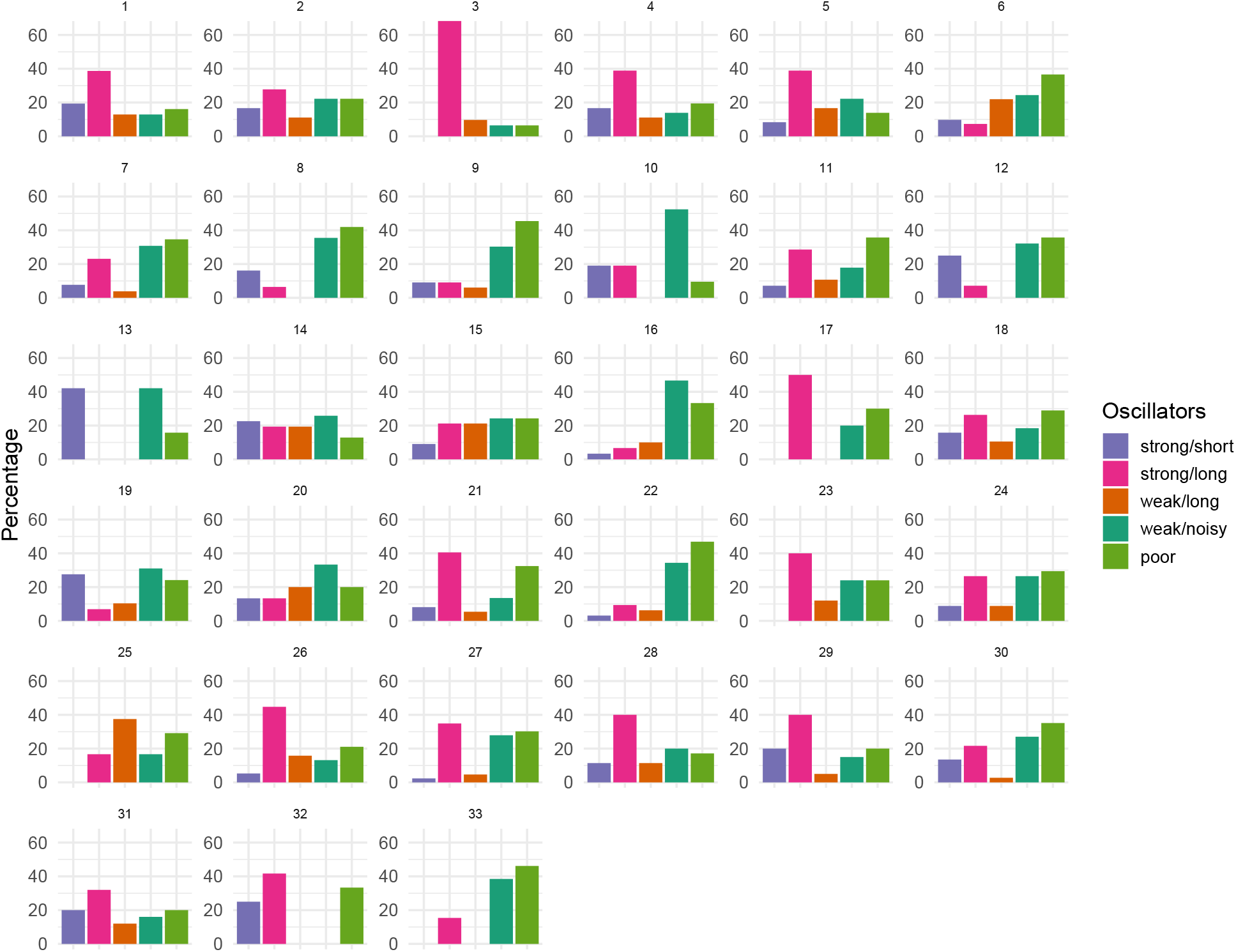
Different oscillator types across each of the 33 nocodazole washout treated cells.

**Figure B23:**
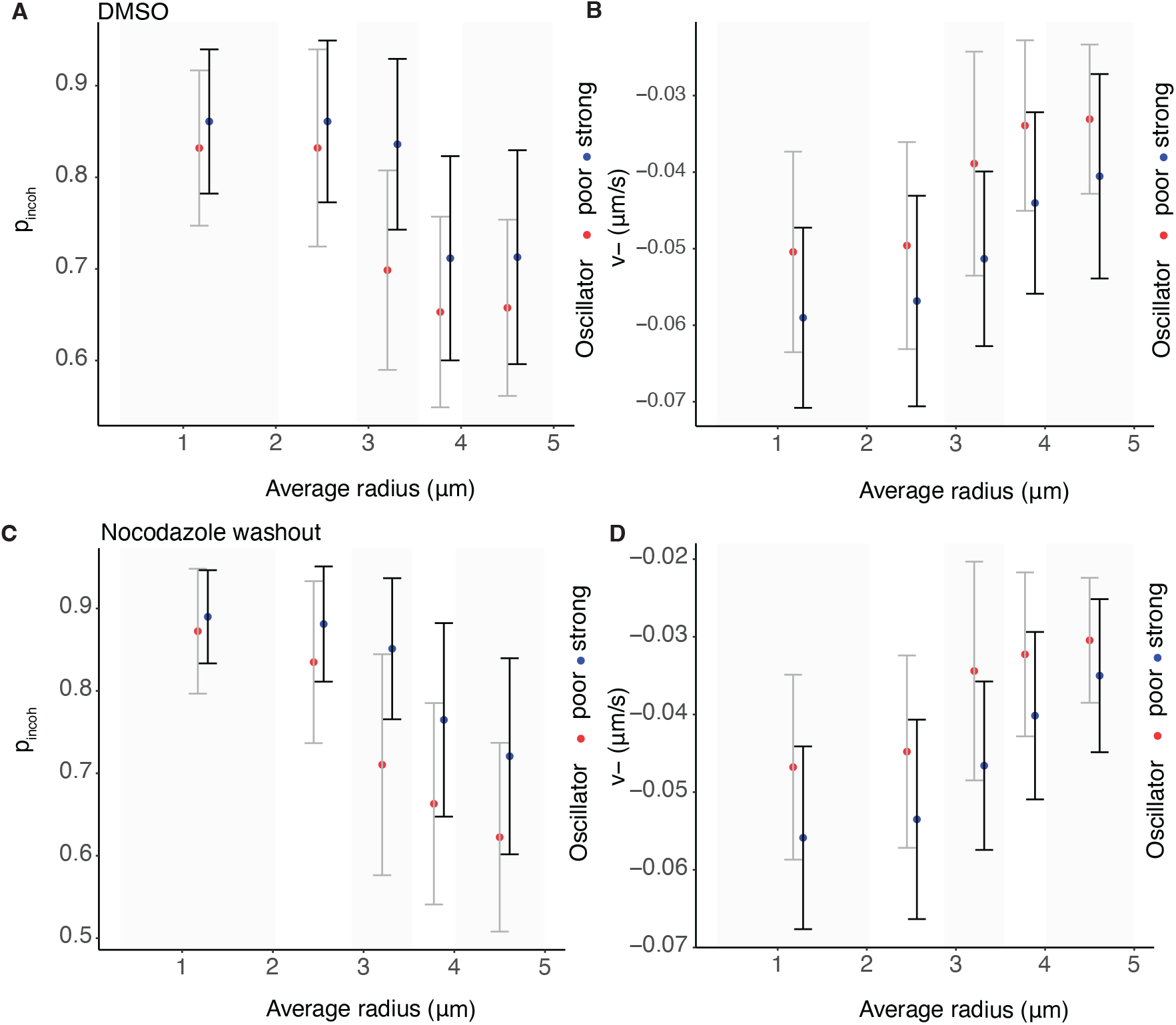
Spatial trends of poor and strong oscillators for DMSO and nocodazole washout treated cells. Parameter estimates for the asymmetric 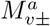 model for *v*_−_ and the probability of remaining incoherent between frames (*p*_*incoh*_).

**Figure B24:**
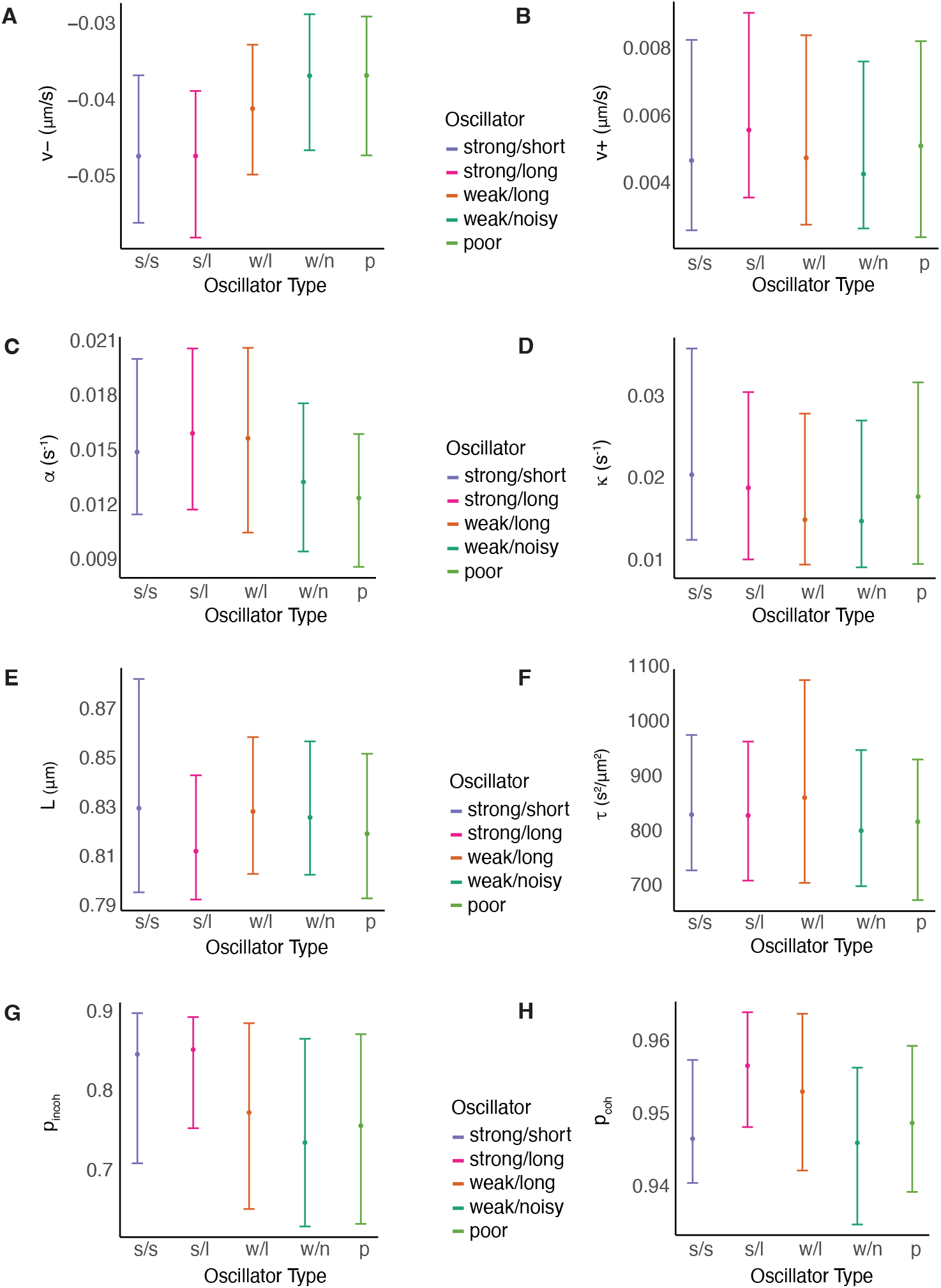
Biophysical parameters with oscillation quality (nocodazole washout). Parameter estimates for the asymmetric 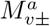 model, are consistent across different types of oscillators in nocodazole washout treated cells, except *v*_−_ and the probability of remaining incoherent between frames (*p*_*incoh*_), where we observe a decreasing trend as the quality of oscillation deteriorates.

**Figure B25:**
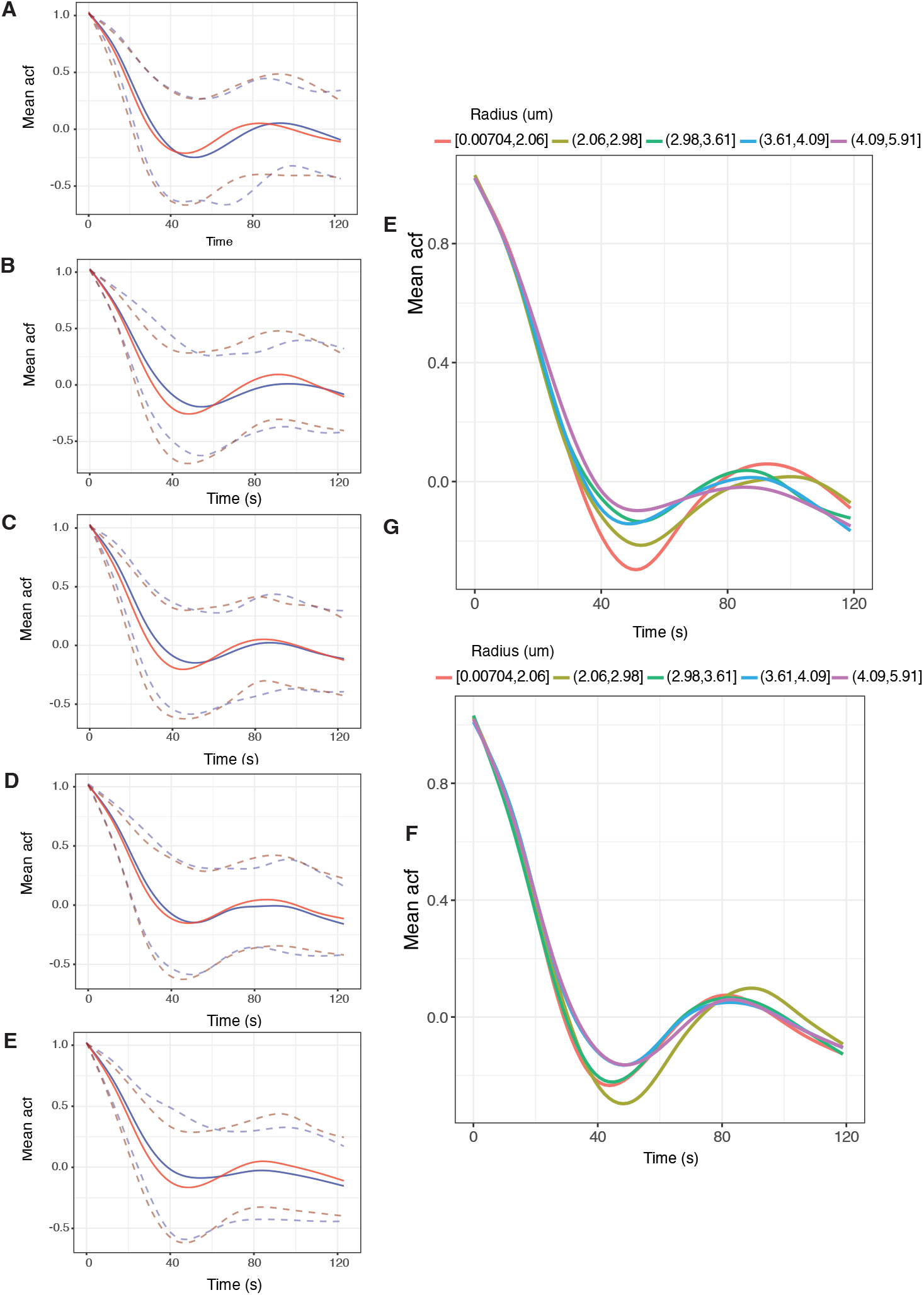
Autocorrelation plot analysis across the metaphase plate: **A-E** Autocorrelation plots for DMSO (blue) and nocodazole washout (red) treated cells. Plots A to E present the mean autocorrelations for sister pairs that belong in one of five groups defined by their radial distances, *i*.*e*., [0, 2.07], (2.07, 2.97], (2.97, 3.59], (3.59, 4.01], (4.01, 5.20] respectively. Dashed lines denote the 5% and 95% autocorrelations percentile for each time point. **E, F** Summary of mean autocorrelation plots for DMSO and nocodazole washout treated cells for all 5 classes of radial distances.

### B. Additional plots

### C Methodology

#### C1 Data processing and Quality Control

We analyse mitosis in immortalised human retinal pigment epithelial cells (RPE1), a karyotypically stable, non-transformed cell line, using Lattice Light Sheet Microscopy (LLSM, [Chen et al., 2014]). We achieve a high signal-to-noise ratio and a temporal resolution of 2s/frame; this enables analysis of mitosis dynamics in a normal human cell line whilst ensuring minimal photobleaching and phototoxicity (all our cells transition to anaphase). We demonstrate near-complete 3D tracking of the 46 kinetochore pairs for up to 15 mins, and parametrise the models by fitting to KT trajectories.

Kinetochore tracking is performed using the software package KiT v3.0. The tracking algorithm proceeds by detecting candidate spots via a constant false alarm rate (CFAR) detection method, [Daniyan et al., 2024], to set a KT-wise dynamic threshold per image frame in a movie. Candidate spot locations are refined by fitting a Gaussian mixture model. Spot locations are linked between frames by solving a linear assignment problem using the Hungarian algorithm, with motion propagation via a Kalman filter. A “ track” refers to a temporally ordered set of KT positions (coordinates) associated with a single KT across sequential image frames, constructed by linking detections frame-to-frame. A “ gap” is defined as one or more consecutive frames within a track during which the KT was not detected; tracks are permitted to contain gaps of up to 3 frames, after which the track is considered terminated.

Tracked kinetochores are paired by solving a second linear assignment problem. Sister kinetochore pairing used spatial and temporal trajectory data. Sister kinetochores are closer (*d*_*ij*_, average distance) and exhibit less distance variation (*v*_*ij*_, distance variance) than non sisters. In the presence of a metaphase plate, their connecting vector aligns with the plate’s normal (*α*_*ij*_, average angle). Pairing costs were calculated as the product *d*_*ij*_*v*_*ij*_ without a plate or *d*_*ij*_*v*_*ij*_*α*_*ij*_ with a plate. Trajectories had to overlap for at least 10 frames, and the pairing configuration with the minimum global cost was selected.

#### Detection and tracking was near complete

The tracklet plot, Figure 1C, shows that 82 KTs were tracked throughout the movie, whilst a total of 100 tracks were recorded (tracks are allowed to have gaps of at most 3 missing frames). Additionally, 43 KT pairs were tracked for at least 75% of the movie, with 38 KTs remaining paired for the entire duration of metaphase (sisters are allowed gaps of at most 3 frames), Figure 1D,E. The scatter plot of duration of pairing against duration either sister exists demonstrates near complete pairing over the duration of KT detection, Figure 1F. The average sister track length for this movie was 705 s. Spot detection and tracking performance are unaffected by the transition to anaphase.

36 RPE1 cells were imaged during metaphase-anaphase of varying durations in DMSO. A quality control filter was applied to ensure good sister pair coverage per cell. Specifically, at least 30 sister pairs had to be tracked for 75% of the movie length, resulting in 31 cells meeting the criteria and 1281 sister pair kinetochores being tracked. On average, 40 sister pairs per cell were tracked (quartiles: Q1 = 38.5, Q3 = 43), with both sisters tracked for at least 200 seconds (100 frames).

Prior to fitting the models, we implemented a data quality control. We discarded cells with a low coverage of sister pairs or a very high proportion of poor oscillators (indicating poor health), and for the metaphase-anaphase model, if there was insufficient anaphase. This reduced the analysed cells from 36 DMSO treated cells to 28 cells. We also filtered KT-pair trajectories, requiring no more than 20% missing data, and cell videos are required to have at least 120 metaphase frames. Trajectories with poor inference, identified by severe divergences in STAN, were excluded, and cells were discarded if a high proportion of sister pair trajectories showed frequent divergences, resulting to a further reduction of the analysed cells to 26. While divergences indicate the model isn’t fully capturing the dynamics, they were rare (∼1.5%, with slight variations between the different models). Hence, in the model analysis there are slight differences in the number of trajectories considered due to performance issues. For the metaphase dynamics models (that don’t include the anaphase transition), we truncated the trajectories to metaphase. Specifically, we used a change point model to detect the anaphase transition per KT pair, Appendix E. Then we truncated 30s before this, to ensure the cell is in metaphase. For nocodazole washout treated data, we have initially tracked 51 cells. Applying our filtering criteria of sister pair coverage, poor oscillators and sufficient anaphase part as in DMSO treated cells, we ended up with 33 nocodazole washout treated cells.

#### C2 Likelihood and posterior distribution computation

The metaphase dynamics of eq. (1), are described in detail in Armond et al. [2015a]. The model is a discretised system of equations – discrete measurement time with equal time steps:

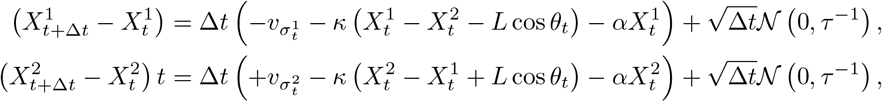

The kinetochore pair K-fiber state, *σ*_*t*_, (hidden), takes values in the 4 states,

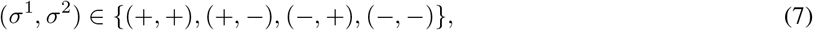

and has transition matrix *P* (parametrised by *p*_*coh*_, *p*_*icoh*_, the probabilities of remaining in the same state (either coherent or incoherent).

In sum, the observations obey a Stochastic Differential Equation (SDE), with the observation at time *t* depends only on the previous observation *x*_*t* −1_ and the state at time *t*, while the hidden states are described by the transition probability matrix.

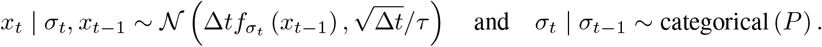

with

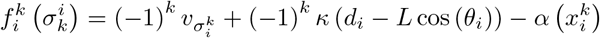

The likelihood of each sister is given by

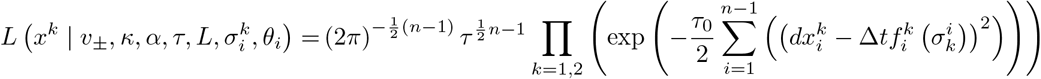

This model can be modified accordingly to allow for asymmetry, as in eq. (3), and time dependency, as in eq. (5). The likelihood calculation for the asymmetric model is straightforward and hence is omitted. However, while the generalization to asymmetric models is relatively straightforward, the increased complexity lead us to use the following transformations for velocities

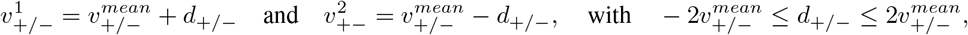

which improve the mixing of the Hamiltonian Monte Carlo algorithm. The likelihood for the full time dependent model is shown below.

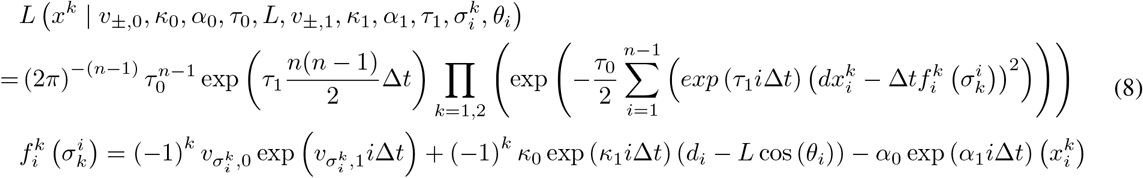

#### C3 Forward - Stochastic Backward Algorithm

We leverage a modified Forward-Backward (FB) algorithm, [Scott, 2002], to calculate the likelihood and make inference on the parameters. In particular, we use the Forward - Backward stochastic recursion within the STAN environment, [Stan Development Team, 2024a].

The FB algorithm is designed to exploit via recursion the conditional independencies in the HMM. The forward likelihood is defined as *l*_*t*_ (*i*) = *P* (*x*_1:*t*_, *σ*_*t*_ = *i* | *θ*), which is the joint likelihood of the data *x*_1:*t*_ and the state *σ*_*t*_ = *i* averaging over the states up to time *t* − 1. The likelihood contribution from data *x*_1:*t*_ is just the sum 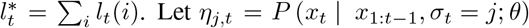, be the likelihood contribution of an observation, *x*_*t*_ under all states *j, P* (*σ*_*t*_ = *i σ*_*t* 1_ = *j*) the probability that at time *t* the state will be *i* given at *t* − 1 was at state *j*, as given by the HMM transition probability matrix and *θ* all the unknown parameters. A recursive relationship can be derived as follows

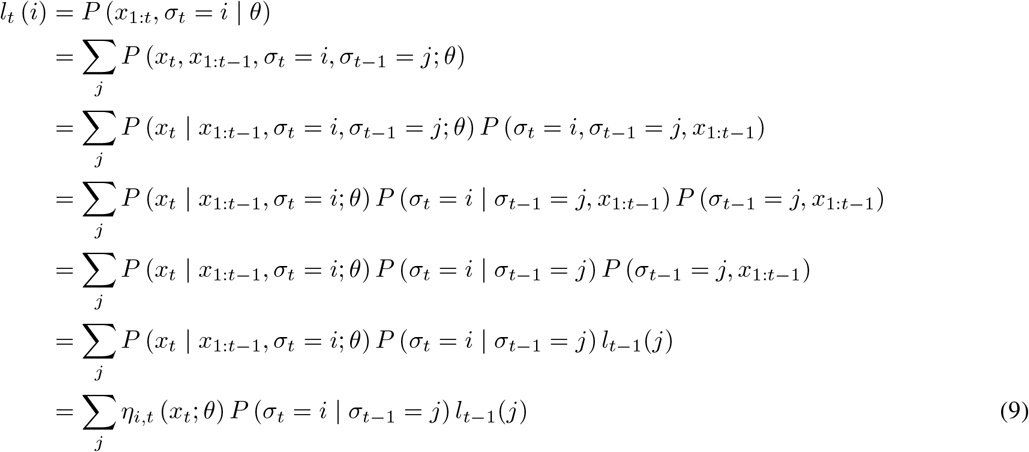

It is then obvious that the log-likelihood is the sum of all states at time *t* = *T*, *i*.*e*.,

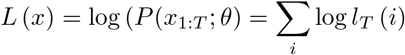

In our codes, we are using log-likelihoods for all our calculations as they are more numerically stable.

Now, let *ξ*_*j*,*t*_ = *P* (*σ*_*t*_ = *j* | *x*_1:*t*_; *θ*) be the probability of being in state *σ*_*t*_ at time *t* given all the data up to that time. Taking as initial condition *ξ*_0_ = *P* (*σ*_0_ = *j* | *θ*) = [0, 0.5, 0.5, 0]^*T*^, *i*.*e*., at time *t* = 0 it is equally probable to be in one of the coherent states (+−, −+). Observe that

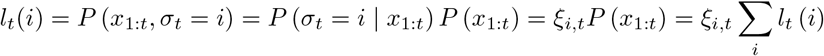

and note that *f*_*t*_ = *P* (*x*_*t*_ | *x*_1:*t*−1_) = ∑_*i*_ ∑_*j*_ *η*_*i*,*t*_ (*x*_*t*_; *θ*) *P* (*σ*_*t*_ = *i* | *σ*_*t*−1_ = *j*) *ξ*_*j*,*t*−1_ (*i*) then eq. (9) can be modified and calculate the forward probabilities *ξ*_*i*,*t*_ = *i* _*l*_*t*_(*i*)_ instead and derive our final recursive relationship

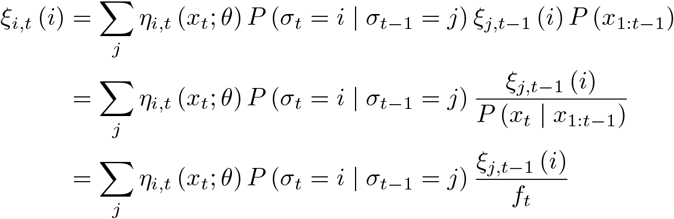

and likelihood is

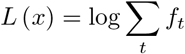

In our setting, we have

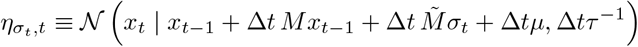

where *M*, 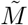, and *µ* are derived from the linear SDE and given by

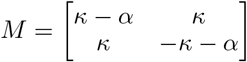

and

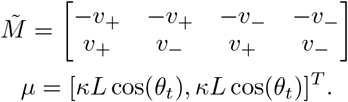

In this notation, the state variable 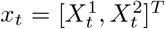 represents the positions of both sisters in a pair at time *t* and the state *σ*_*t*_ ∈ { [1, 0, 0, 0]^*T*^, [0, 1, 0, 0]^*T*^, [0, 0, 1, 0]^*T*^, [0, 0, 0, 1]^*T*^} corresponding to the states {++, +−, −+,−−} for the sister kinetochore pair at time *t*.

In order to make statements about switching events, we need to consider sequences of states forming a pattern corresponding to coherent switches from one coherent state to another via an intermediate state. To address this, we sample from the full hidden state sequence given all the data, and assess this for switches. We use the stochastic backward simulation to simulate/sample the hidden states, which is a rapidly mixing algorithm, [Scott, 2002]. Observe that

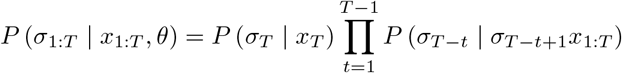

Then, keeping the terms that depend on *σ*_*t*_, the it is easy to show the following

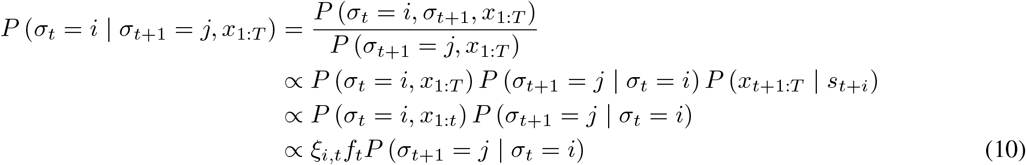

The strategy for the backward sampling algorithm is therefore to sample initially from *p* (*σ*_*T*_ |*x*_1:*T*_) = *ξ*_*T*,*i*_ as dervived in the forward recursion and subsequently to simulate backward in time from *T*− 1 to 1 via the conditional distribution given in eq. (10).

#### C4 Missing data treatments

Suppose we have missing observations. Let’s assume that the data comprises two sections [1 : *T*_1_] and [*T*_2_ : *T*], with *T*_2_ = *T*_1_ + 1 + *l > T*_1_, *i*.*e*., a gap of size *l* ≥1 where data on both sisters is missing. Given the Markovian property of the observations, we can construct the multi-time point conditional, *P* (*X*_*t*+*k*Δ*t*_, *σ*_*t*+*k*Δ*t*_ |*X*_*t*_, *σ*_*t*_, *θ*). Incorporating the dependence on the hidden state path, we get the following expression,

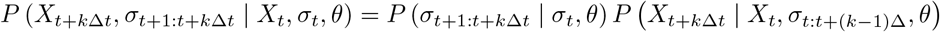

Thus, we have the full history dependency *P* (*X*_*T*_2 | *X*_1:*T*_1) across a gap.

We consider then obtain the likelihood,

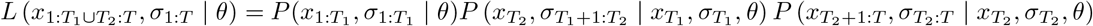

and the marginalised likelihood 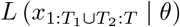 given by summation over the hidden state *σ*_1:*T*_. There are a lot of different methods that we can utilise that deal with the gap (middle) terms,

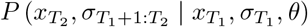

and each comes with its own set of strengths and weaknesses.

In brief, there we have four options. First, we could fill in the missing values with linear interpolation, secondly, treat sections as independent, *i*.*e*., multiply the likelihoods for each section, thirdly, remove data points from the likelihood calculation but retain hidden state propagation, fourthly, impute the hidden values by sampling from the posterior predictive distribution

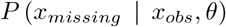

(using a Brownian Bridge sampler). The first two methodologies are approximating the actual time-series, as if there were no missing values and thus, they introduced a bias which can be, in some cases, non-negligible. The third method, doesn’t utilise all the available information, while the fourth option – even though is the most flexible option for extending to missing values of a single sister – it is likely computationally expensive for large gaps. Taking into account the advantages and limitations of each method, we have chosen to work with the third method, which we have found that it optimally balances the trade-off between reducing computational complexity and introducing estimation bias.

#### C5 Reduced likelihood method

Consider the full likelihood. The gap term can be expressed as,

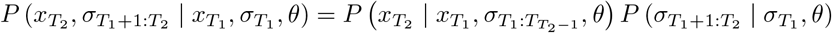

The idea here is to exclude the propagation term for *x*_*T*_2, i.e, use the reduced full likelihood

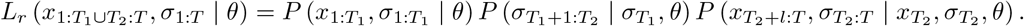

giving the corresponding reduced marginalised likelihood 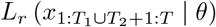 (here we removed 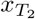 in notation as information in knowing 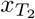 is not used. However, the likelihood depends on 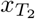, as it is part of the history).

We *propose* to modify the forward algorithm, based on the above lines. In particular, we change the forward likelihood calculation by assuming that the likelihood contribution of the missing observations is 0 and hence, the likelihood contribution at that specific time point, is only defined by the transition probabilities.

We have made a series of experiments, to proof that this methodology is robust and gives unbiased results. In particular, we have compared the likelihood estimations of different simulated data sets, with varying missing data proportions and different distributed missing values (changing the size of the gaps). Also, we have compared the results of this methodology with the results of the first method,*i*.*e*., filling the missing data by linear interpolation, which gave similar results, when the missing values’ gaps were small. In Appendix Table C1 we present a case study where we havesimulated 10 different time-series of length *T* = 180 and we have assessed whether the different mean posterior estimates where greater than the actual values with probability *p* ≠ 0.5. To asses this hypothesis, we performed a two-sided binomial test. In all of our experiments the “ reduced likelihood computation” was accurate, with small estimation bias.

**Table C1:**
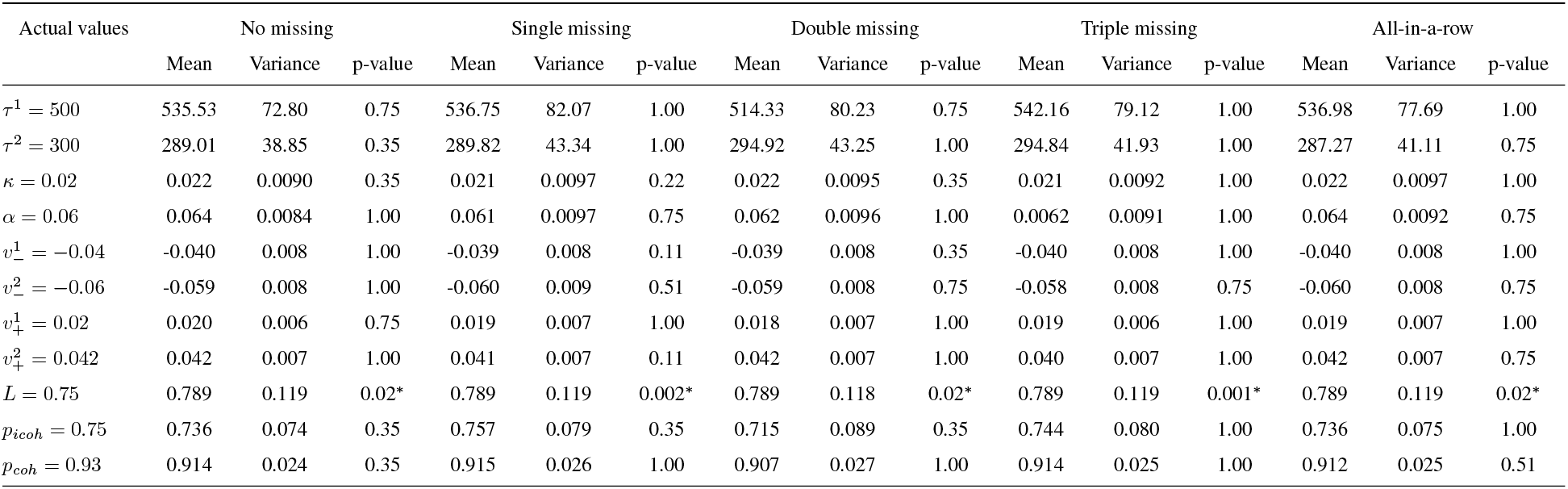
Summary of the case study for checking the bias of our algorithm when we have no missing data and 10% missing data points under various settings, *i*.*e*., single missing data points, double missing data points, triple missing data points and all-in-a-row missing data points. The case study comprises ten different simulated time-series of length *T* = 180, from a common multivariate distribution, in an effort to Average posterior mean MCMC estimates, average posterior MCMC variance and p-values of the two-sided exact binomial test presented. The null hypothesis is that there is 0.5 probability that the posterior mean estimates are greater than the actual value. Our algorithm gives unbiased results

#### C6 Prior distributions

We impose broad prior distributions on the parameters of the biophysical model, as shown in Appendix Table C2. For the natural length of the spring, *L*, we impose an informative prior based on an additional nocodazole washout experiment (Nocodazole interferes with polymerization of microtubules). This avoids an unidentifiabilty in the model as in Armond et al. [2015a]. Additionally, we use an informative prior for the time of anaphase, *t*_*A*_, based on first fitting a changepoint model (with a uniform prior on *t*_*A*_) to get an initial estimate for anaphase onset to guide the biophysical model and avoid exploring parameter space corresponding to pathological behaviour such as anaphase at the start or end of movies.

#### C7 Statistical Tests

The tests that have been used throughout this study are summarised in Appendix Table C3, describing each test and the abbreviation we use when quoting the p-values in the text.

**Table C2:**
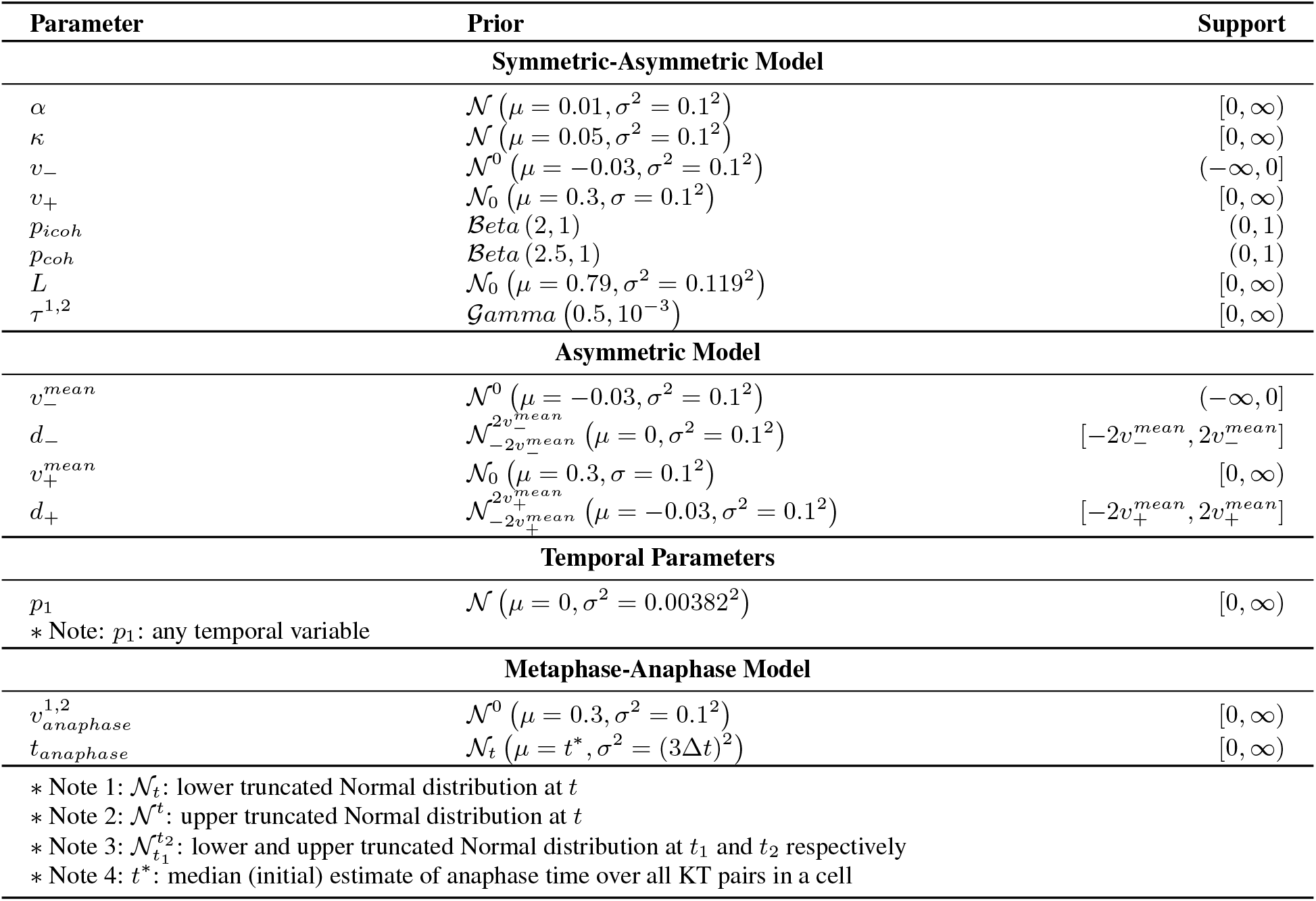
Priors for the symmetric, asymmetric, temporal, metaphase-anaphase models.

| Parameter | Prior | Support |
| --- | --- | --- |
| **Symmetric-Asymmetric Model** |  |  |
| $\alpha$ | $\mathcal{N}(\mu = 0.01, \sigma^2 = 0.1^2)$ | $[0, \infty)$ |
| $\kappa$ | $\mathcal{N}(\mu = 0.05, \sigma^2 = 0.1^2)$ | $[0, \infty)$ |
| $v_-$ | $\mathcal{N}^0(\mu = -0.03, \sigma^2 = 0.1^2)$ | $(-\infty, 0]$ |
| $v_+$ | $\mathcal{N}_0(\mu = 0.3, \sigma = 0.1^2)$ | $[0, \infty)$ |
| $p_{icoh}$ | $Beta(2, 1)$ | $(0, 1)$ |
| $p_{coh}$ | $Beta(2.5, 1)$ | $(0, 1)$ |
| $L$ | $\mathcal{N}_0(\mu = 0.79, \sigma^2 = 0.119^2)$ | $[0, \infty)$ |
| $\tau^{1,2}$ | $Gamma(0.5, 10^{-3})$ | $[0, \infty)$ |
| **Asymmetric Model** |  |  |
| $v_-^{mean}$ | $\mathcal{N}^0(\mu = -0.03, \sigma^2 = 0.1^2)$ | $(-\infty, 0]$ |
| $d_-$ | $\mathcal{N}_{-2v_-^{mean}}^{2v_-^{mean}}(\mu = 0, \sigma^2 = 0.1^2)$ | $[-2v_-^{mean}, 2v_-^{mean}]$ |
| $v_+^{mean}$ | $\mathcal{N}_0(\mu = 0.3, \sigma = 0.1^2)$ | $[0, \infty)$ |
| $d_+$ | $\mathcal{N}_{-2v_+^{mean}}^{2v_+^{mean}}(\mu = -0.03, \sigma^2 = 0.1^2)$ | $[-2v_+^{mean}, 2v_+^{mean}]$ |
| **Temporal Parameters** |  |  |
| $p_1$ | $\mathcal{N}(\mu = 0, \sigma^2 = 0.00382^2)$ | $[0, \infty)$ |
| * Note: $p_1$: any temporal variable | | |
| **Metaphase-Anaphase Model** |  |  |
| $v_{anaphase}^{1,2}$ | $\mathcal{N}^0(\mu = 0.3, \sigma^2 = 0.1^2)$ | $[0, \infty)$ |
| $t_{anaphase}$ | $\mathcal{N}_t(\mu = t^*, \sigma^2 = (3\Delta t)^2)$ | $[0, \infty)$ |
| * Note 1: $\mathcal{N}_t$: lower truncated Normal distribution at $t$ | | |
| * Note 2: $\mathcal{N}^t$: upper truncated Normal distribution at $t$ | | |
| * Note 3: $\mathcal{N}_{t_1}^{t_2}$: lower and upper truncated Normal distribution at $t_1$ and $t_2$ respectively | | |
| * Note 4: $t^*$: median (initial) estimate of anaphase time over all KT pairs in a cell | | |

**Table C3:**
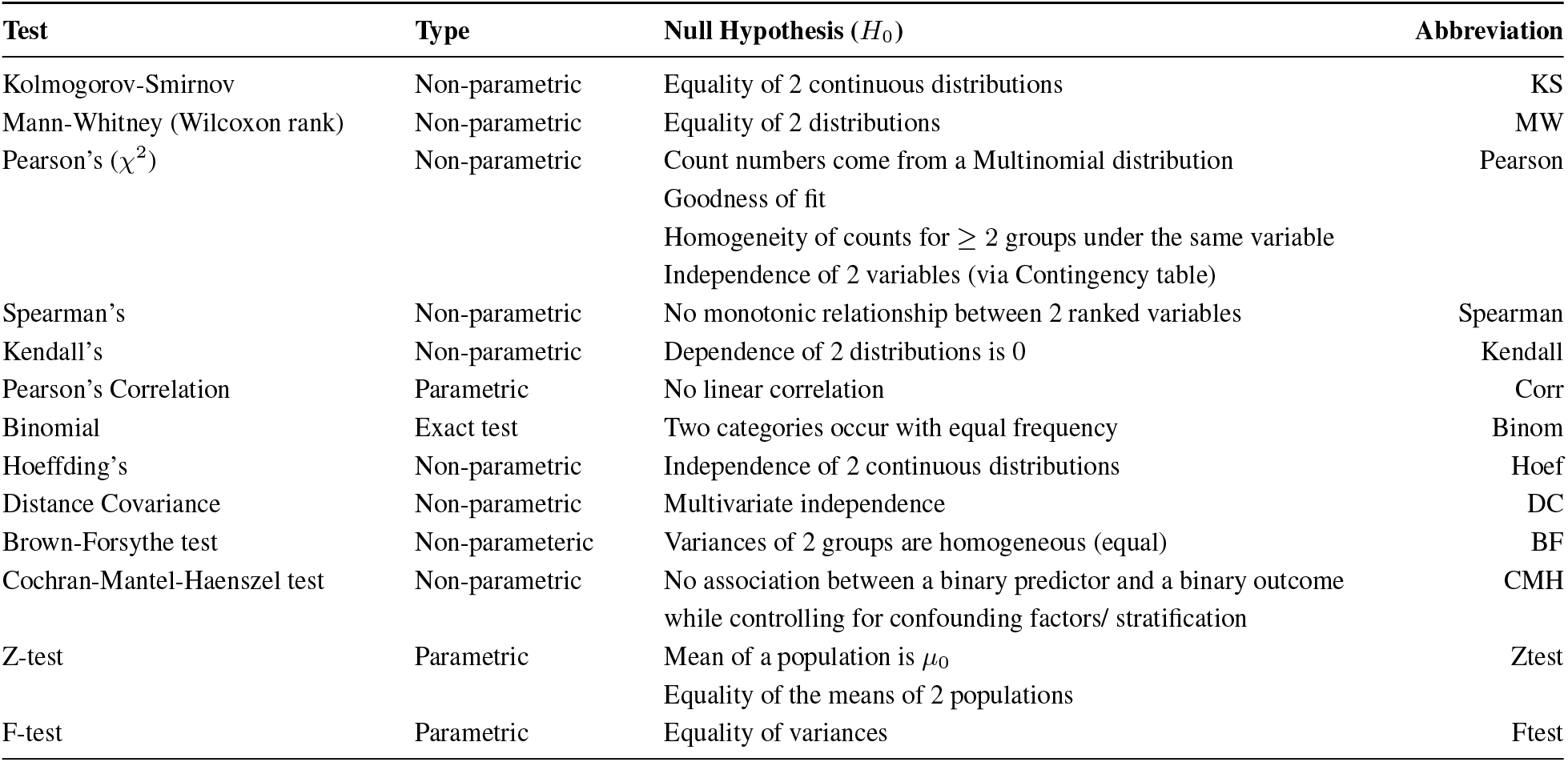
Summary of tests used in this analysis. The first column refers to the test, the second the type of the test, *i*.*e*., assumptions, the third column describes the null hypothesis and hence the test, while the last column states the abbreviation that is used throughout this study, when referring to p-values.

| Test | Type | Null Hypothesis ($H_0$) | Abbreviation |
| --- | --- | --- | --- |
| Kolmogorov-Smirnov | Non-parametric | Equality of 2 continuous distributions | KS |
| Mann-Whitney (Wilcoxon rank) | Non-parametric | Equality of 2 distributions | MW |
| Pearson's ($\chi^2$) | Non-parametric | Count numbers come from a Multinomial distribution | Pearson |
|  |  | Goodness of fit |  |
| | | Homogeneity of counts for $\geq 2$ groups under the same variable | |
|  |  | Independence of 2 variables (via Contingency table) |  |
| Spearman's | Non-parametric | No monotonic relationship between 2 ranked variables | Spearman |
| Kendall's | Non-parametric | Dependence of 2 distributions is 0 | Kendall |
| Pearson's Correlation | Parametric | No linear correlation | Corr |
| Binomial | Exact test | Two categories occur with equal frequency | Binom |
| Hoeffding's | Non-parametric | Independence of 2 continuous distributions | Hoef |
| Distance Covariance | Non-parametric | Multivariate independence | DC |
| Brown-Forsythe test | Non-parametric | Variances of 2 groups are homogeneous (equal) | BF |
| Cochran-Mantel-Haenszel test | Non-parametric | No association between a binary predictor and a binary outcome while controlling for confounding factors/ stratification | CMH |
| Z-test | Parametric | Mean of a population is $\mu_0$ | Ztest |
|  |  | Equality of the means of 2 populations |  |
| F-test | Parametric | Equality of variances | Ftest |

#### C8 Implementation

All our models were implemented in STAN, [Stan Development Team, 2024a] and fitted using the R-package “ rstan”. STAN is based on a C++ language which obtains posterior samples of the Bayesian models by defining the aforementioned likelihoods and priors. We then used R to access the output, evaluate the posterior densities and likelihoods to make inference on the parameters.

#### Model selection

Model selection is based on pairwise comparison of models using the Bayes Factors (BF). The BF for model *M* ^′^ relative to a simpler model *M* is is just the ratio of the models’ marginal likelihoods

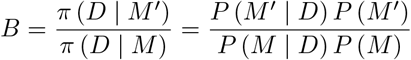

Assuming that the models are equally probable a priori, *i*.*e*., *P* (*M* ^′^) = *P* (*M*), then the BF can be seen as the posterior odds ratio. We use the criteria of Kass and Raftery [1995a] and chose the more complex model if it had at least substantial preference over the simpler model (*BF >* 3.2). We confirmed that this gave reasonable false positive rates on a single example; the false positive rate (preferring (BF *>* 3.2) the asymmetric model when the original data is generated from the symmetric/vanillla model) was evaluated for the asymmetric *v*_±_ model relative to the symmetric model, giving a false positive rate of around 1% on simulated data.

We compute the BF of all models relative to our base model eq. (1). Then, we determine which models have a substantial preference over the base model. If there are more than one models with substantial preference, and all of these models have the same complexity (*i*.*e*., equal number of parameters), we choose the model with the highest BF. If the set of models with substantial preference, contains models with different complexities and the more complex models (*i*.*e*., models with higher number of parameters), are nested to any of the other models in the set,then we compute the Bayes Factor of the complex-nested model relative to the simpler model and we choose the more complex model only when the preference of the complex model relative to simpler model is at least substantial and hence increasing the complexity is justified. Finally, if the set of models with substantial preference are models with different complexities but none of them is nested to another, then again we choose the model with the highest BF over the base model.

We use the “ bridgesampling” package in R, which computes the log-marginal likelihood for every model, using the log-likelihood samples passed from STAN HMC samples (which were in turn computed in the Forward Backward algorithm). This package utilizes the bridge sampling, [Meng and Wong, 1996, Gronau et al., 2020]. Then, the BF is just the ratio of the marginal likelihoods.

#### Convergence diagnostics

Convergence and mixing of MCMC chains is assessed via the Gelman-Rubin 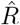 statistic, [Gelman and Rubin, 1992, Vehtari et al., 2021], using only results where 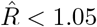 for all parameters.

Of *N* = 82 cells tracked, MCMC chains were run successfully for *N* = 59 cells, *i*.*e*., 26 and 33 DMSO and nocodazole washout. Where MCMC chains failed to run this was due either to poor tracking results in that cell (insufficient tracked kinetochore pairs or existence of too many missing data) or long time series such that the MCMC chains progressed extremely slowly (failed to find the typical set). Of the *n* = 1577 kinetochore pairs across 59 cells where MCMC chains ran successfully, MCMC chains from *n* = 27 (1.5%) kinetochore pairs failed to converge as assessed by the 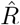 statistic, leaving estimates from *n* = 1550 kinetochore pairs.

Convergence proportions were slightly lower for the time dependent and metaphase-anaphase model, due to the increased complexity of these models.

### D Anaphase transition: modelling and analysis

#### D1 Modelling the metaphase-anaphase transition

To analyse anaphase further, we extended the asymmetric model (eq. (3)) to incorporate anaphase dynamics and an anaphase transition event, implemented similar to the metaphase-anaphase model in Sen et al. [2021]. Specifically, we introduce an additional hidden state, *A*, the anaphase state with dynamics given by,

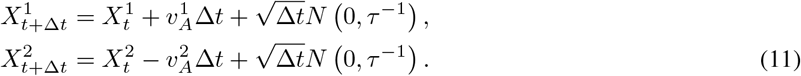

where 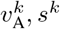, *s*^*k*^ are sister *k*’s velocity and diffusion coefficient in anaphase. Since the sisters are no longer connected the spring forces are removed, whilst the velocity in anaphase may be different to the depolymerising K-fibre velocity. We also removed the mid-plane centralising forces from the model since the PEF decays over anaphase, although the decay dynamics is unknown. Our model is only appropriate for the part of anaphase when the speed is approximately constant and the separation between the descending KT clusters is not too large, since as they approach the poles they slow down. We only fitted the model to linear regions (truncating movies if there were signs of slow down). Anaphase portions were around 50 frames, enough to infer the anaphase speed and anaphase onset time.

The transition to anaphase is modelled as a smooth switch with a transition probability to *A* at time *t* given by,

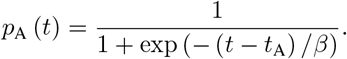

Thus, the transition to anaphase occurs around time *t*_A_ with *β* = Δ*t/*2 (fixed) determining the range over which switching can occur. We assume that the anaphase state *A* is equally accessible from each of the other states, but transitions back from anaphase to metaphase are not possible, *i*.*e*., state *A* is absorbing. Thus, once anaphase onset occurs the chromosomes segregate.

Switches between hidden states (see Figure 3B) occur at each time step according to the time dependent transition matrix (state order as eq. (7), with the anaphase state *A* in the last column/row),

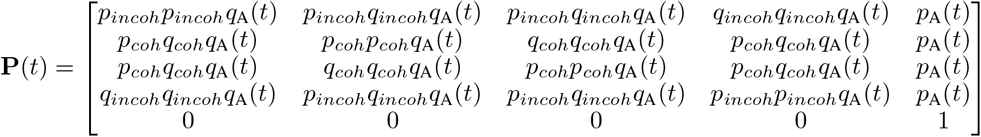

with *q*_*coh*_ = 1 − *p*_*coh*_ and *q*_*incoh*_ = 1 − *p*_*incoh*_ as before, and *q*_A_(*t*) = 1 − *p*_A_(*t*), where *p*_*coh*_ and *p*_*incoh*_ are the probabilities of a kinetochore remaining in the coherent, respectively incoherent state over a time interval Δ*t*, and *p*_*A*_ is the probability of transition to the anaphase state, *A*.

#### D2 Inference of anaphase parameters

We fitted the anaphase model to 790 KT pair trajectories (out of 798 pairs, 99.0%, 26 cells in DMSO) requiring at least 20 frames in anaphase. At anaphase the sister pairs separate, with 2 clusters of KTs approaching respective poles. These clusters separate at the same speed in DMSO and nocodazole washout treated cells, Appendix Figure D1A, although nocodazole washout treated cell clusters spread substantially compared to DMSO, that have approximately constant cluster width, Appendix Figure D1B. Although K-fibers are depolymerising in anaphase, we observed occasional reversals in individual KT trajectories, whereby a short reversal of direction occurs, Appendix Figure D1C. The inference algorithm for this metaphase-anaphase model displayed more divergences than our base model, rising to 3% and reducing the analysable KT-pairs to 767. In addition, as this model doesn’t incorporate events with transient anti-poleward KT movement in anaphase (reversals), which would reduce the inferred anaphase speed, we filtered out the KT pairs which had anaphase reversals by visual inspection. Hence, we report the fitting of the anaphase model to 611 KT pairs (of 767 pairs, 79.7%) in DMSO.

Inference of the anaphase model on nocodazole washout treated cells, (eq. (11)), had more divergences than in DMSO, *i*.*e*., 835 out of 927 pairs had no divergences, (90.1%) in nocodazole washout, compared to 767 out of 790 (97.1%) in DMSO, consistent with the higher frequency of reversals under nocodazole washout treated cells, Appendix Figure D1D. We filtered out KT pair trajectories with reversals leaving 456 KT pairs (from 30 cells) for which the inference is reliable. Anaphase dynamics in nocodazole washout treated cells was similar to DMSO based on the reversal free trajectories, Appendix Figure D5, with similar dispersion of anaphase onset times, and similar anaphase speeds.

An example of anaphase inference on an experimental trajectory is shown in Appendix Figure D3A, where both anaphase timing and the anaphase speed are inferred with high confidence and consistent with expert judgment. Within a cell we observe substantial variation in anaphase timing of KTs, Appendix Figure D3B, with an average standard deviation of 10.3 secs (range 5 secs to 32 secs) across the 26 cells. Anaphase timing varies with MPP location, Appendix Figure D3D, (*ρ* = − 0.27, *p*_*Corr*_ *<* 10^−14^), peripheral KT pairs entering anaphase earlier. The average anaphase pulling force was typically smaller than the average K-fiber pulling force, *i*.*e*., 67% of the pulling force on average. This is expected since the model ignores the centralising force in anaphase, and the K-fiber pulling force typically decreases towards anaphase whilst we are comparing *v*_*a*_ to the average metaphase force *v*_−_. There is a weak correlation between *v*_−_ and *v*_*a*_ (*ρ* = 0.17, *p*_*Corr*_ *<* 10^−15^), Appendix Figure D3F, although there is no trend in *v*_*a*_ across the MPP (Appendix Figure D3C), which is in stark contrast to the strong decrease in the strength of the pulling force with *r*. The only correlations of note are that the anaphase speed of sisters is correlated (*ρ* = 0.39, *p*_*Corr*_ *<* 10^−15^), Appendix

**Figure D1:**
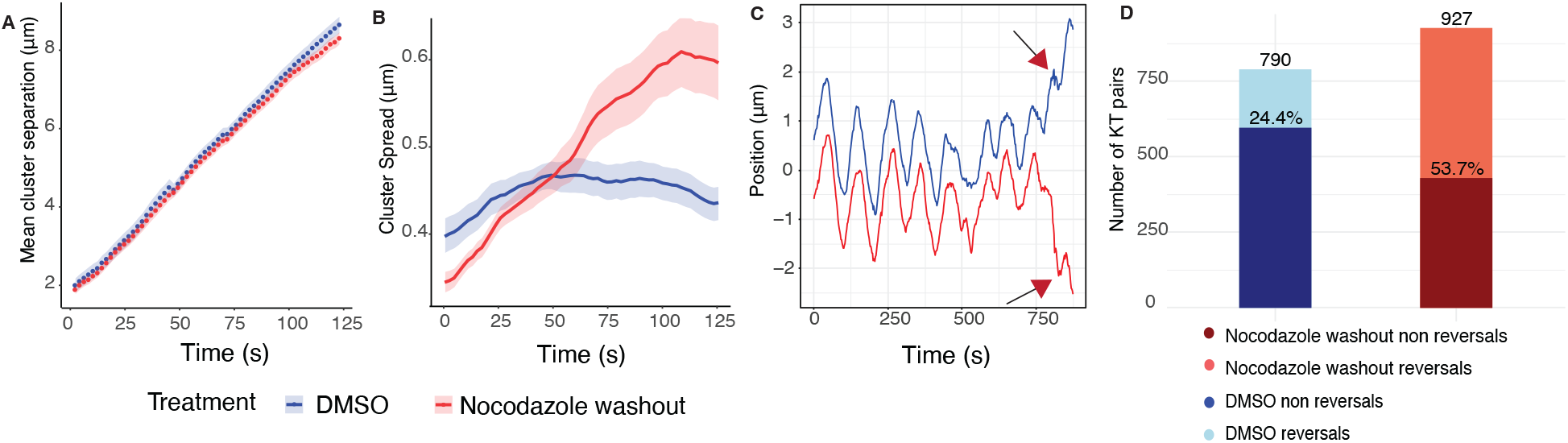
Anaphase clusters for DMSO (blue) and nocodazole washout (red) treated cells. Time *t* = 0 marks the median of estimated anaphase onset time per cell. **A** Mean cluster separation. Shaded areas denote the standard errors. **B** Cluster spreads during time and left axis based on 699 DMSO treated (blue) and 644 nocodazole washout treated (red) KT-pairs, which had long enough anaphase time points. **C** Example of a sister pair trajectory with reversals in anaphase. Arrows indicate reversal events. **D** Summary of KT-pairs with reversals and no reversals in DMSO (blue) and nocodazole washout cells (red). From 790 (927) KT-pairs 611 (465) pairs had no reversals, *i*.*e*., 77.3% (49.2%) for DMSO (nocodazole washout treatment).

Figure D3H, and there is positive correlation (*ρ* = 0.247, *p*_*Corr*_ *<* 10^−9^) between anaphase speed and anaphase time relative to median anaphase time, *i*.*e*., sisters with a later transition to anaphase have a higher anaphase speed. Similar results were obtained if reversals were not filtered out.

The onset of anaphase, when kinetochore pairs begin to separate and segregate towards their respective spindle poles is tightly controlled temporally, [Holt et al., 2008], and appears entirely synchronous at low time resolution, but is in fact asynchronous, Appendix Figure D3B, (previously reported for RPE1 cells in Armond et al. [2019], Sen et al. [2021]). Peripheral sister-chromatids initiate anaphase earlier than central sister-chromatids by 4s in RPE1 cells, and have a lower anaphase speed, Appendix Figure D3C,D (there is positive correlation (*ρ* = 0.247, *p*_*Corr*_ *<* 10^−9^) between anaphase speed and relative anaphase time). However, sister pairs with stronger pulling asymmetry are not biased towards having a higher anaphase speed, Appendix Figure D4. Further, we observed a weak correlation between the pulling speed 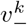 and 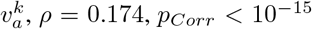, *ρ* = 0.174, *p*_*Corr*_ *<* 10^−15^, Appendix Figure D3F, thus although our estimated anaphase force (our model is ignoring PEF in anaphase) is the same order as the K-fiber pulling forces in metaphase, there is only a weak correlation between K-fiber pulling in metaphase and anaphase, whilst spatial trends in the MPP are distinctly different.

This likely reflects the fact that K-fibers in metaphase and anaphase are regulated by different mechanisms and the KTs are in different states, although it is also reminiscent of the anaphase speed governor, [Anjur-Dietrich et al., 2021]. K-fibers in metaphase undergo dynamic instability and rarely have high coherence between MTs in the bundle, [Armond et al., 2015b], whilst anaphase K-fibers depolymerise for extended periods and high coherence is expected. However, we do observe a fraction of KTs undergoing transient reversals of the usual poleward motion during anaphase, Appendix Figure D1. These reversals were reported in previous studies, [Skibbens et al., 1993], whilst metaphase-like chromosome oscillations have been shown to persist into anaphase upon inhibition of protein dephosphorylation, [Su et al., 2016]. The cause of these reversals is not understood, but suggests that coherence of microtubules within anaphase K-fibers may be dependent on factors that are perturbed under nocodazole washout, potentially the efficacy of dephosphorylation.

### E Additional Models

#### Changepoint model

A simple changepoint model [Armond et al., 2019] to assess the time of anaphase assumes that the intersister distance between a kinetochore pair is constant in metaphase, and increases linearly in time during anaphase. If *d*_*t*_ is the 1D intersister distance (in the *x* direction perpendicular to the metaphase plate) between kinetochore sisters, then at time point *t*_*i*_

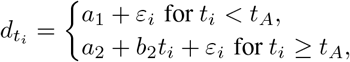

with the condition that *a*_1_ = *a*_2_ + *b*_2_*t*_*A*_. Since this model is simpler than the biophysical model, a uniform prior can be used for *t*_*A*_ and weakly informative priors for other parameters.

**Figure D2:**
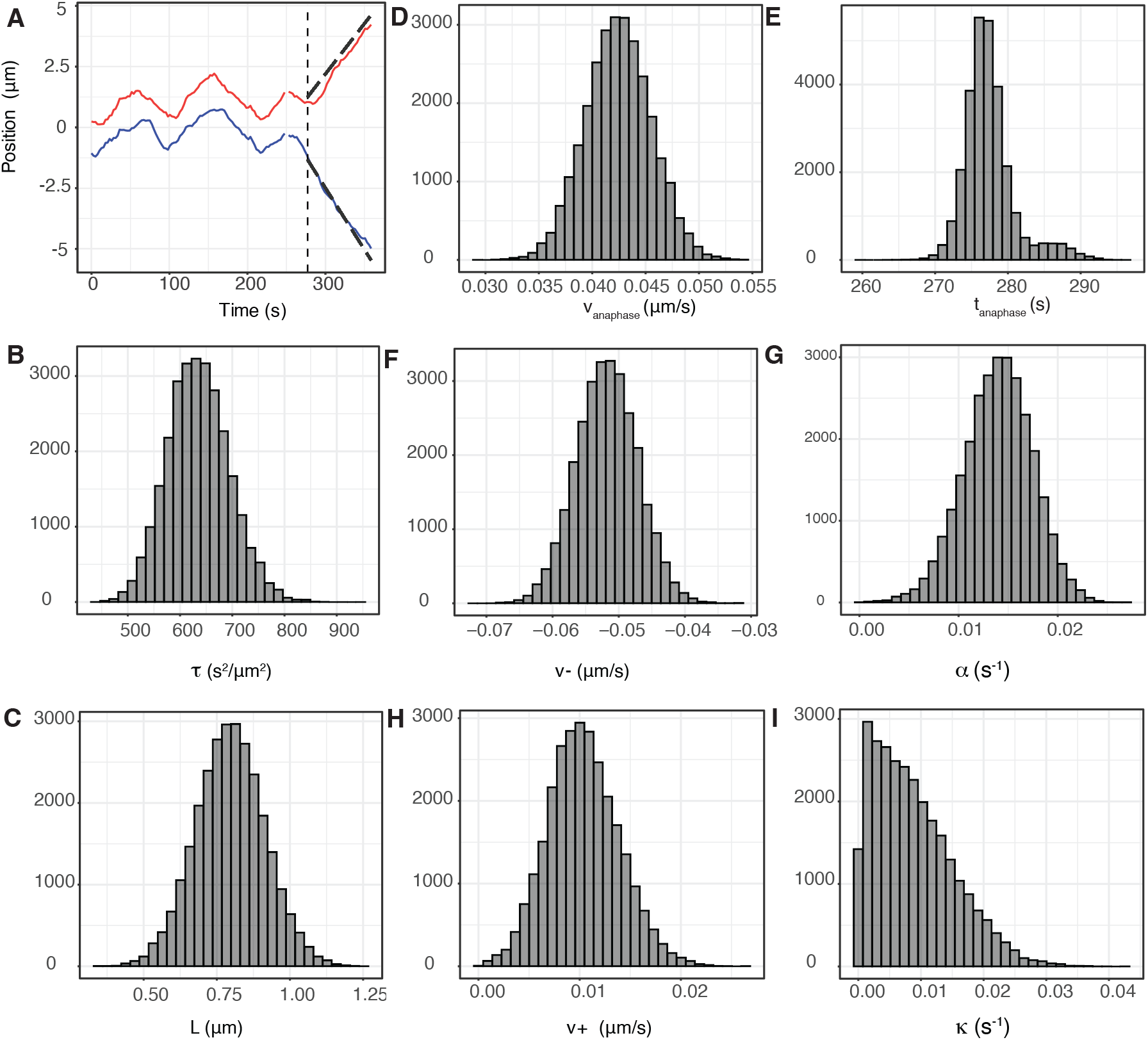
Single trajectory metaphase-anaphase model inference showing parameter estimates and trajectory annotation (DMSO treated cell). **A** Observed KT sister pair trajectory. **B-I** Marginal posterior distribution of the biophysical parameters for the trajectory data in A using the metaphase-anaphase model. Note: *v* _−_, *v*_+_ and *v*_*anaphase*_ are averaged over the two sisters.

**Figure D3:**
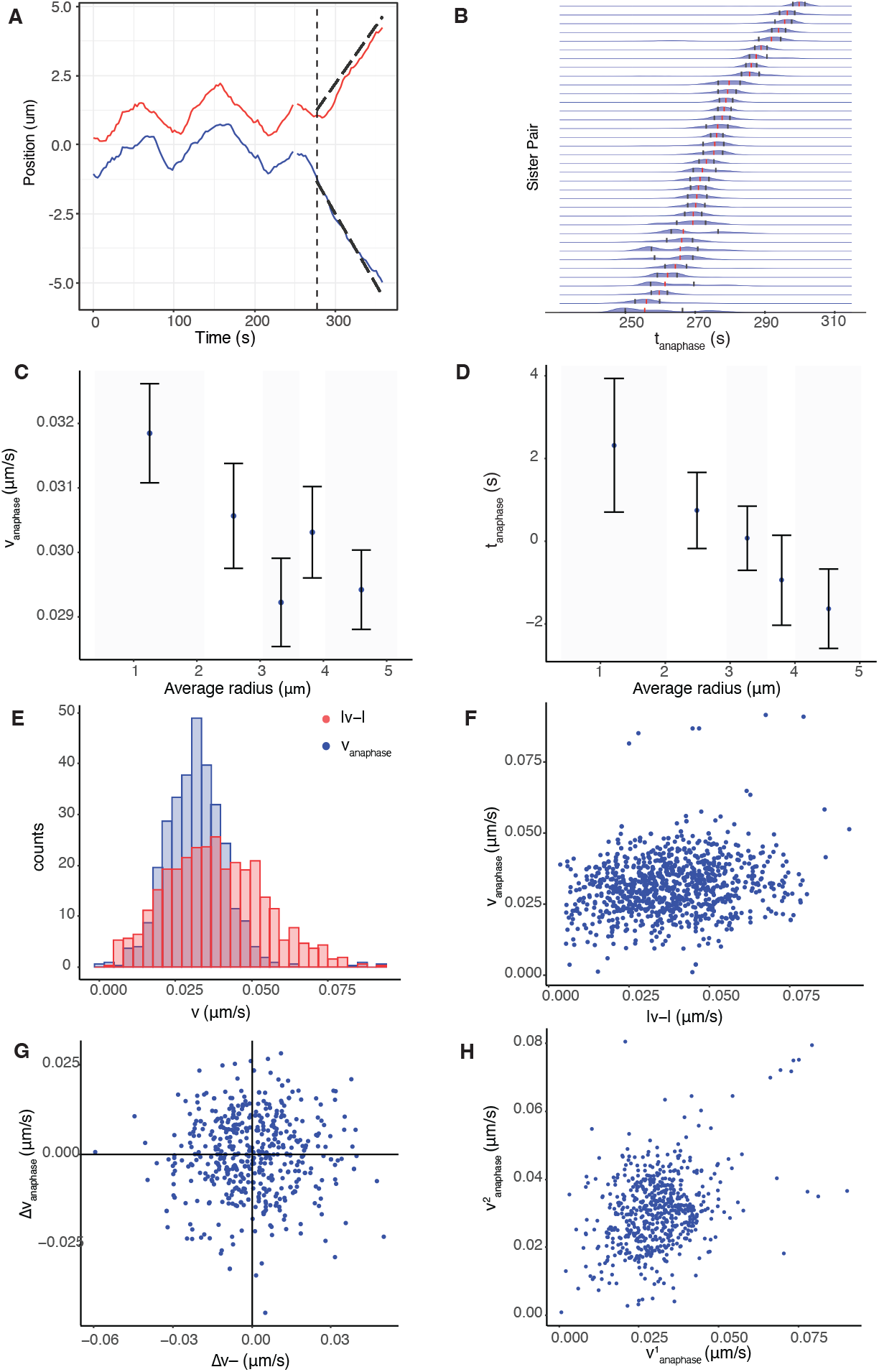
Anaphase K-fibers are dynamically homogeneous across the MPP despite variation in anaphase initiation (DMSO). **A** Example of the anaphase inference on a KT sister pair trajectory with anaphase event indicated with vertical line, and dashed lines above/below anaphase trajectory have gradients of 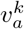. **B** Anaphase timing of KT pairs in an individual cell. Red dots denote the distribution median and blue dots define the interquartile range. **C/D** Spatial biophysical parameter trends across the metaphase plate for anaphase speed/anaphase timings (relative to the median anaphase time per cell). The radial position of the KT pairs refers to their position during metaphase. The correlation between anaphase speed and anaphase timings is 0.247 (*p*_*Corr*_ *<* 10^−9^). **E** Estimated densities of *v*_*anaphase*_ (blue) and pulling force (red), (normalised by the drag coefficient). **F** Correlation of anaphase speed and absolute *v*_−_ (correlation = 0.174, *p*_*MW*_ *<* 10^−15^). **G** Difference of 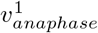 and 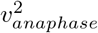 versus 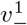 and 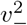only on asymmetric (on *v*_−_) sister pairs. No statistically significant correlation is observed. **H** Correlation of anaphase speeds for sister 1 and sister 2 (correlation = 0.386, *p*_*Corr*_ *<* 10^−15^). Figures are based on a subset of KT pair trajectories (*N* = 611 from 26 cells cultured in DMSO), which exhibited no reversals during anaphase (visual inspection), had adequate coverage of anaphase.

**Figure D4:**
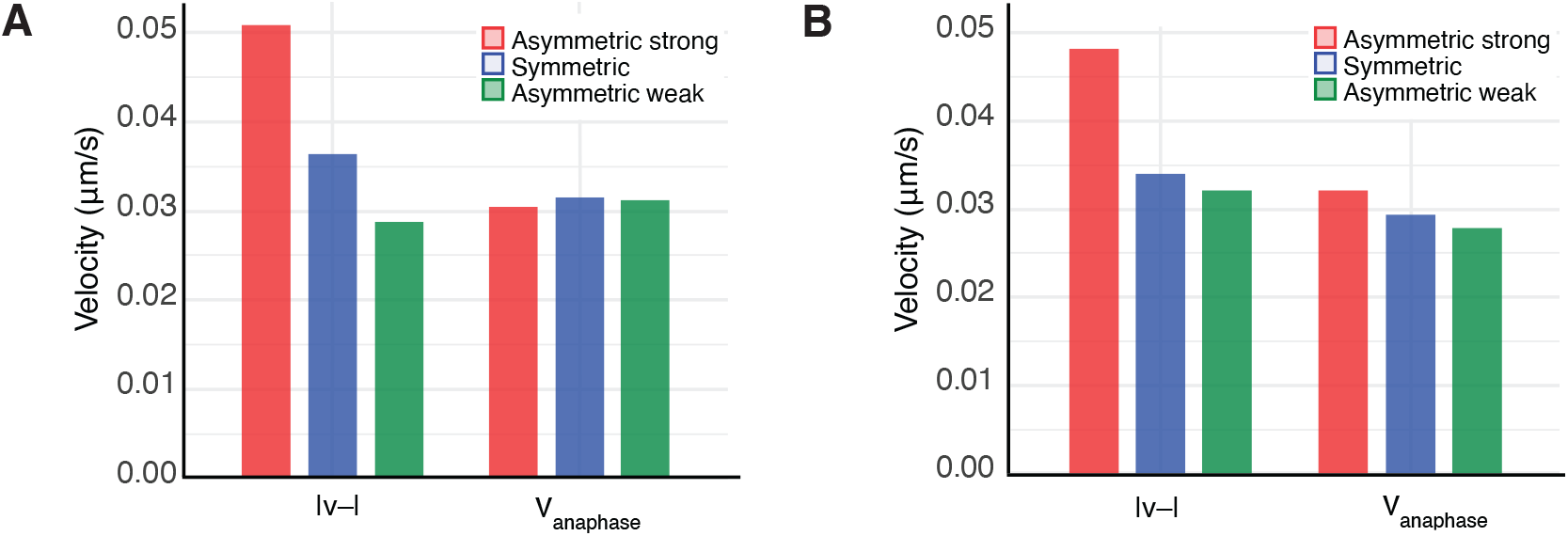
Comparison of averaged median speeds *v*_−_ and *v*_*anaphase*_ for pulling strength groups on **A** DMSO (*N* = 611) and **B** nocodazole washout (*N* = 767) treated cells, (restricted to KT pairs with no reversals).

#### Modelling joint pulling/pushing strength of K-fibers: a multivariate Normal mixture model analysis

We addressed if there is evidence of s specific asymmetric subpopulation. The 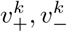 parameters are not independent (testing for independence on the full asymmetric model, using all KTs: *p*_*Hoef*_ *<* 10^−18^, *p*_*DC*_ *<* 10^−4^, *p*_*Corr*_ *<* 10^−5^), with a significant correlation between pushing (*v*_+_) and pulling (*v*_−_) forces, Figure 5D. Therefore we modelled the joint empirical distribution of *v*_−_, *v*_+_ by fitting a mixture of multivariate Gaussians.

The multivariate Normal (Gaussian) model is a generalisation of the univariate normal model

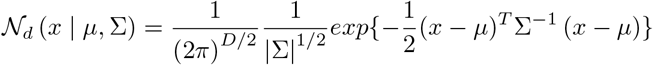

with *µ* ∈ *ℛ* ^*d*^, the location vector and Σ ∈*ℛ* ^*d*×*d*^, the covariance matrix. We assume that *v*_−_ and *v*_+_ follow a multivariate normal distribution (allowing for *v*_−_ and *v*_+_ to be dependent). Moving a step further, we have found that the model which explains the 2-dimensional distribution of pulling and pushing forces, is a 3-component multivariate normal mixture model which is defined as a linear superposition of the aforementioned multivariate normal model, *i*.*e*., *v*_−_ and *v*_+_ have the following distribution,

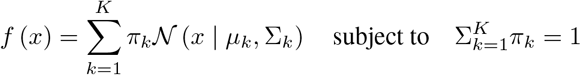

with *k* = 3, the number of Gaussian components, *π*_1_…*π*_*k*_, the mixture weights of the components, *µ* = *µ*_1_…*µ*_*k*_, a vector containing the mean of each component and Σ_1_…Σ_*k*_, the covariance matrices of the components.

This model fitted the data extremely well, the QQ plot in fact indicating that there are no extremal subpopulations with large, or small, | *v* _−_|, Figure E1B. Hence, this suggests that sister asymmetry is caused by natural variation of the K-fibres within a cell.

**Figure D5:**
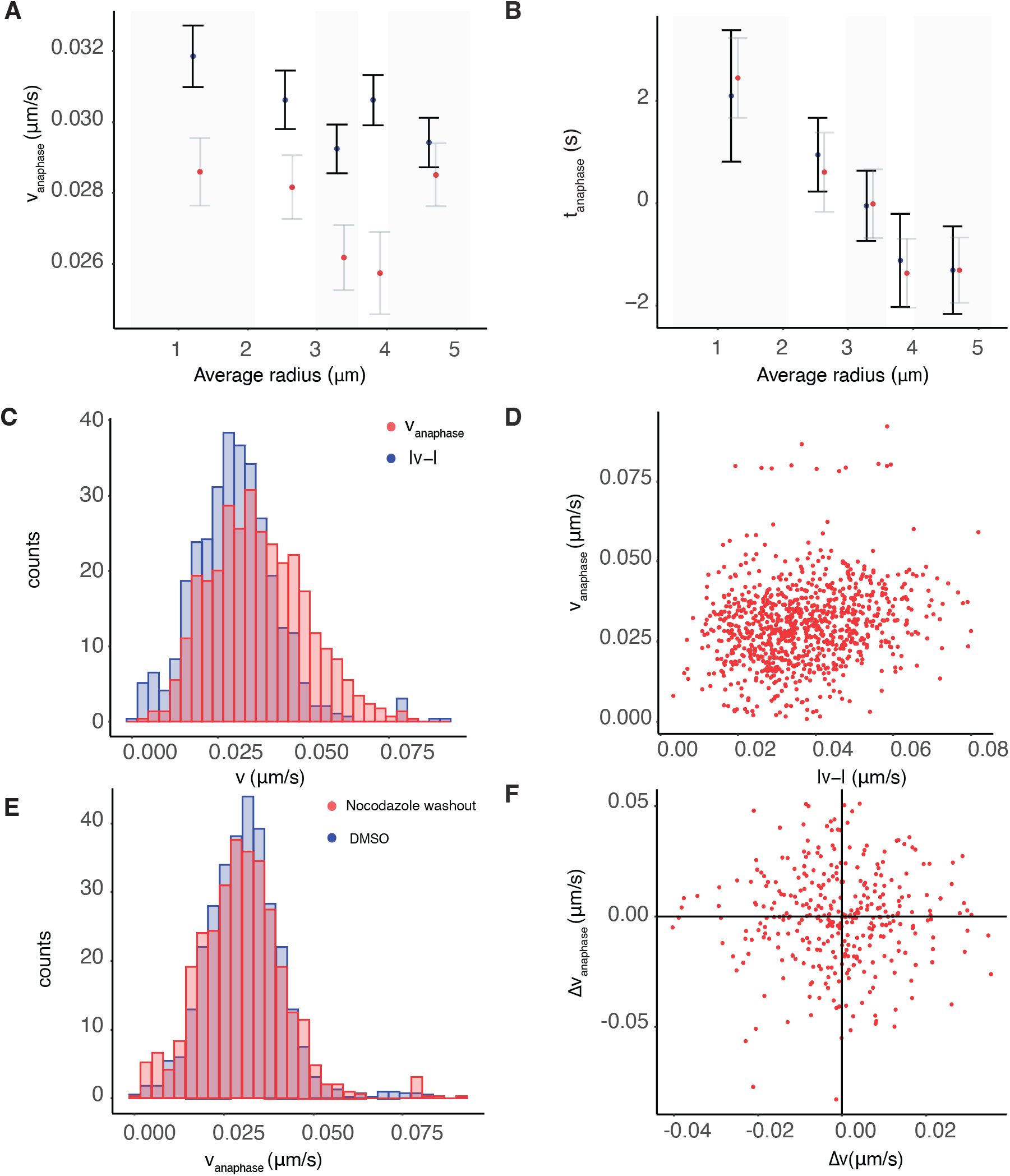
Homogeneity of anaphase K-fibers in DMSO cultured and nocodazole washout cells. **A/B** Spatial biophysical parameter trends across the metaphase plate for anaphase speed/anaphase timings for DMSO (blue) and nocodazole washout (red) treated cells. The estimated correlation in nocodazole washout treated cells between anaphase speed and relative anaphase timings is 0.125 (*p*_*Corr*_ *<* 10^−4^). **C:F** Analysis based on nocodazole washout treated cells. **C** Estimated densities of *v*_*anaphase*_ (red) and pulling force (blue) for nocodazole washout treated cells. **D** Correlation of anaphase speed and absolute *v*_−_ (estimated correlation = 0.174, *p*_*Corr*_ *<* 10^−15^). **E** *v*_*anaphase*_ for DMSO (blue) and nocodazole washout treated cells. Distributions are not statistically different (*p*_*MW*_ = 0.165), with the variance of *v*_*anaphase*_ nocodazole washout pairs being greater (*p*_*F* *test*_ *<* 10^−4^). **F** Difference of 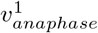 and 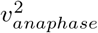 versus 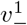 and 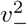 only on asymmetric (on *v*_−_) sister pairs, for nocodazole washout treated cells. No statistically significant correlation is observed. Inference is based on 456 pairs across 30 nocodazole treated cells.

**Figure D6:**
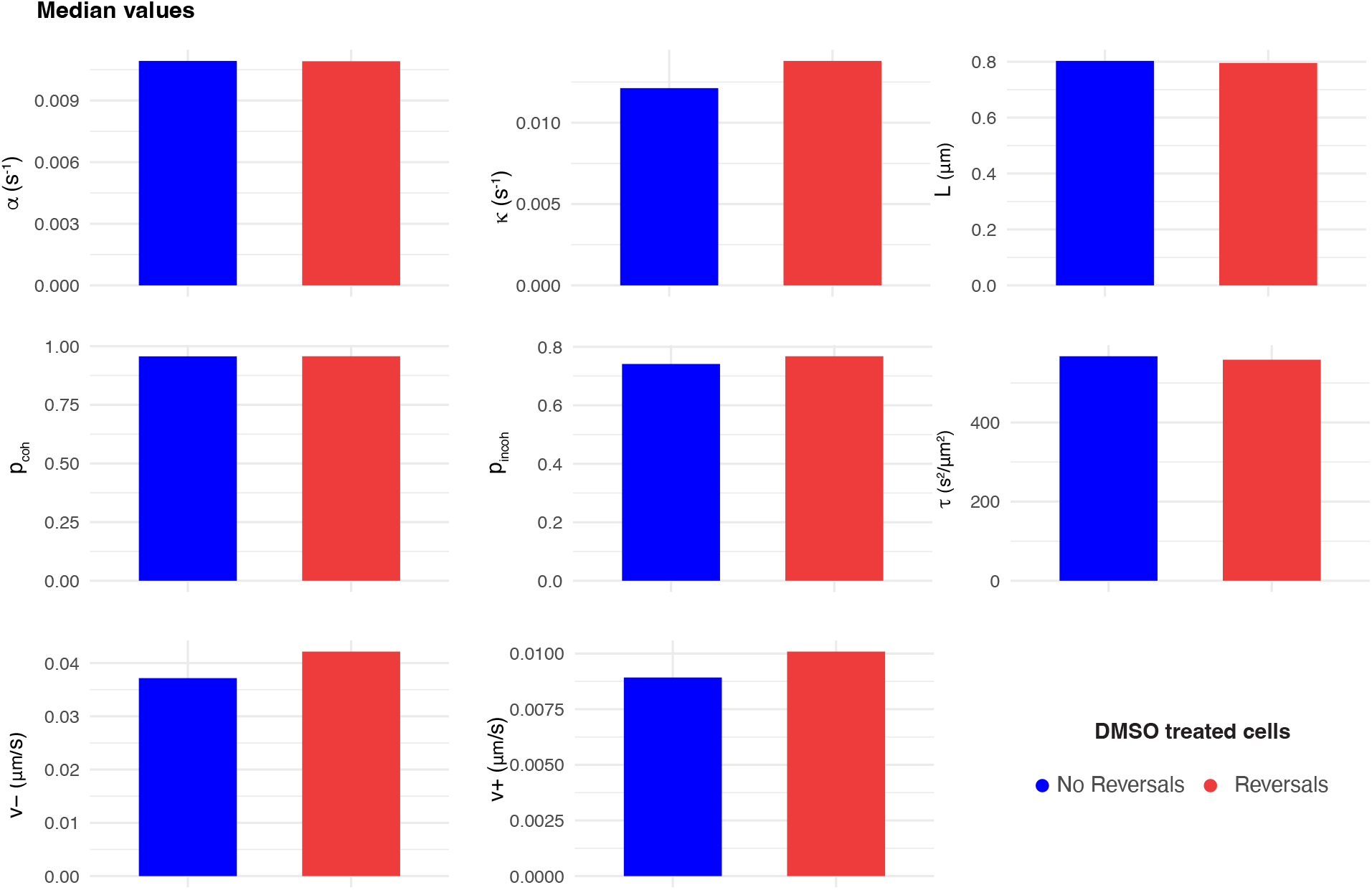
Bar plots of the median of the posterior median estimates of the trajectories with reversal events (red) and no reversal events (blue) during anaphase for DMSO treated cells. As shown in Table A12, the distributions of *v*_−_, *v*_+_ and *L* are significantly different.

**Figure D7:**
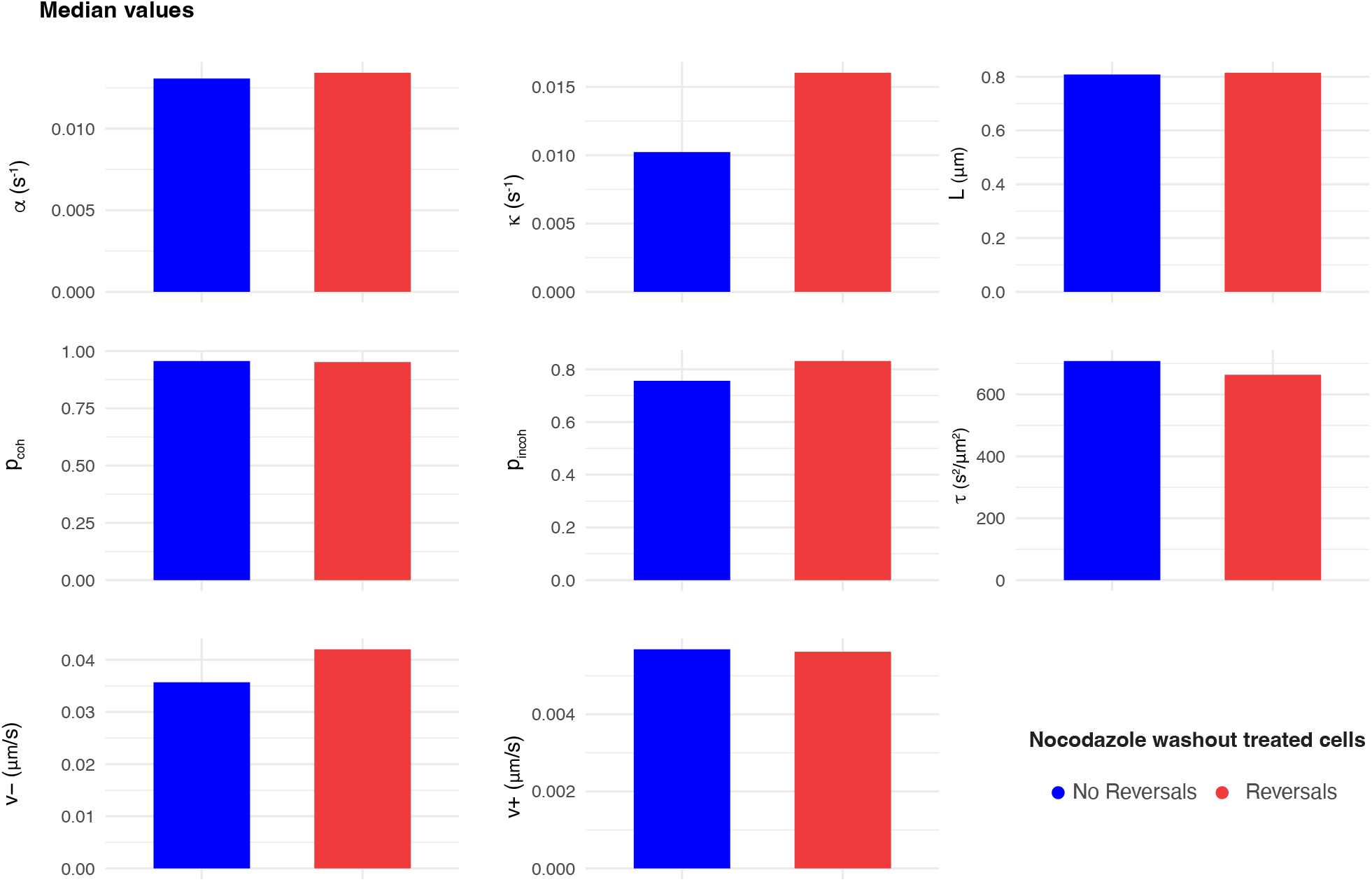
Bar plots of the median of the posterior median estimates of the trajectories with reversal events (red) and no reversal events (blue) during anaphase for nocodazole washout treated cells. As shown in Table A12, the distributions of all parameters but *v*_+_ and *α* are significantly different.

**Figure E1:**
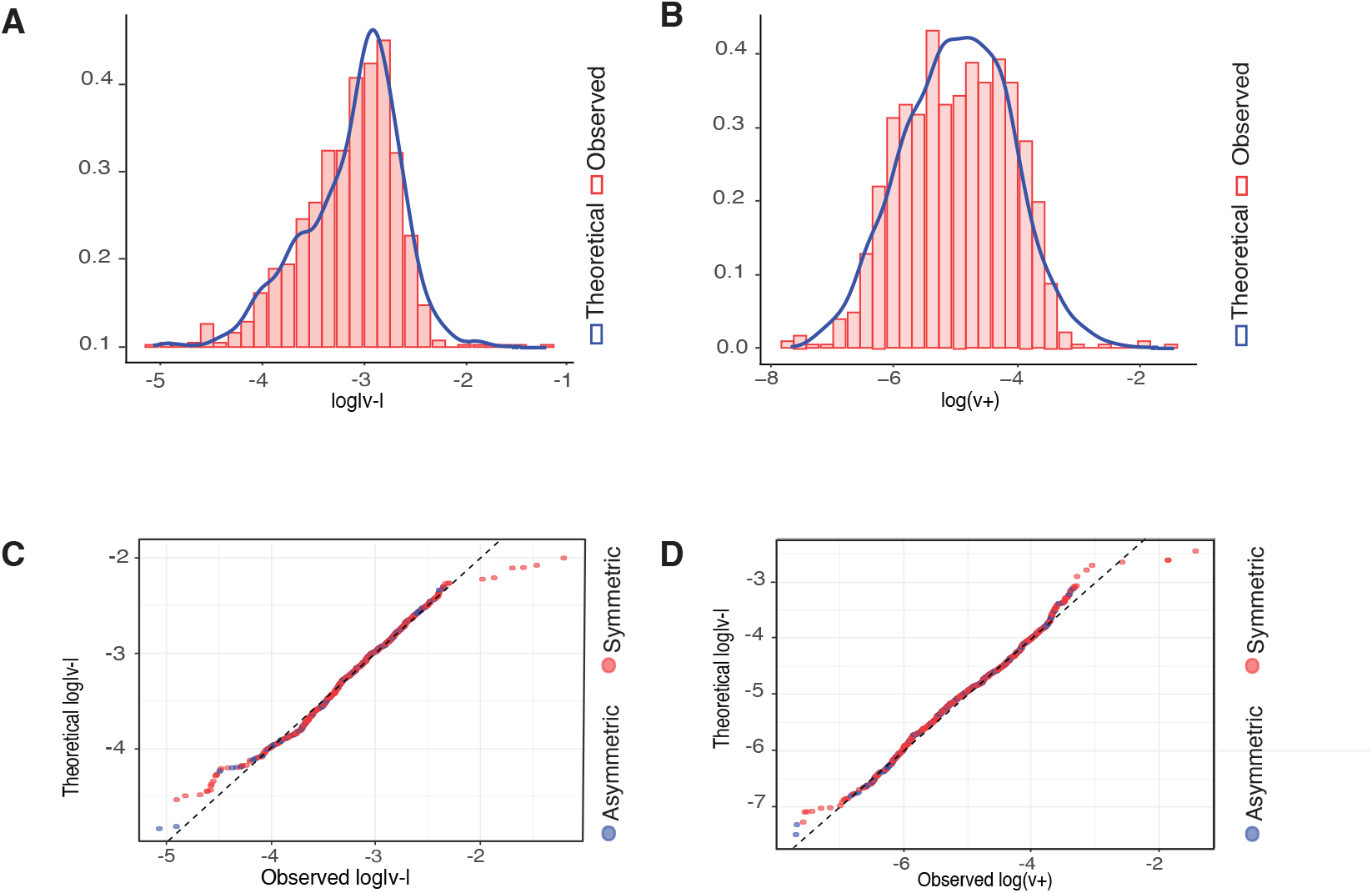
Mixture model of *v*_±_ joint distribution. **A**,**B** Histograms of the estimated (red) and theoretical (blue) **A** log | *v*_−_| and **B** log (*v*_+_) of 3 − component multivariate normal mixture model. **C**,**D** QQ plots for 3-component multivariate mixture model showing excellent fit.

## Footnotes

1 Details and abbreviations on the statistical tests used in throughout our analysis can be found in Appendix Table C3.

